# Coupling of habitat-preference barriers leads to reproductive isolation in sympatric speciation

**DOI:** 10.1101/2025.07.03.662681

**Authors:** Zakia Sultana, Curtis Heyda, Karyn How, John Lin

## Abstract

Habitat preference is a widely recognized mechanism of reproductive isolation, yet its role in initiating premating barriers and coupling with other barrier mechanisms to establish robust and irreversible reproductive isolation (RI) in sympatric speciation remains unclear. In this study, we developed mathematical models and computer applications to investigate one- and two-allele models of habitat-preference barriers in sympatric populations under disruptive ecological selection. We examined two spatial arrangements: an open-space model inspired by sympatric cichlid fishes that meet in open water with niche habitats that are small relative to lake size, and a no-open-space model inspired by sympatric hawthorn and apple maggot flies that move directly between trees without lingering in midair. Next, we examined coupling between habitat-preference barriers and a two-allele mating-bias barrier developed in a prior study to analyze how their invasion and coupling dynamics could lead to stronger RI. Our findings confirm that habitat preference is an independent mechanism capable of establishing initial premating barriers in sympatric speciation. Moreover, it can couple with additional barriers, such as those based on mating-trait discrimination, to enhance overall RI. Because habitat preference is an adaptive barrier mechanism, its invasion and coupling are driven by selection pressures arising from maladaptive hybrid loss, and it is readily reversed when disruptive ecological selection weakens. The one-allele model is easier to evolve than the two-allele model because it is immune to recombination by gene flow. Habitat-preference barriers can facilitate the emergence of mating-bias barriers. Open-space systems provide fewer opportunities for inter-niche encounters and tend to result in stronger RI compared to no-open-space systems. By elucidating the habitat-preference mechanism, our study reinforces the important role of habitat-preference barriers in sympatric speciation. It also provides insights into a wide range of premating isolating mechanisms—temporal, behavioral, and mechanical—that function similarly by reducing mating encounters.

## I. INTRODUCTION

**I**n evolutionary biology, understanding how initial reproductive isolation (RI) arises in sympatric speciation remains a challenge [1, 2]. The difficulty stems from the presence of gene flow, which inhibits the nonrandom assortment of adaptive ecological traits and sexual discrimination traits between specialized niche ecotypes. Intuitively, when separate and distinct niche habitats exist, this makes habitat preference an ideal initial barrier mechanism to counteract the homogenizing effect of gene flow and jumpstart the speciation process. In sympatric populations under disruptive ecological selection, maladaptive hybrids are produced through inter-niche mating between locally adapted ecotypes. A mutation that promotes habitat preference—causing individuals to remain within their adaptive niches—can reduce hybrid offspring loss, thereby conferring a fitness advantage to ecotypes carrying the mutation over their same-niche counterparts lacking it. This fitness advantage allows the habitat-preference mutation to invade and spread, ultimately reducing gene flow and establishing premating RI between niche ecotypes [3-5].

However, just because two locally adaptive niche populations do not encounter each other to mate does not, by itself, render them reproductively incompatible or qualify them as distinct species under the biological species concept [6]. For long-term reproductive isolation to persist—especially if encounter barriers are later removed—some form of sexual discrimination or incompatibility is ultimately required. In nature, most established sympatric sister species possess mating-bias barriers as part of their premating isolation. Moreover, when distinct habitats exist to support the evolution of habitat preference, both habitat-preference and mating-bias barriers are often observed together [7-11].

Reproductive barriers can be classified as either a one-allele model or a two-allele model [12, 13]. In a two-niche ecosystem, a one-allele model requires only a single mutation that can invade and spread across both niches to establish RI. In contrast, a two-allele model requires different mutations to arise and persist in each niche. Its barrier effects depend on the nonrandom assortment of mutant alleles between niches—a condition that is more difficult to achieve in the presence of gene flow. As a result, a one-allele model is generally easier to establish, as it is unaffected by the homogenizing effects of gene flow [1].

Many researchers have investigated the one-allele and two-allele models of habitat-preference barriers and their potential to couple with sexual discrimination barriers [7, 14-19]. Most studies employ computer simulations based on mutation-driven models to explore the invasion and coupling dynamics of these barriers in a two-island or a continent–island spatial arrangement, where interactions between habitat populations are governed by a migration parameter, *m* [7, 14, 15, 20].

In our study, we used a mathematical model that is more analytical and less dependent on simulation-based approximations to investigate barrier dynamics in a purely sympatric environment, where individuals can migrate and intermix freely in the absence of habitat preferences. We modeled both one- and two-allele habitat-preference barriers. For sexual isolation, we employed a two-allele mating-bias model developed in a prior study [21]. In this model, niche ecotypes evolve distinct mating traits and preferences—such as two different color morphs of fish—that determine matching success during mating encounters. We selected the two-allele model because we believe it more closely reflects how sexual cues are commonly observed between incipient sister species to prevent interbreeding [22]. We did not examine the one-allele model of mating-bias barriers (e.g., preferences based on size or morphology within conspecifics) [12, 14, 23-27], as it is theoretically easier to evolve than the two-allele model—an outcome confirmed by previous studies [7, 15, 28, 29].

We investigated habitat-preference barriers under two spatial arrangements. The first is an open-space system, inspired by sympatric cichlid fishes [30, 31], in which niche ecotypes do not visit each other’s habitats and inter-niche encounters occur only in a communal open space. The second is a no-open-space system, based on sympatric hawthorn and apple maggot flies [32-34], where ecotypes can visit one another’s habitats, and all encounters take place only within those habitats.

User-friendly MATLAB applications were developed to solve the different barrier models and display the results as phase portraits. These visualizations help clarify the properties of each barrier mechanism, as well as their invasion and coupling dynamics. Using this approach, we aim to understand how habitat-preference barriers can independently initiate premating RI in the early stages of sympatric speciation, how they may couple with mating-bias barriers to establish sexual isolation, and how the order of coupling influences the final outcome and interactions among these early-stage premating barriers.

Broadly, our study reveals the following key findings: (1) Habitat preference is an adaptive barrier mechanism, with its invasion driven by selection against maladaptive hybrid loss; (2) as long as hybrid loss persists, a habitat-preference barrier can always invade and couple to reduce that loss, to the extent permitted by its preference biases; (3) one-allele models are easier to evolve and produce stronger RI than two-allele models; (4) habitat-preference barriers evolve more readily and result in stronger RI in open-space systems than in no-open-space systems; (5) when habitat preference evolves first, it can facilitate the coupling of a mating-bias barrier; and (6) very low gene flow can hinder the coupling of a mating-bias barrier by generating an invasion-resistant pattern in its phase portrait—a constraint that may be mitigated by increasing the number of mating rounds, reducing niche size, or strengthening selection against hybrids.

Both habitat preference and mating bias are important premating barrier mechanisms capable of independently establishing initial RI in the early stages of sympatric speciation [35]. Our study clarifies the properties of habitat-preference barriers under two spatial configurations and their coupling dynamics with a two-allele mating-bias barrier. Since habitat preference is easier to evolve as an initial barrier than the more finicky two-allele mating-bias barrier—and can facilitate the subsequent coupling of a mating-bias barrier—a “habitat preference first, mating bias second” sequence may offer a more realistic pathway to establishing initial premating RI in sympatric speciation, whenever distinct habitats exist to support the evolution of habitat preferences [7, 15].

Notably, distinct habitats do not need to be spatially separated for habitat-preference barriers to evolve. Separation can occur along various structural dimensions—spatial, temporal, mechanical, or behavioral [36-39]. As a result, the general models presented in this study are broadly applicable to a wide range of reproductive barrier mechanisms that generate RI by reducing mating encounters, rather than through sexual discrimination during those encounters.

## II. METHODOLOGY

Our study used MATLAB (version R2021a) and its App Designer tool to develop user-friendly graphical user interface (GUI) applications that solve mathematical models of habitat-preference barriers and their coupling with a two-allele mating-bias barrier from a previous study [21]. The results are displayed as phase portraits to provide a bird’s-eye view of the solution landscape and an intuitive understanding of the underlying dynamics.

### 1. Mathematical model of a two-allele mating bias barrier

For the mating-bias barrier, we use the 2-mating-bias-allele, 2-ecological-niche mathematical model without viable hybrids that was developed in our previous study (see Fig M3) [21].

The flowchart in Fig M1 shows the life cycle of a sympatric population used in our simulations. In each generation, individuals undergo ecological and sexual selection to produce offspring for the next generation. The ecosystem contains two distinct niche resources, *A* and *B*, generated by disruptive ecological selection. During the ecological selection phase, only individuals with genotypes adapted to one of the two niches (i.e., ecotype *A* or ecotype *B*) can access the niche-specific resources and survive.

Next, in the sexual selection phase (see flowchart in Fig M2), survivors of ecological selection encounter one another randomly to find mates. The outcome of each matching encounter is determined by the mating-bias alleles—*X* or *Y*—that individuals carry at a single gene locus. Compatibility between alleles is defined by the matching compatibility table shown in Fig M4. Individuals with identical mating-bias alleles always match (i.e., with a probability of 1), while individuals with different alleles match with a probability defined by the mating-bias value *α*, which ranges from 0 to 1.

Unmatched individuals may attempt to find a mate again in subsequent rounds, up to a maximum of *n* rounds. Those who fail to match after *n* rounds die without producing offspring, while matched pairs reproduce offspring through random assortment of their alleles at each gene locus. Thus, the number of matching rounds, *n*, reflects the cost of assortative mating, and the values of *α* and *n* together determine the strength of sexual selection.

Fig M3 shows the two-mating-bias-allele, two-niche mathematical model used to describe the population dynamics of the sympatric population during a matching round. The model includes four genotype groups—*NAx, NAy, NBx*, and *NBy*— representing the normalized population ratios of the different ecotypes and their associated mating-bias alleles. Because all parametric values in the model are normalized, *NAx* + *NAy* + *NBx* + *NBy* = 1.

As these genotype groups encounter one another randomly in sympatry, their pairwise encounter probabilities are summarized in the Encounter Probability Matrix shown in Fig M5. The elements of this matrix represent all possible pairings and collectively sum to 1.

After *n* matching rounds, all successfully matched genotype pairs are collected. These parental pairs reproduce by randomly assorting alleles at each gene locus. Fig M6 presents a Unit Offspring Matrix that lists the expected genotype ratios of offspring produced by different parental genotype combinations. In this matrix, the variable *f* denotes the “offspring return ratio,” which specifies the proportion of offspring that inherit the same genotype as their parents. The value of *f* ranges from 0 to 0.5 and serves as a measure of the strength of ecological selection.

The unit offspring ratios represent the normalized proportions of different offspring genotypes produced by each generation of matched parental pairs, under the assumption that each parent produces only one offspring to replace itself. The values in each cell of the Unit Offspring Matrix sum to 1. Because the ratios of all parental pairs are normalized (i.e., they sum to 1), the total offspring genotype ratios produced by the parental pairs also sum to 1. To compute the final offspring population sizes, the unit offspring ratios are multiplied by a fertility parameter, *off*, which specifies the actual number of offspring each parent produces.

Although the model assumes no viable hybrid offspring by default, scenarios involving viable hybrids supported by hybrid niche resources can still be represented by increasing the value of *f*. Raising the value of *f* decreases the effective strength of ecological selection.

To avoid the effect of “incumbent selection”—a phenomenon in which a dominant population can use its numerical advantage to eliminate a minority population through their attritive interactions—our mathematical model assumes that ecotypes in each niche reproduce sufficient offspring to fully occupy the niche’s carrying capacity after each mating generation [21]. Therefore, in our model, the normalized niche population ratios, *NA* and *NB*, are always fixed at the beginning of the sexual selection phase in each generation.

We then use computer applications to generate *Ax*/*Bx* phase portrait solutions of the model using user-defined parameter values. In the phase portrait, *Ax* represents the normalized ratio of niche-*A* ecotypes carrying the *X* allele, while *Ay* denotes those with the *Y* allele, such that *Ax* + *Ay* = 1. The same definitions apply to *Bx* and *By* for niche *B*. The phase portrait displays changes in these normalized genotype ratios over *g* generations.

We classify a barrier’s phase portrait as either “convergent,” in which the vector field leads (converges) to fixed points, or “divergent,” where no fixed points are present. Assuming that *NAx* represents the largest genotype group in the model, maximum RI is achieved when *NAx* and *NBy* are large, while *NAy* and *NBx* are small. In the *Ax*/*Bx* phase portrait, this outcome corresponds to a fixed point located as close as possible to the bottom right corner, where *Ax* = 1 and *Bx* = 0.

**Fig M1.**
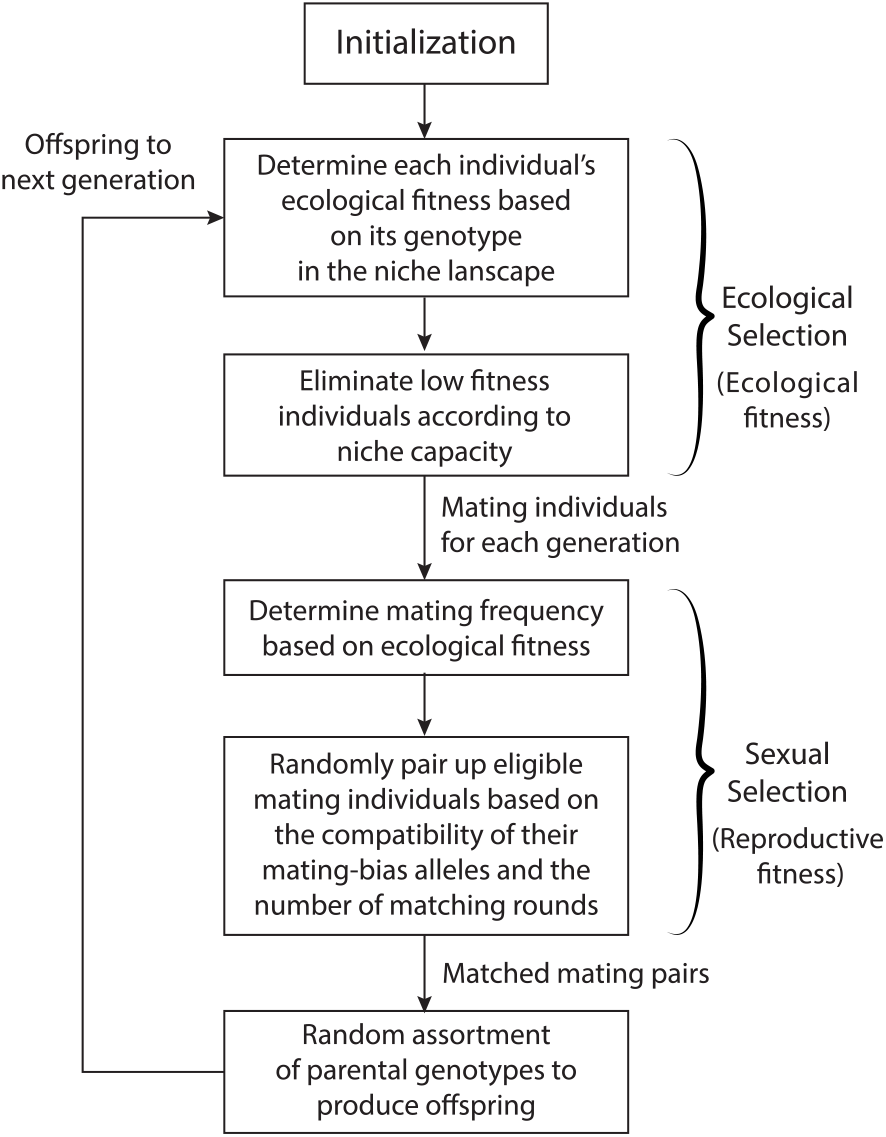
A flowchart showing the life cycle of a sympatric population used in the computer simulations.

**Fig M2.**
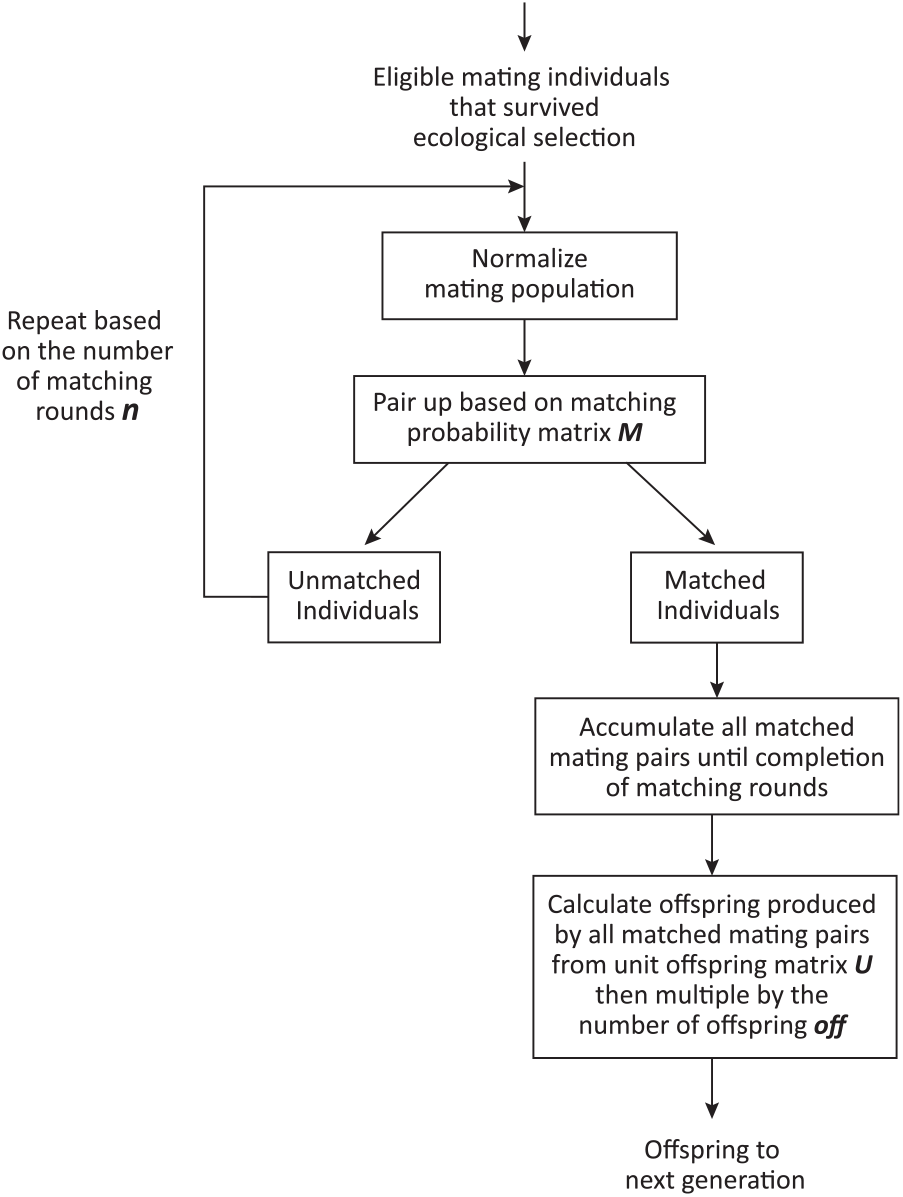
A flowchart showing the algorithm used to match eligible mating individuals and reproduce offspring.

**Fig M3.**
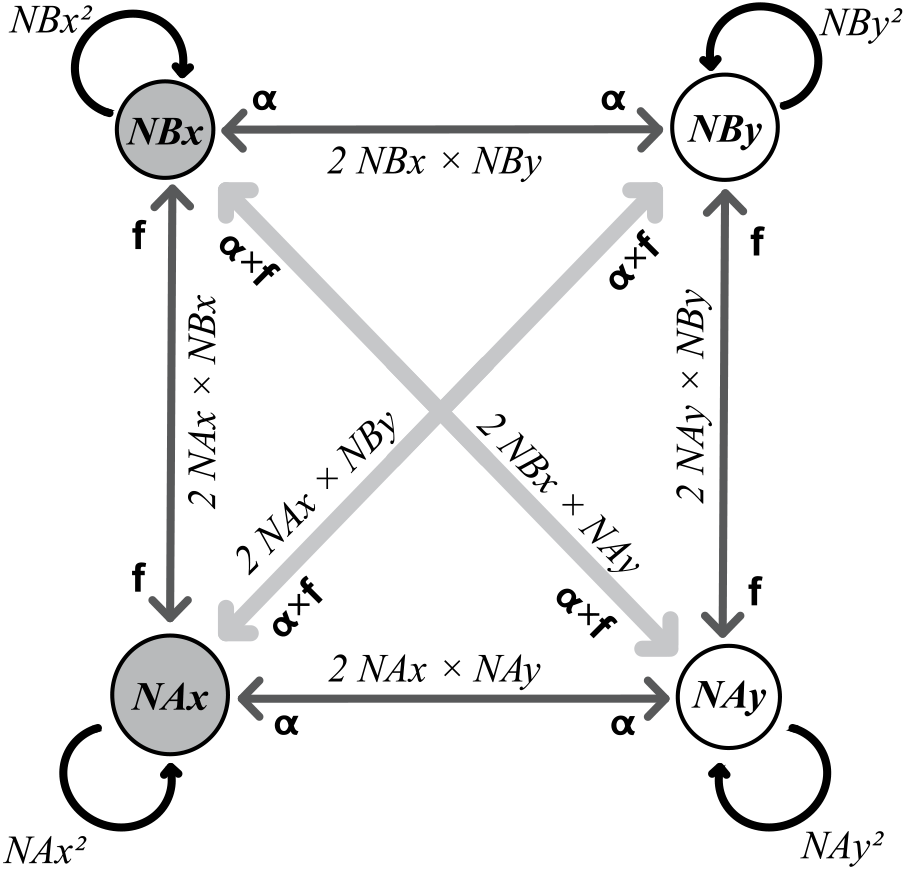
Mathematical model of a 2-mating-bias-allele, 2-ecological-niche sympatric ecosystem.

**Fig M4.**
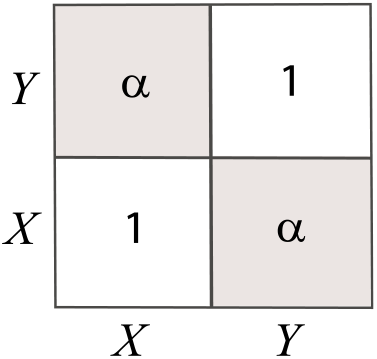
A matching compatibility table for mating-bias alleles. Same-allele individuals (*X* or *Y*) always match with a probability of 1 (no barrier); different-allele individuals match with probability *α* (barrier strength 1 − *α*).

**Fig M5.**
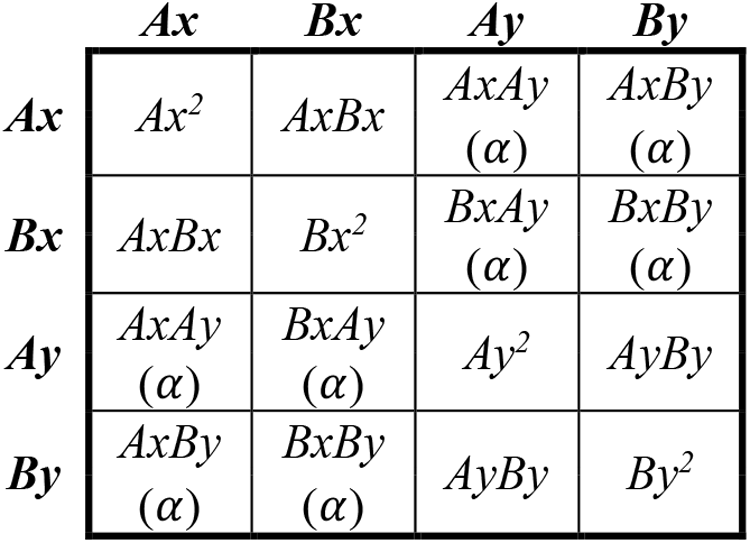
Encounter Probability Matrix (*M*) for a sympatric model with no viable hybrids.

**Fig M6.**
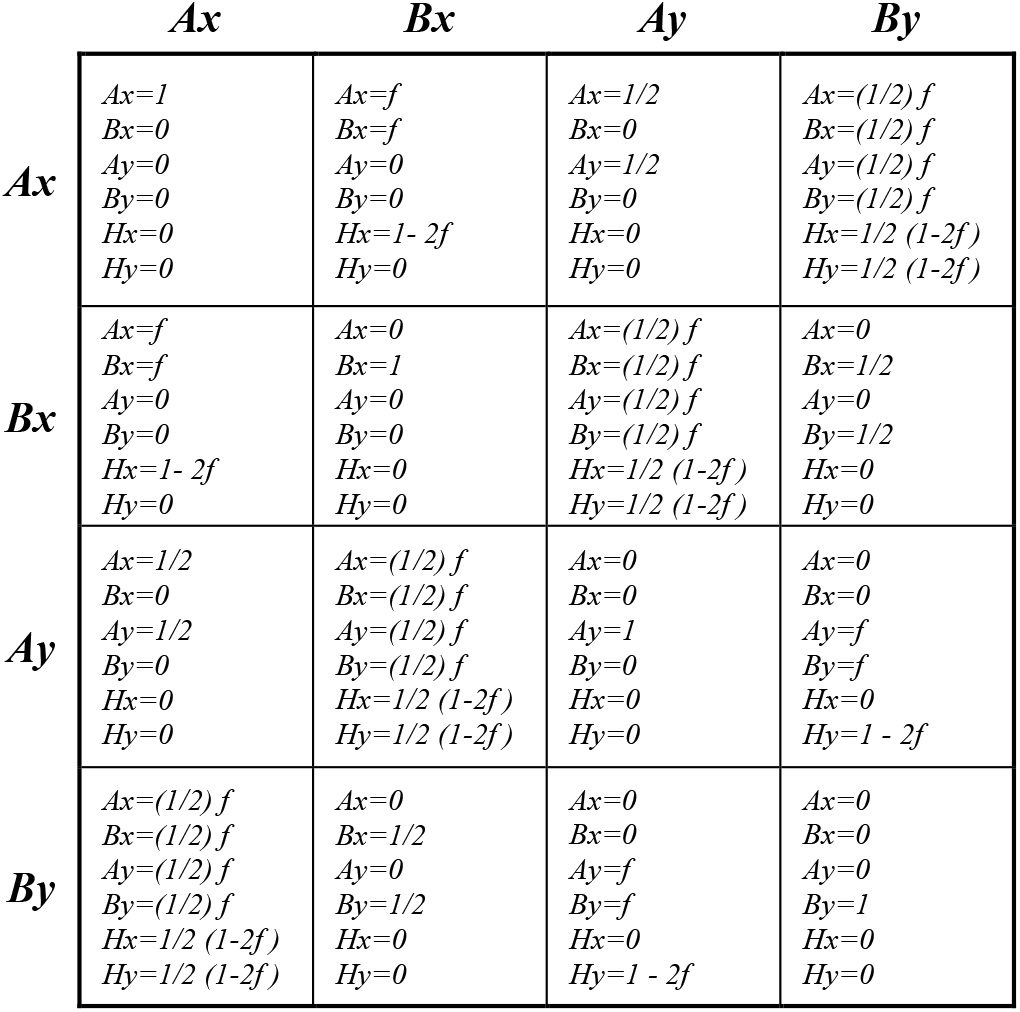
Unit Offspring Matrix (*U*).

### 2. Mathematical models of habitat-preference barriers

Next, we developed mathematical models to describe the one-allele and two-allele models of habitat-preference barriers in a sympatric population under disruptive ecological selection. These models were implemented under two spatial configurations: an open-space system and a no-open-space system.

To conceptualize the open-space system, imagine a group of sympatric cichlid fish living in a lake. The lake has two niche habitats, *A* and *B*, that are small relative to the size of the lake. The cichlids have two ecotypes that are most adaptive to the two spatially separated niche habitats, while their hybrid offspring are maladaptive and have decreased viability. The strength of disruptive ecological selection against hybrids is specified by the variable *f*, the offspring return ratio, which has a value ranging from 0 to 0.5. When the niche-*A* and niche-*B* ecotypes are not feeding in their adapted niches, they swim in the open water of the lake (free-roaming), where they can encounter one another and pair off to produce offspring. Because the niche sizes are small relative to the lake size, we assume that niche-*A* ecotypes do not visit the niche-*B* habitat, and niche-*B* ecotypes do not visit the niche-*A* habitat.

The no-open-space system configuration can be conceptualized using the sympatric speciation of hawthorn and apple maggot flies as an illustrative example. Imagine a group of sympatric fruit flies living in an orchid. The orchid contains two niche habitats, *A* and *B*, which represent hawthorn and apple trees. The flies have two ecotypes that are best adapted to these two distinct niche habitats, while their hybrid offspring are maladaptive and exhibit decreased viability. Similar to the open-space model, the strength of disruptive ecological selection against hybrids is specified by the variable *f*. The flies randomly fly among trees to feed on fruits. Therefore, niche ecotypes can visit each other’s habitats. Even though there is open space between the apple and hawthorn trees, the flies have little incentive to linger there, because flying requires a lot of energy. As a result, they encounter one another only on trees (i.e., either habitat *A* or *B*), where mating and reproduction occur.

The habitat-preference model adopts the same life cycle for the sympatric population as shown in Fig M1. However, in this case, the sexual selection phase is omitted; that is, the population is assumed to be panmictic, with random mating. This simplification is implemented by setting *α* = 1 in the mating-bias model (Fig M4), effectively removing any mating bias between alleles.

In the one-allele model of habitat preference, a single allele, *K*, confers a habitat-preference bias *θ*, ranging from 0 to 1. The value of *θ* specifies the probability that an individual carrying the *K* allele is “free-roaming,” while 1 − *θ* indicates the probability of being “home-bound.” For example, if *θ* = 0.2, then 80% of individuals with the K allele remain in their adaptive habitats, while 20% roam freely, as if they had no habitat preference. In this model, the same *K* allele confers habitat preference for ecotypes in both niches.

By contrast, the two-allele model requires separate alleles to confer habitat preferences in different niche ecotypes. Here, two alleles—*K* and *J*—are defined, each with its own habitat-specific preference bias: *θ*1 for *K* in niche *A*, and *θ*2 for *J* in niche *B*.

Next, instead of using the Encounter Probability Matrix in Fig M5—which specifies the probabilities of random encounters—new encounter matrices are derived to reflect the habitat preferences imposed by different habitat-preference barrier mechanisms.

Fig M7 shows the Encounter Probability Matrix for habitat-preference barriers in an open-space system. In each generation, individuals eligible for mating are classified into four habitat-preference groups—*AH, AF, BH*, and *BF*—representing the normalized population ratios of ecotypes (“*A*” or “*B*”) and their movement status: home-bound (“*H*”) or free-roaming (“*F*”). These group ratios are derived from the habitat-preference alleles and their associated bias values within each ecotype group. Because all parametric values in the model are normalized, *AH* + *AF* + *BH* + *BF* = 1.

The schematic diagram in Fig M8 illustrates how different habitat-preference groups may encounter one another in an open-space system. To compute the Encounter Probability Matrix (Fig M7), individuals are allocated to different mating pools based on their ecotypes and habitat preferences, as defined by their habitat-preference group. For example, individuals in groups *AH* and *BH* can only meet others within their native habitats, whereas those in groups *AF* and *BF* may encounter others within their own habitats or roam into open space to meet. These constraints determine which groups may encounter one another.

To calculate the encounter probabilities for the *AF* group, we assume that individuals from this group encounter others at rates proportional to the number of individuals present in the mating pools they can access. Applying this same assumption across all groups yields a system of simultaneous equations. Solving this system of equations provides the optimal encounter probability ratios among the habitat-preference groups and ensures that all individuals encounter another individual, with no one left unpaired. See the Appendix for the full mathematical derivation.

Similarly, for the no-open-space system, Fig M9 shows the corresponding Encounter Probability Matrix, and Fig M10 presents the schematic diagram illustrating how the different habitat-preference groups may encounter one another. In this system, individuals can only meet within habitats *A* or *B*. Those that prefer to remain in their adaptive habitats are assigned to either group *AH* or *BH*. Free-roaming individuals from both habitats (*AF* and *BF*) are grouped together into a single category, *FF*. We assume that individuals in group *FF* encounter others in habitats at rates proportional to the number of individuals present in each habitat. Solving the resulting system yields the optimal encounter probability ratios among the habitat-preference groups and ensures that all individuals are matched. See the Appendix for the full mathematical derivation.

### 3. Computer simulations of the coupling between habitat-preference and mating-bias barriers

All GUI applications in this study follow the same life cycle algorithms as outlined in Figures M1 and M2. Coupling an additional barrier mechanism to an existing one is achieved either by modifying the mating-bias value (α) for a mating-bias barrier (Fig M4) or by substituting the appropriate Encounter Probability Matrix for a habitat-preference barrier in the computation—i.e., replacing the random-encounter matrix shown in Fig M5 with a matrix from either Fig M7 or Fig M9.

Once eligible individuals encounter one another and successfully pair based on the compatibility of their mating-bias alleles, all matched parental pairs produce the next generation through random assortment of alleles at each gene locus. The resulting offspring genotype and hybrid ratios are calculated using a Unit Offspring Matrix—analogous to the one shown in Fig M6—based on the strength of ecological selection, specified by the parameter *f*.

To examine the invasion and coupling dynamics of various barrier mechanisms, our general approach is to introduce a mutant allele that embodies the characteristics of a specific barrier type. We then vary model parameters to determine the conditions under which the mutant can successfully invade and proliferate in the presence of preexisting barriers. Successful invasion indicates successful coupling, and the resulting changes in population dynamics and overall RI are visualized through the model’s phase portraits.

For instance, modeling the coupling of a mating-bias barrier following the establishment of an initial habitat-preference barrier begins by disabling mating bias in the population (i.e., setting *α* = 1). A mutant allele with habitat preference is then allowed to invade the sympatric population under disruptive ecological selection until the system reaches equilibrium and initial RI is established. Subsequently, the mating-bias barrier is introduced by assigning a nonzero *α* value, thereby allowing a high-mating-bias mutant allele to invade. The coupled dynamics of both barriers are then analyzed using the updated phase portraits.

Conversely, to couple a habitat-preference barrier after the establishment of a mating-bias barrier, the process is reversed. Once the initial mating-bias barrier has reached equilibrium, the habitat-preference barrier is introduced by replacing the random encounter matrix in Fig M5 with the appropriate Encounter Probability Matrix (Fig M7 or M9). The resulting invasion and coupling dynamics are then displayed and analyzed using the corresponding phase portraits.

Fig M11 summarizes all GUI applications developed in this study. Table M1 provides an overview of how the various habitat-preference barriers are implemented within these applications.

**Fig M7.**
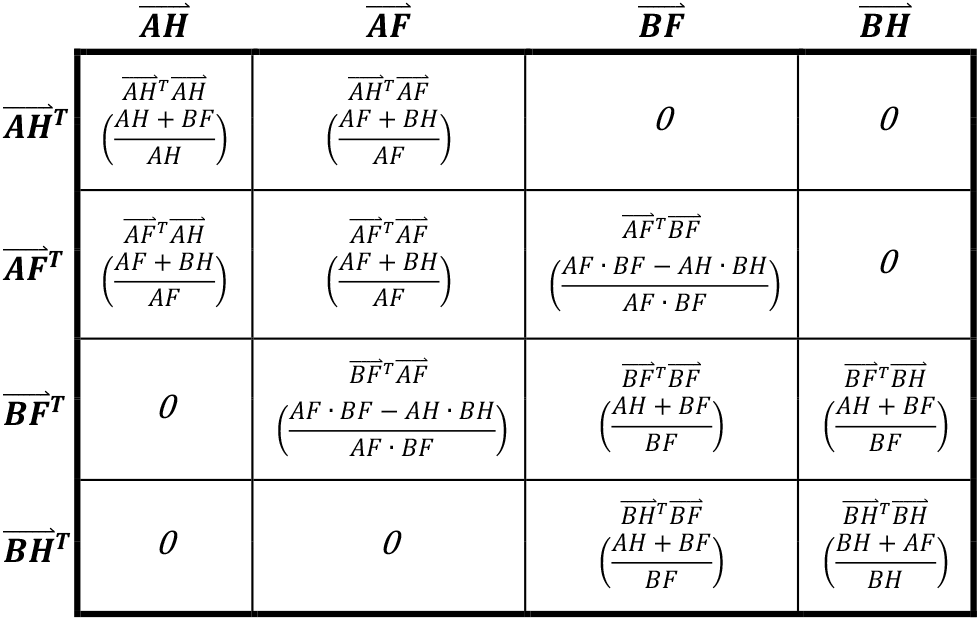
Encounter Probability Matrix for habitat-preference barriers in an open-space system. See Appendix for derivations.

**Fig M8.**
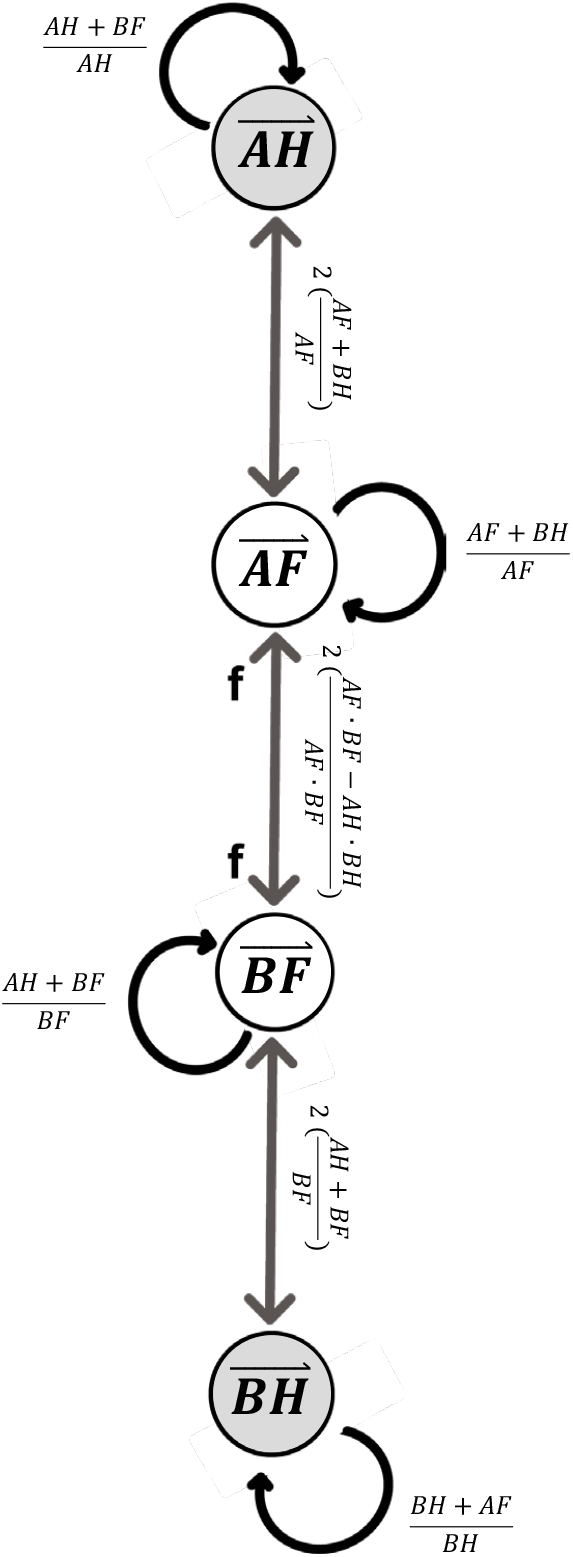
Mathematical model of habitat-preference barriers in an open-space system. See Appendix for derivations.

**Fig M9.**
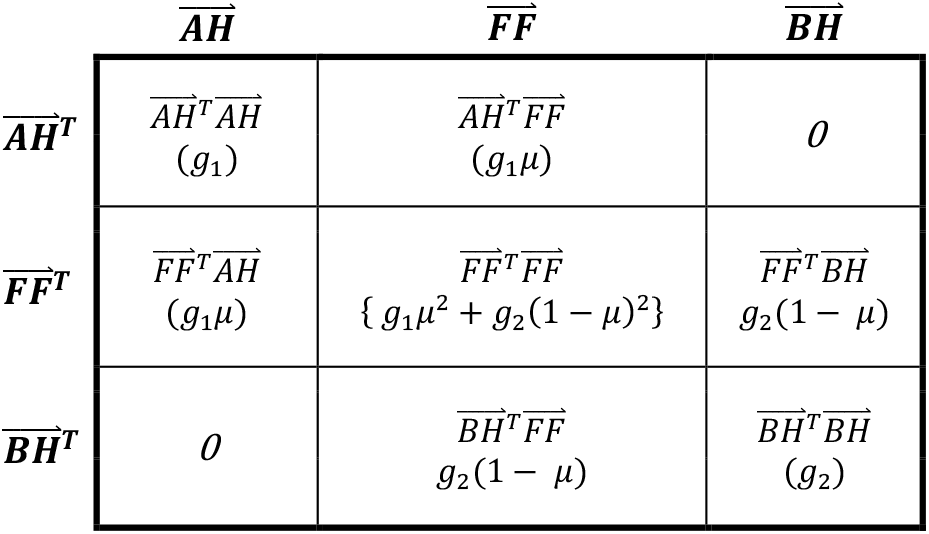
Encounter Probability Matrix for habitat-preference barriers in a no-open-space system. See Appendix for derivations.

**Fig M10.**
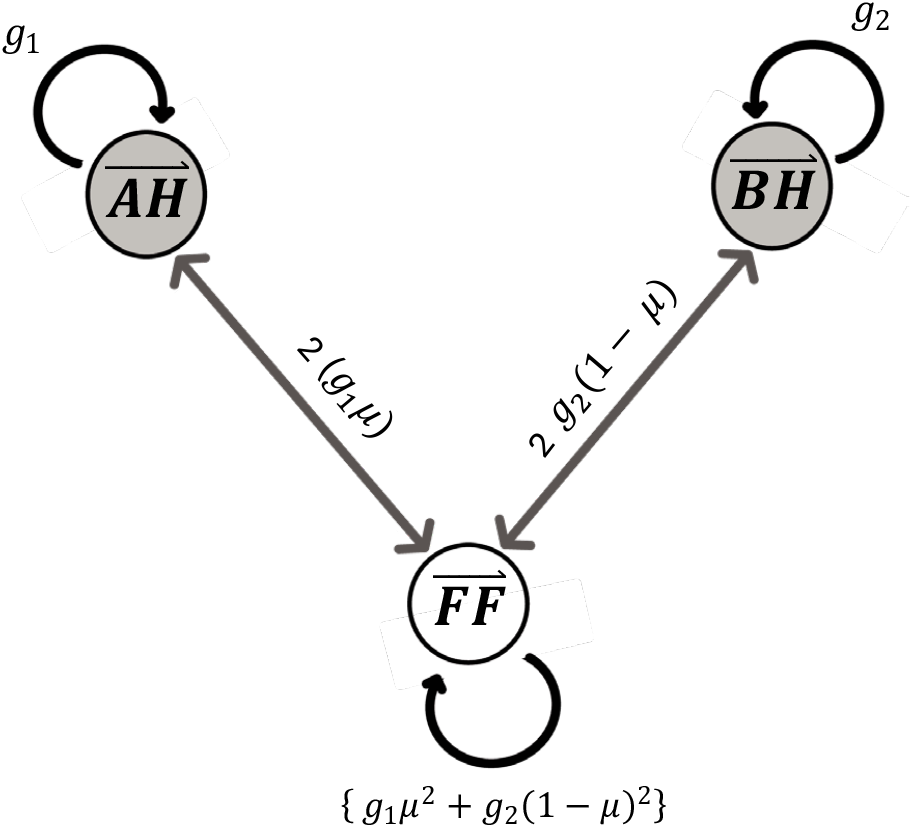
Mathematical model of habitat-preference barriers in a no-open-space system. See Appendix for derivations.

**Fig M11.**
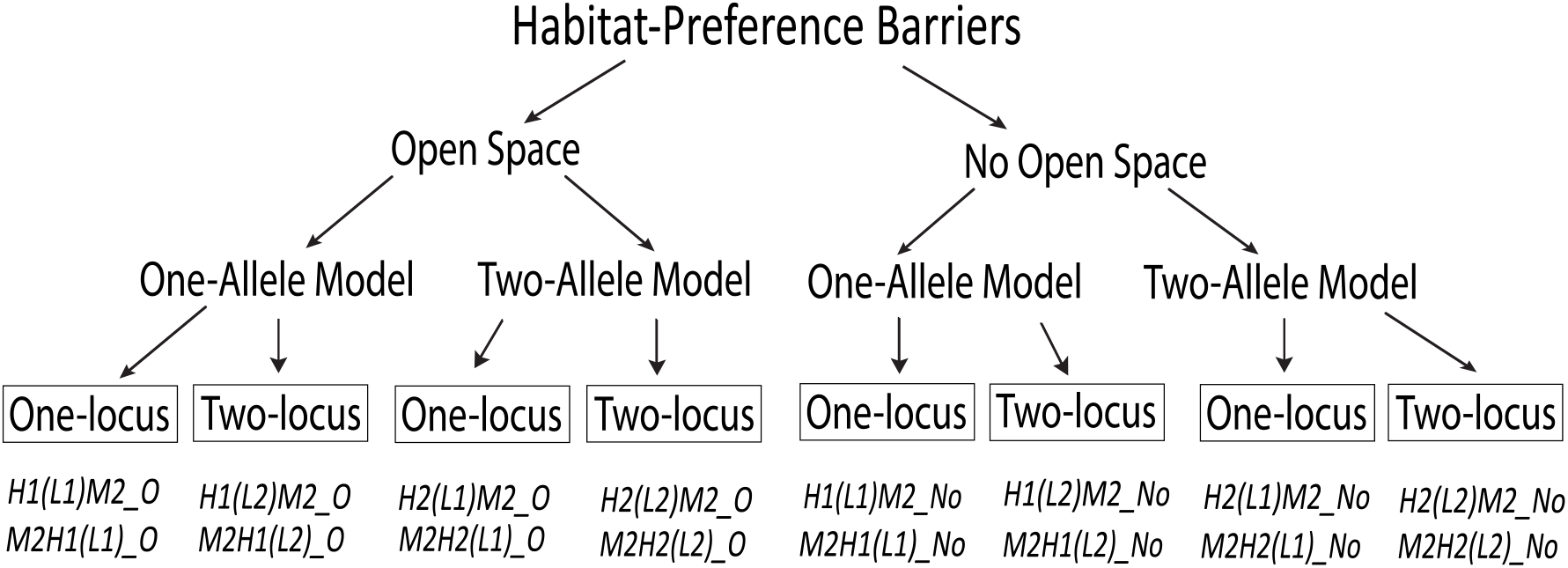
Summary of the GUI applications used to model the coupling between habitat-preference barriers and mating-bias barriers. Habitat-preference barriers can be based on either a one-allele (*H*1) or two-allele (*H*2) model, with one-gene-locus (*L*1) or two-gene-locus (*L*2) structures, in either an open-space system or a no-open-space system. The names of the GUI applications indicate the types of barriers modeled and the order in which they are coupled. Specifically, *H*1 denotes a one-allele model, *H*2 a two-allele model; *L*1 and *L*2 indicate one- and two-gene-locus barriers, respectively. *M*2 represents a two-allele mating-bias barrier. “*O*” refers to an open-space system, and “*No*” refers to a no-open-space system. For example, *H*1(*L*1)*M*2_*O* indicates that *H*1(*L*1) is the first habitat-preference barrier, followed by *M*2 as the second barrier, in an open-space system.

**Table M1.** Summary of how habitat-preference barriers are modeled in the GUI applications.

|  |
| --- |
| **1-allele, one-gene-locus habitat-preference barrier, $H1(L1)$:** A single allele $K$ at a single gene locus affects individuals in both habitats, $A$ and $B$. If present, it causes individuals of the $A$ ecotype and the $B$ ecotype to prefer staying in their respective habitats with a probability of $1 - \theta$. This means that only a proportion $\theta$ of individuals carrying the habitat-preference mutation are inclined to free-roam. The absence of the $K$ allele at the same gene locus is denoted by an inert $J$ allele, which confers no habitat preference. |
| **1-allele, 2-gene-locus habitat-preference barrier, $H1(L2)$:** Two gene loci determine habitat preference. The habitat-preference allele at each gene locus can have different effects in different niches. The first gene locus has allele $K$ or $nk$ ($nk$ means the absence of $K$). If present in the $A$ ecotype, allele $K$ causes the $A$ ecotype to want to stay in habitat $A$ with a probability of $1 - \theta ka$. If present in the $B$ ecotype, allele $K$ causes the $B$ ecotype to want to stay in habitat $B$ with a probability of $1 - \theta kb$. The presence of $nk$ has no effect on the $A$ ecotype or the $B$ ecotype. Similarly, the second gene locus has allele $J$ or $nj$ ($nj$ means the absence of $J$). If present in the $B$ ecotype, allele $J$ causes the $B$ ecotype to want to stay in habitat $B$ with a probability of $1 - \theta jb$. If present in the $A$ ecotype, allele $J$ causes the $A$ ecotype to want to stay in habitat $A$ with a probability of $1 - \theta ja$. The presence of $nj$ has no effect on the $A$ ecotype or the $B$ ecotype. An individual can have the following combinations of habitat-preference alleles in its genome: $[K\ nj]$, $[nk\ J]$, $[nk\ nj]$, and $[K\ J]$. |
| **2-allele, 1-gene-locus habitat-preference barrier, $H2(L1)$:** A single gene locus has two alleles, $K$ and $J$. If present in the $A$ ecotype, allele $K$ causes the $A$ ecotype to want to stay in habitat $A$ with a probability of $1 - \theta 1$. The $K$ allele has no effect on the $B$ ecotype. Similarly, if present in the $B$ ecotype, allele $J$ causes the $B$ ecotype to want to stay in habitat $B$ with a probability of $1 - \theta 2$. The $J$ allele has no effect on the $A$ ecotype. An individual has either the $K$ or the $J$ allele in its genotype. It cannot have both, and it cannot have neither. |
| **2-allele, 2-gene-locus, habitat-preference barrier, $H2(L2)$:** Two gene loci determine habitat preference. The first gene locus has allele $K$ or $nk$ ($nk$ means the absence of $K$). If present in the $A$ ecotype, allele $K$ causes the $A$ ecotype to want to stay in habitat $A$ with a probability of $1 - \theta 1$. The $K$ allele has no effect on the $B$ ecotype. The presence of $nk$ has no effect on the $A$ ecotype or the $B$ ecotype. Similarly, the second gene locus has allele $J$ or $nj$ ($nj$ means the absence of $J$). If present in the $B$ ecotype, allele $J$ causes the $B$ ecotype to want to stay in habitat $B$ with a probability of $1 - \theta 2$. The $J$ allele has no effect on the $A$ ecotypes. The presence of $nj$ has no effect on the $A$ ecotype or the $B$ ecotype. An individual can have the following combinations of habitat-preference alleles in its genome: $[K\ nj]$, $[nk\ J]$, $[nk\ nj]$, and $[K\ J]$. |

## III. RESULTS

### Invasion Dynamics and Reversibility of Habitat-Preference BArriers

We developed computer applications to simulate the invasion dynamics of a habitat-preference mutant allele in a two-habitat ecosystem under disruptive ecological selection. Matings between ecotypes adapted to each of the two niche habitats produce maladaptive hybrids with reduced viability. This hybrid loss then drives the invasion and population dynamics of the habitat-preference alleles to decrease inter-niche mating encounters, resulting in premating RI between niche populations.

We analyzed both one- and two-allele models of habitat-preference barriers in open-space and no-open-space habitat structures. In general, our results show that as long as hybrid loss persists in the system, a mutant allele associated with habitat preference can always invade to reduce hybrid loss to the extent allowed by its habitat-preference bias, *θ*. This holds true for both one-allele and two-allele models. Because one-allele models are not affected by the homogenizing effects of gene flow, they tend to evolve more rapidly and generate stronger habitat-preference RI than two-allele models. Lastly, we investigate the reversibility of habitat-preference barriers when disruptive ecological selection is removed.

#### 1. Open-space habitat-preference models

Fig 1a shows the computer screenshot of a MATLAB GUI application that we developed to calculate and display the phase portrait solutions of a one-allele, one-gene-locus habitat-preference barrier in an open-space system. As described in the Methodology section, this system is inspired by sympatric cichlid fishes living in a lake with two niche habitats, *A* and *B*, which are small relative to the overall lake size. Two niche ecotypes are each adapted to one of the two habitats, while their hybrid offspring suffer reduced viability due to maladaptation. The strength of disruptive ecological selection against hybrids is specified by the variable *f*, the offspring return ratio, which ranges from 0 to 0.5. When not foraging in their respective niche habitats, individuals from both ecotypes can swim into the open water, where inter-niche encounters and mating may occur. Because the niche habitats are small and spatially separated, we assume that niche-*A* and niche-*B* ecotypes do not visit each other’s habitats.

In a one-allele model (Fig 1a), a habitat-preference mutant allele, *K*, has an associated habitat-preference bias, *θ*, with a value ranging from 0 to 1. The value of *θ* specifies the likelihood that an individual carrying the *K* allele is free-roaming (i.e., free to swim around, as specified by the ratio *θ*) or habitat-bound (i.e., stays in its adapted habitat, as specified by the ratio 1 − *θ*).

At the beginning of each mating generation, we normalize the four possible genotype groups in the population—*Ak, Aj, Bk*, and *Bj*—based on their niche ecotypes and the habitat-preference alleles they carry (here, the *J* allele, denoting the absence of the *K* allele, is functionally inert and confers no habitat preference to the ecotypes carrying it). We then use a matching algorithm to calculate the encounter probabilities between the four different genotype groups in the open-space structure. This allows us to determine the different types of matched parental pairs and the offspring genotype ratios they will produce for the next generation.

As shown in Fig 1a, a small population of mutant *K* allele in niche *A* (*Ak* = 0.01) is able to invade a sympatric population under disruptive ecological selection (*f* = 0.125) in a two-niche open-space system, become fixed in both niche *A* and niche *B* after 200 generations, and establish premating RI by reducing inter-niche encounters.

**Fig 1a.**
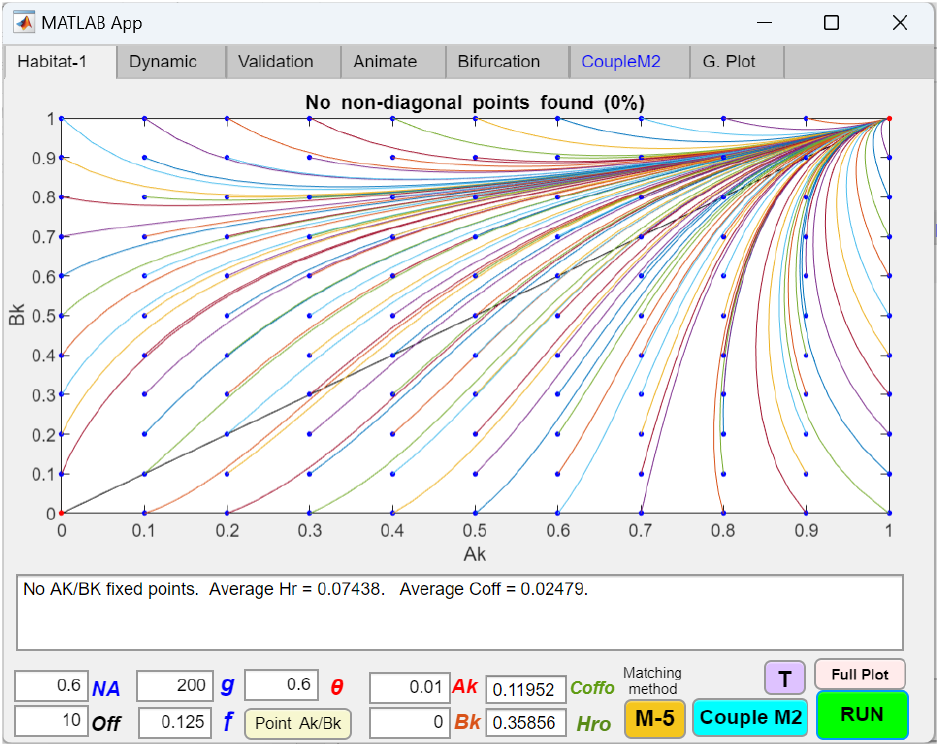
One-allele, one-gene-locus model of a habitat-preference barrier (H1/L1, O) in an open-space system, showing fixation of the habitat-preference allele in both niches. A computer screenshot of a MATLAB GUI application shows the phase portrait solution of a one-allele, one-gene-locus habitat-preference barrier in an open-space system. In this example, a mutant *K* allele confers a habitat-preference bias, denoted by *θ*. When *θ* = 0.6, an individual carrying the *K* allele remains in its home habitat 40% of the time and free-roams 60% of the time. *Ak* and *Bk* represent the proportions of the *K* allele in niches *A* and *B*, respectively. The absence of the *K* allele at the same gene locus is denoted by an inert *J* allele, which confers no habitat preference. Because all values are normalized, the proportions of the *J* allele in the two niches are *Aj* = 1 − *Ak* and *Bj* = 1 − *Bk*. The rest of the parametric variables are the same as those described in the Methodology section. *f* is the offspring return ratio that reflects the strength of ecological selection. *g* is the number of generations. *NA* is the normalized niche-*A* carrying capacity; therefore, the niche-*B* carrying capacity is *NB* = 1 − *NA. off* represents the number of offspring each parent can produce. The model assumes that each mating generation always produces more offspring than the niche carrying capacities can support, ensuring that at the beginning of each generation, the population ratios in niches *A* and *B* remain fixed. *Coff* is the ratio of non-hybrid offspring produced from inter-niche mating, and *Hr* is the ratio of hybrid offspring produced from inter-niche mating. The total ratio of inter-niche offspring produced in a mating generation is *Coff* + *Hr*. Here, the phase portrait plots the dynamic population vector trajectories of *Ak* and *Bk* over *g* generations. The blue dots represent the initial *Ak*/*Bk* population ratios, while the red dots indicate the final *Ak*/*Bk* population ratios after *g* generations. As shown, a starting mutant genotype population of *Ak* = 0.01/*Bk* = 0 can invade and become fixed in both niche *A* and niche *B* (i.e., *Ak* = *Bk* = 1) after 200 generations. Consequently, the invasion of the *K* allele lowers the pre-invasion *Coff* (*Coffo*) from 0.11952 to *Coff* = 0.02479 and the pre-invasion hybrid ratio (*Hro*) from 0.35856 to *Hr* = 0.07438, indicating reduced inter-niche mating encounters and resulting in premating RI due to the habitat-preference barrier.

**Fig 1b.**
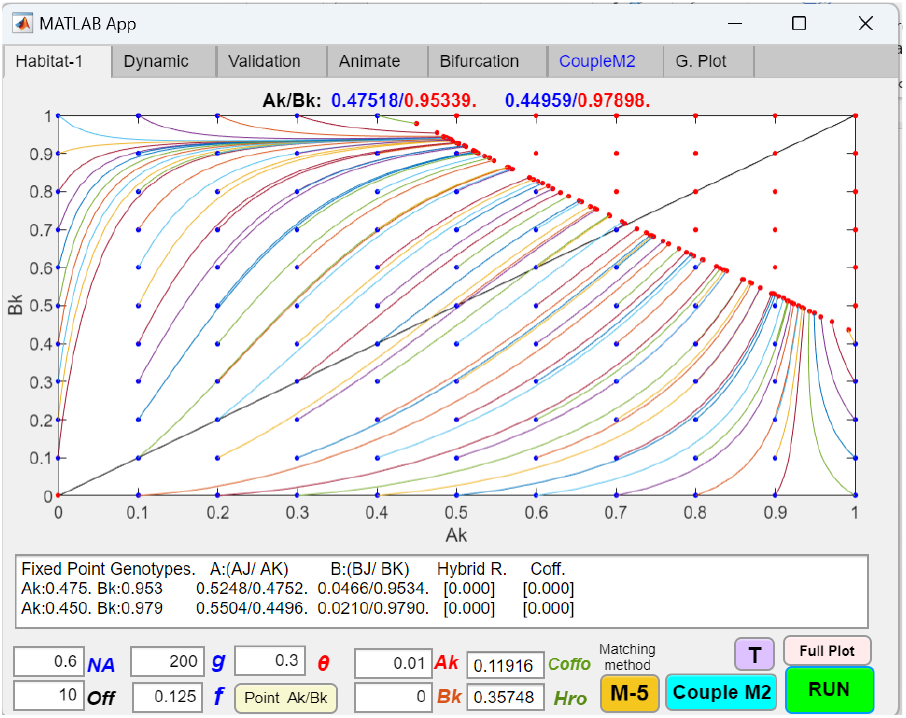
One-allele, one-gene-locus model of a habitat-preference barrier (H1/L1, O) in an open space system, showing population vector trajectories stopping at points where *Hr* = 0. The habitat-preference bias, *θ*, in Fig 1a is reduced from 0.6 to 0.3, strengthening the habitat preference associated with the *K* allele. The phase portrait solution shows that population vector trajectories halt at points where *Hr* reaches zero. There is a region in the upper right corner where the population ratios remain stationary. At these stationary points, enough individuals prefer to remain in their own niches so that free-roaming individuals in each niche no longer feel the need to venture into open space to find mates. Staying within their own niches and matching with their ecotypes ensures that all individuals can find a mate and that *Hr* = 0, so the population ratios remain unchanged because there is no hybrid loss.

Our simulation results reveal that the invasion of a habitat-preference allele is driven by hybrid loss, measured by the ratio *Hr*. As long as *Hr* is greater than zero, a habitat-preference barrier can always invade to minimize *Hr*. The caveat is that even when *Hr* is zero, it does not always mean no inter-niche mating has occurred or that gene flow between the two niche ecotypes is absent. This is because the total offspring ratio from inter-niche mating is the sum of the hybrid offspring ratio (*Hr*) and the non-hybrid offspring ratio (*Coff*). In our models, *Coff* represents the ratio of offspring from inter-niche mating that have the same genotypes as the parents. The total inter-niche offspring ratio, *Hr* + *Coff*, is determined by encounter probabilities between the two niche ecotypes, while the offspring return ratio, *f*, determines the relative proportions of *Hr* and *Coff* within this total inter-niche offspring ratio.

By definition, if *f* = 0.5, then *Hr* = 0, and the total offspring ratio from inter-niche mating is simply *Coff*. Conversely, when *f* = 0, *Coff* = 0, and *Hr* represents the total offspring ratio from inter-niche mating. Nevertheless, for a given value of *f*, the relative ratio between *Hr* and *Coff* is fixed. Therefore, as habitat-preference barriers reduce inter-niche matching encounters, they decrease the total offspring produced by inter-niche mating, which is then reflected by a proportional decrease in *Hr*. Thus, for any given value of *f* < 0.5, a decrease in *Hr* can be used as a measure of the relative strength of RI produced by different habitat-preference barriers.

As long as *Hr* (maladaptive hybrid loss) is not zero, habitat-preference barriers can always invade to eliminate the hybrid loss, to the extent allowable by their associated habitat-preference bias, *θ*. In the Fig 1a example, the habitat-preference *K* allele is able to invade, become fixed in both niches, and reduce *Hr* from 0.35856 to 0.07438, leading to premating RI between the niches due to reduced inter-niche matching encounters.

In Fig 1b, we strengthen the habitat-preference bias of the *K* allele by reducing *θ* in Fig 1a from 0.6 to 0.3 while keeping all other parameter values unchanged. The phase portrait solution shows that population vector trajectories halt at points where *Hr* reaches zero, indicating that hybrid loss is no longer present to drive the invasion of the habitat-preference allele. There is a region in the upper right corner where the population ratios remain stationary. At these stationary points, enough individuals prefer to stay in their own niches that free-roaming individuals no longer need to venture into open space outside their niches to find mates. In this matching arrangement, all individuals can find mates within their adapted niches, and *Hr* remains zero, so no hybrid loss exists to drive changes in population ratios.

Fig 2a shows the invasion dynamics of a two-allele model of habitat-preference barriers in an open-space system. In a two-allele model, different niche ecotypes must carry different habitat-preference alleles to exhibit the preference effect. In Fig 2a, a single gene locus has two habitat-preference alleles, *K* and *J*, with associated habitat-preference biases, *θ*1 and *θ*2. An individual either carries the *J* allele or the *K* allele; it cannot have both, nor can it have neither. A niche-*A* ecotype carrying the *K* allele has the *Ak* genotype, and the presence of the *K* allele causes an *Ak* individual to prefer staying in niche *A* with probability *θ*1. When carried by a niche-*B* ecotype, the *K* allele has no effect on the *Bk* genotype; therefore, a *Bk* individual is always free-roaming. Similarly, the presence of the *J* allele causes a *Bj* individual to prefer staying in niche *B* with probability *θ*2. The *J* allele has no effect on the *Aj* genotype; therefore, an *Aj* individual is always free-roaming.

In Fig 2a, with *f* = 0.125 and *θ*1 = *θ*2 = 0.6, the phase portrait shows population vector trajectories converging on a fixed point (*Ak* = 0.83334, *Bk* = 0.16667) by generation 200. Consequently, the hybrid offspring ratio (*Hr*) declines from 0.21456 to 0.120, indicating the establishment of premating RI due to reduced inter-niche matching encounters. Increasing habitat-preference biases by decreasing *θ*1 and *θ*2 shifts the fixed point toward the bottom right corner of the phase portrait, where *Ak* = 1 and *Bk* = 0, further reducing inter-niche encounters and strengthening RI. When *θ*1 and *θ*2 are set to one, the niche ecotypes have no habitat preferences, and all population vectors settle on stationary points along the diagonal line (where *Ak* = *Bk*), with equal ratios of the *K* and *J* alleles in both niches.

**Fig 2a.**
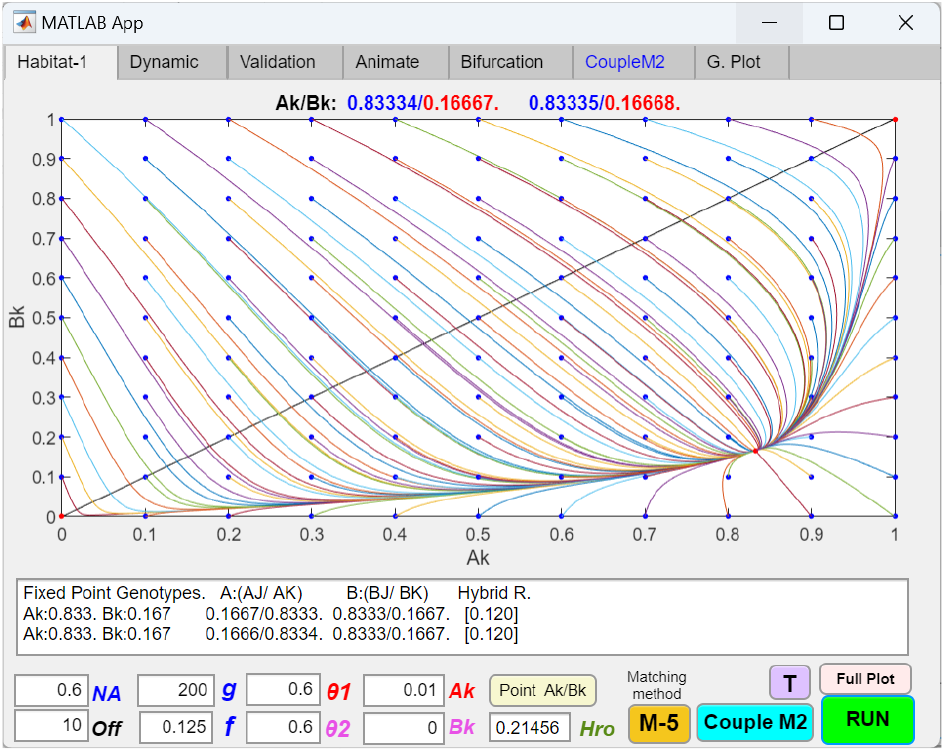
Two-allele, one-gene-locus model of a habitat-preference barrier (H2/L1, O) in an open-space system, showing fixed-point polymorphism of the habitat-preference alleles. A single gene locus has two habitat-preference alleles, *K* and *J*. An individual either carries the *K* allele or the *J* allele; it cannot have both, nor can it have neither. All parameters are normalized. Therefore, the genotype ratio of the niche-*A* ecotype carrying the *K* allele is *Ak*, and the genotype ratio of the niche-*A* ecotype carrying the *J* allele is 1 − *Ak*. The same genotype ratio relationships apply to niche-*B* individuals carrying the *J* or *K* alleles and their corresponding *Bj* and *Bk* genotypes. If present in the niche-*A* ecotype, the *K* allele causes the niche-*A* ecotype to prefer staying in habitat *A* with probability 1 − *θ*1. The *K* allele has no effect on the niche-*B* ecotype; thus, *Bk* is always free-roaming. Similarly, if present in the niche-*B* ecotype, the *J* allele causes the niche-*B* ecotype to prefer staying in habitat *B* with probability 1 − *θ*2. The *J* allele has no effect on the niche-*A* ecotype; thus, *Aj* is always free-roaming. The phase portrait solution shows the *K* allele invading and reaching a fixed-point polymorphism in both niches. This reduces *Hr* from 0.21456 to 0.120 and establishes premating RI between niche ecotypes by decreasing inter-niche matching encounters.

However, if *θ*1 and *θ*2 are further reduced, a pattern analogous to the phase portrait in Fig 1b emerges. As shown in Fig 2b, when *θ*1 and *θ*2 in Fig 2a are decreased from 0.6 to 0.3, all vector trajectories stop at points where *Hr* reaches zero. A region of stationary population ratios appears near the bottom right corner of the phase portrait. Similar to the stationary region in Fig 1b, all individuals in this region are able to find matches in their own niches without the need to venture out of their adaptive habitats to find mates; *Hr* remains zero, so no hybrid loss is present to drive the movement of existing population ratios.

**Fig 2b.**
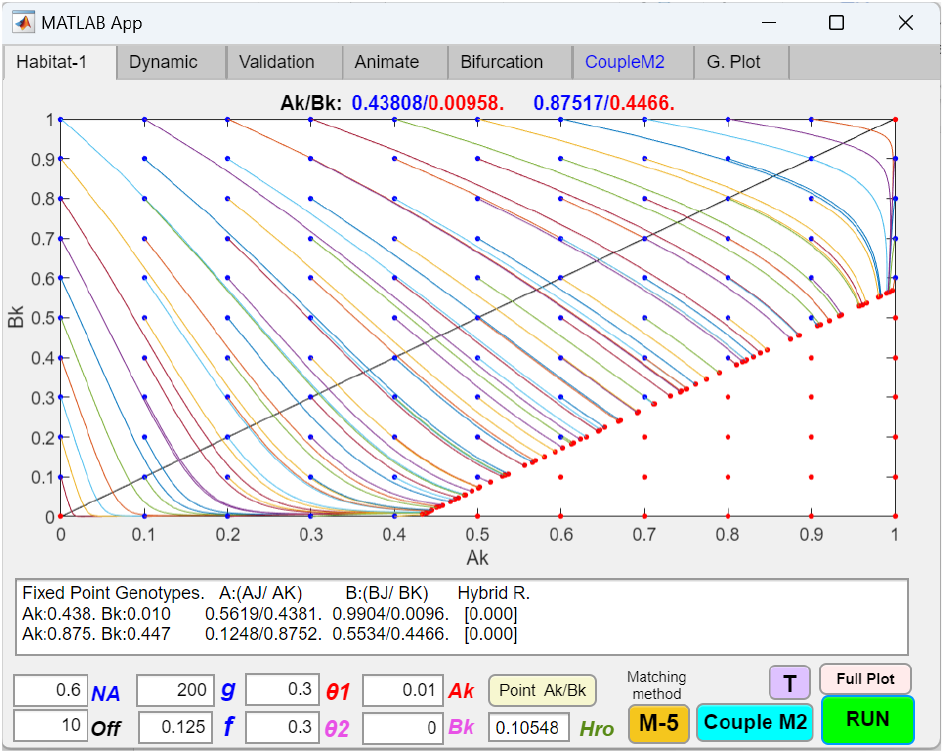
Two-allele, one-gene-locus model of a habitat-preference barrier (H2/L1, O) in an open-space system, showing population vector trajectories stopping at points where *Hr* = 0. The habitat-preference biases, *θ*1 and *θ*2, in Fig 2a are reduced from 0.6 to 0.3, strengthening the habitat preferences associated with the *J* and *K* alleles. The phase portrait solution shows that population vector trajectories stop at points where *Hr* reaches zero. Similar to the phase portrait in Fig 1b, a region in the lower right corner exhibits stationary population ratios. At these stationary points, enough individuals prefer to remain in their own niches such that free-roaming individuals in each niche no longer feel the need to venture into open space to find mates.

Notice that vector trajectories can never stop above the diagonal line, as this would imply negative habitat preference—i.e., an increased tendency to free-roam—which cannot occur when *θ* values are less than or equal to 1 (meaning habitat preference remains positive). As long as there is hybrid loss, the tendency of the system is to always drive the equilibrium points toward the lower right corner to eliminate the hybrid loss.

#### 2. No-open-space habitat-preference models

Fig 3 shows a computer screenshot of a MATLAB GUI application that calculates and displays the phase portrait solutions of a one-allele, one-gene-locus habitat-preference barrier in a no-open-space system. As described in the Methodology section, this system is inspired by the sympatric speciation of hawthorn and apple maggot flies. Imagine an ancestral fly species living in an orchard with two niche habitats— hawthorn and apple trees (habitats *A* and *B*). The population contains two niche ecotypes, each adapted to one of the two habitats, while their hybrid offspring are maladaptive and suffer reduced viability. The strength of disruptive ecological selection is again specified by the offspring return ratio, *f*. The flies move randomly among trees, allowing ecotypes to enter each other’s habitats. Although open space exists between trees, the flies have little incentive to linger in midair, as flight is energetically costly. As a result, all encounters and mating take place on trees (i.e., in habitat *A* or *B*).

Fig 3 shows the invasion dynamics of a one-allele, one-gene-locus habitat-preference mutant allele *K* in a no-open-space system. The parametric values are the same as those in Fig 1a. Overall, the results are similar to those of the open-space system; however, because free-roaming ecotypes from the opposite niche can directly visit native ecotypes in their niche without having to meet in a communal open-space forum, inter-niche mating always occurs and the hybrid ratio (*Hr*) is never zero— unless, of course, *θ* = 0, which causes all individuals to remain in their adaptive niches. Consequently, a region of stationary population ratios, as shown in Fig 1b, does not appear in the phase portrait of a no-open-space system. As long as there is maladaptive hybrid loss in the system, the habitat-preference *K* allele can always invade and go to fixation in both niches to reduce this hybrid loss.

Fig 4a shows the invasion dynamics of the same two-allele habitat-preference model in Fig 2b, but in a no-open-space system. Again, because free-roaming ecotypes can visit all niches, inter-niche mating always occurs and hybrid loss is never zero—unless all the preference biases are zero (i.e., *θ*1 = *θ*2 = 0), in which case the fixed point moves to the bottom right corner of the phase portrait, where *Ak* = 1 and *Bk* = 0, and there is complete RI. A region of stationary population ratios, such as that shown in Fig 2b, never appears. As shown in Fig 4b, when all preference biases are set to 1 (i.e., *θ*1 = *θ*2 = 1), habitat preference is absent in the population, and all population vectors converge to stationary points along the diagonal line. As the habitat preferences, *θ*1 and *θ*2, are gradually decreased from 1 toward zero, fixed points appear that gradually move away from the diagonal line toward the bottom right corner of the phase portrait, where habitat-preference mediated RI is strongest.

#### 3. Two-gene-locus models of habitat-preference barriers

We developed two-gene-locus models for the one- and two-allele habitat-preference barriers in open-space and non-open-space systems to investigate the coupling of additional habitat-preference barriers after an initial habitat-preference barrier has been established. The results are consistent with our expectation that as long as hybrid loss remains after establishing a first habitat-preference barrier, additional barriers can always be coupled to reduce this hybrid loss and further augment overall RI.

**Fig 3.**
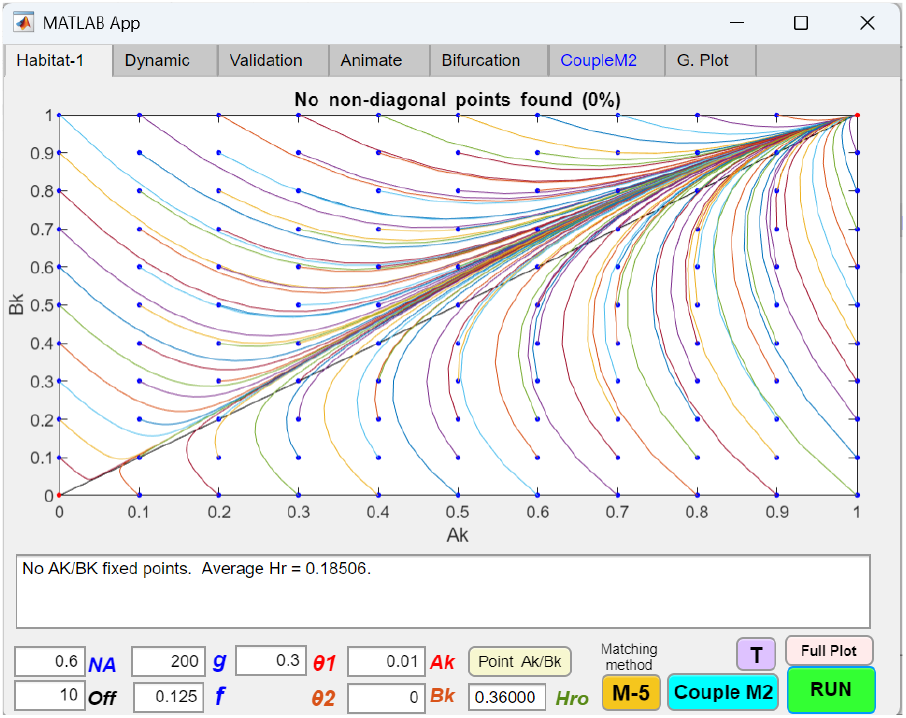
One-allele, one-gene-locus model of a habitat-preference barrier (H1/L1, No) in a no-open-space system, showing fixation of the habitat-preference allele in both niches. A computer screenshot of a MATLAB GUI application displays the phase portrait solution of a one-allele, one-gene-locus habitat-preference barrier in a no-open-space system. The parametric variables are the same as in Fig 1b. Overall, the results for the no-open-space system resemble those of the open-space system, except that even with strong habitat preference (low *θ*), a region of stationary population ratios, such as the one in Fig 1b, does not appear. The *K* allele always goes to fixation in both niches as long as *Hr* > 0, indicating that a habitat-preference mutant allele can always invade and establish premating RI as long as maladaptive hybrid loss exists in a sympatric population under disruptive ecological selection.

**Fig 4a.**
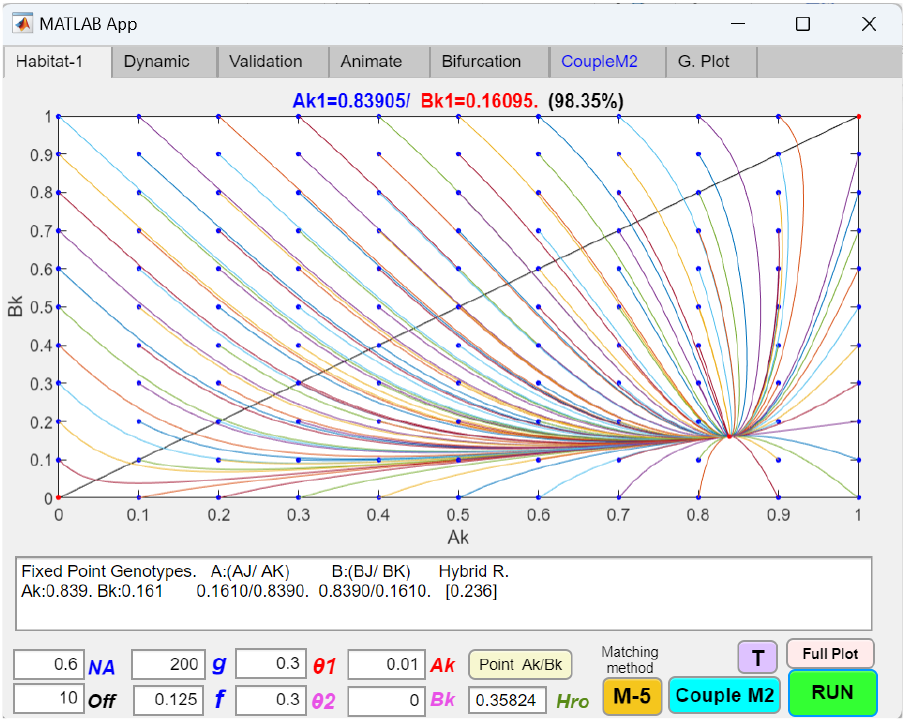
Two-allele, one-gene-locus model of a habitat-preference barrier (H2/L1, No) in a no-open-space system, showing fixed-point polymorphism of the habitat-preference alleles. The same variables are used as in Fig 2b. A single gene locus contains two habitat-preference alleles, *K* and *J*, with associated habitat-preference biases *θ*1 and *θ*2, which are effective in niche *A* and niche *B*, respectively. Overall, the phase portrait solutions for the no-open-space system resemble those for the open-space system, except that even with strong habitat preferences (low *θ*1 and *θ*2), no region of stationary population ratios, such as the one in Fig 2b, appears. As long as *Hr* > 0, the two-allele habitat-preference barrier can always invade and establish fixed-point polymorphism and premating RI. As preference biases decrease from 1 to 0, the fixed points in the phase portrait shift from the diagonal line to the bottom-right corner, where *Ak* = 1, *Bk* = 0, and *Hr* reaches zero.

**Fig 4b.**
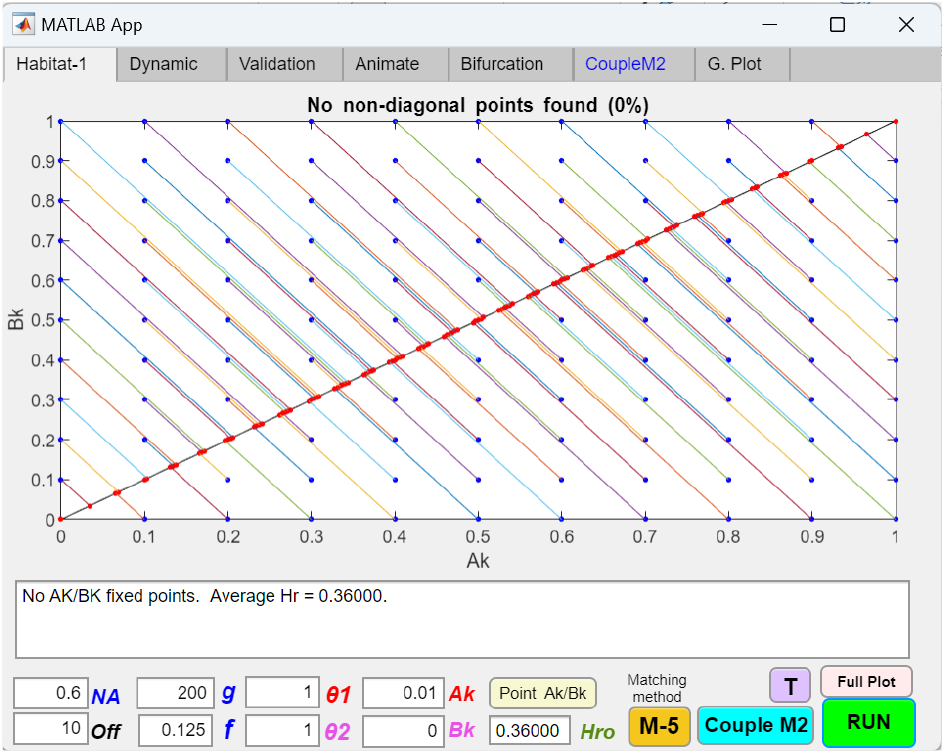
Two-allele, one-gene-locus model of a habitat-preference barrier (H2/L1, No) in a no-open-space system, showing the population vectors converging on the diagonal line when *θ*1 = *θL* = 1, indicating the absence of habitat preference. When the values of *θ*1 and *θ*2 in Fig 4a are set to one, individuals in the population exhibit no habitat preference, and gene flow homogenizes the allele ratios in both niches. As a result, all vector trajectories converge on the diagonal line, where *Ak* = *Bk* and *Aj* = *Bj*.

Overall, the dynamic behaviors of the two-gene-locus models closely mirror those of the one-gene-locus models, with only minor differences in the invasion dynamics of the two-allele models between the open-space and no-open-space systems during the establishment of initial RI.

Fig 5a shows a two-allele, two-gene-locus model of a habitat-preference barrier (H2/L2, O) in an open-space system. In this model, a *K* allele at the first gene locus induces habitat-preference bias (*θ*1) only in niche-*A* ecotypes, while a *J* allele at the second gene locus induces habitat-preference bias (*θ*2) only in niche-*B* ecotypes. As shown in Fig 5a, when the second gene locus is disabled by setting *θ*2 = 1, a mutant *K* allele in niche *A* successfully invades and becomes fixed in both niches, reducing the hybrid ratio (*Hr*) from 0.35856 to 0.21719 and establishing initial premating RI. Subsequently, Fig 5b shows that after the establishment of this initial barrier by the *K* allele at the first gene locus, a mutant *J* allele emerging in niche *B* can invade and couple at the second gene locus, further reducing *Hr* and strengthening RI.

In contrast, in a no-open-space system, shown in Fig 6a, when the second gene locus is disabled, a small mutant genotype population *Ak* = 0.01 cannot invade. This occurs because, in a no-open-space system without habitat preference, all niche-*B* ecotypes in niche *B* can visit niche-*A* ecotypes in niche *A*. As a result, the *Ak* genotype loses the fitness advantage it would otherwise gain from reduced inter-niche mating due to its habitat preference for niche *A*.

**Fig 5a.**
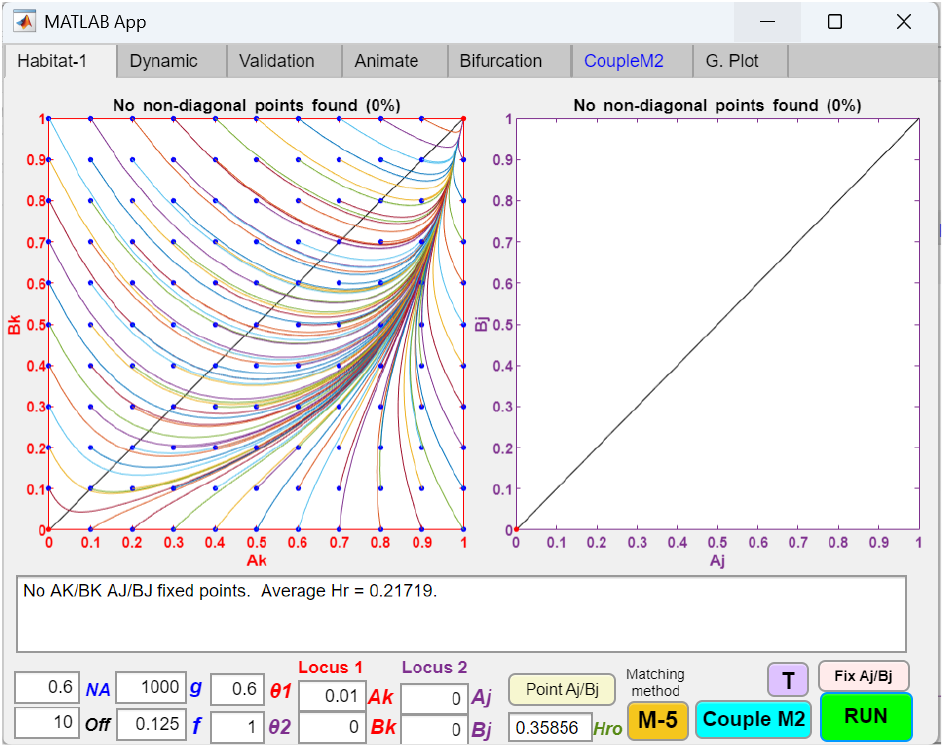
Two-allele, two-gene-locus model of an initial habitat-preference barrier (H2/L2, O) in an open-space system, showing the invasion of a mutant allele that affects habitat preference in only one niche. In a two-allele, two-gene-locus model, two gene loci determine habitat preference in an open-space system. The first gene locus has either allele *K* or *nk* (*nk* denotes the absence of *K*). When carried by the niche-*A* ecotype, the *K* allele causes the *Ak* genotype to prefer staying in habitat *A* with a probability ratio of 1 − *θ*1. The *K* allele has no effect when carried by the niche-*B* ecotype, and the *nk* allele has no effect when carried by either the niche-*A* or niche-*B* ecotype. Similarly, the second gene locus has either allele *J* or *jn*. The *Bj* genotype prefers to stay in niche *B* with a probability ratio of 1 − *θ*2. The *J* allele has no effect when carried by the niche-*A* ecotype, and the nj allele has no effect when carried by either the niche-*A* or niche-*B* ecotype. The phase portrait solution of the model shows that when the second gene locus is disabled by setting *θ*2 = 1 and *Aj* = *Bj* = 0, a small mutant population *Ak* = 0.01 at the first gene locus can invade and reach fixation at *Ak* = 1 and *Bk* = 1, reducing the initial hybrid ratio from 0.35856 to 0.21719 and establishing initial premating RI.

However, as soon as habitat preference emerges in niche-*B* ecotypes, the *Ak* genotype gains a fitness advantage to invade. As shown in Fig 6b, in the presence of a small mutant population *Bj* = 0.01, both the *Ak* and *Bj* mutant populations can invade, become fixed in both niches, and establish initial premating RI.

This underscores the greater challenge of establishing initial RI in the two-allele model of habitat preference compared to the one-allele model. While the one-allele model is immune to the homogenizing effects of gene flow, the two-allele model requires the evolution of two separate mutations, each effective in a different niche. Notably, once an initial habitat-preference barrier is established—i.e., some niche-*B* ecotypes already exhibit habitat preference—the *Ak* mutant population in Fig 6a and 6b can always invade and couple to further strengthen RI.

**Fig 5b.**
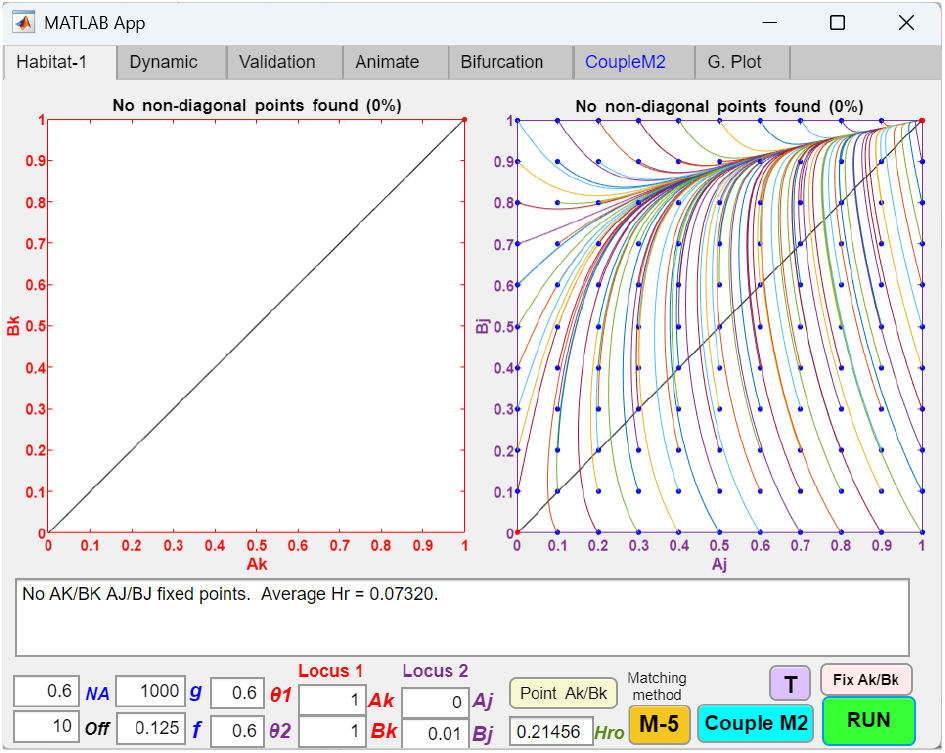
Two-allele, two-gene-locus model of an initial habitat-preference barrier (H2/L2, O) in an open-space system, showing the invasion and coupling of a second habitat-preference mutant allele at the second gene locus after an initial habitat-preference allele has become fixed at the first gene locus, resulting in increased RI. After the fixation of the initial habitat-preference *K* allele (*Ak* = *Bk* = 1) at the first gene locus in Fig 5a, the phase portrait shows that a second mutant population *Bj* = 0.01, arising at the second gene locus, can invade and become fixed at *Aj* = *Bj* = 1, further reducing the hybrid ratio to 0.07323 and increasing overall RI.

**Fig 6a.**
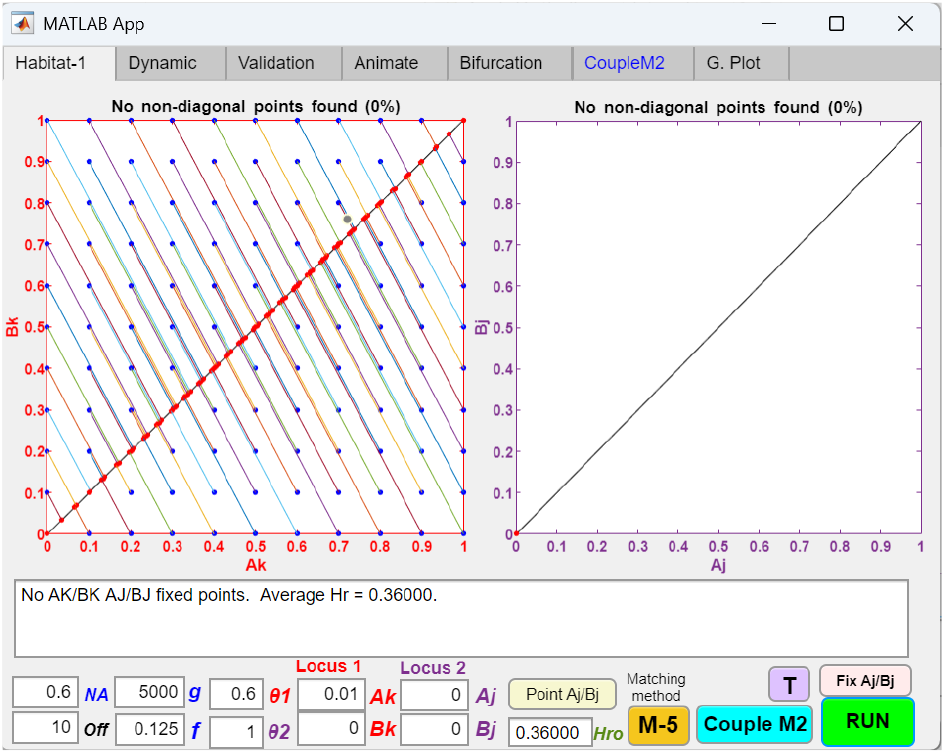
Two-allele, two-gene-locus model of an initial habitat-preference barrier (H2/L2, No) in a no-open-space system, showing a mutant allele with habitat-preference bias at a single niche habitat failing to invade. The two-allele, two-gene-locus model from Fig 5a is used to investigate the invasion dynamics of a habitat-preference mutant allele in a no-open-space system. Adopting the same parametric values as in Fig 5a, the phase portrait solution shows that a small mutant population *Ak* = 0.01 at the first gene locus cannot invade when the second gene locus is disabled by setting *θ*2 = 1 and *Aj* = *Bj* = 0.

**Fig 6b.**
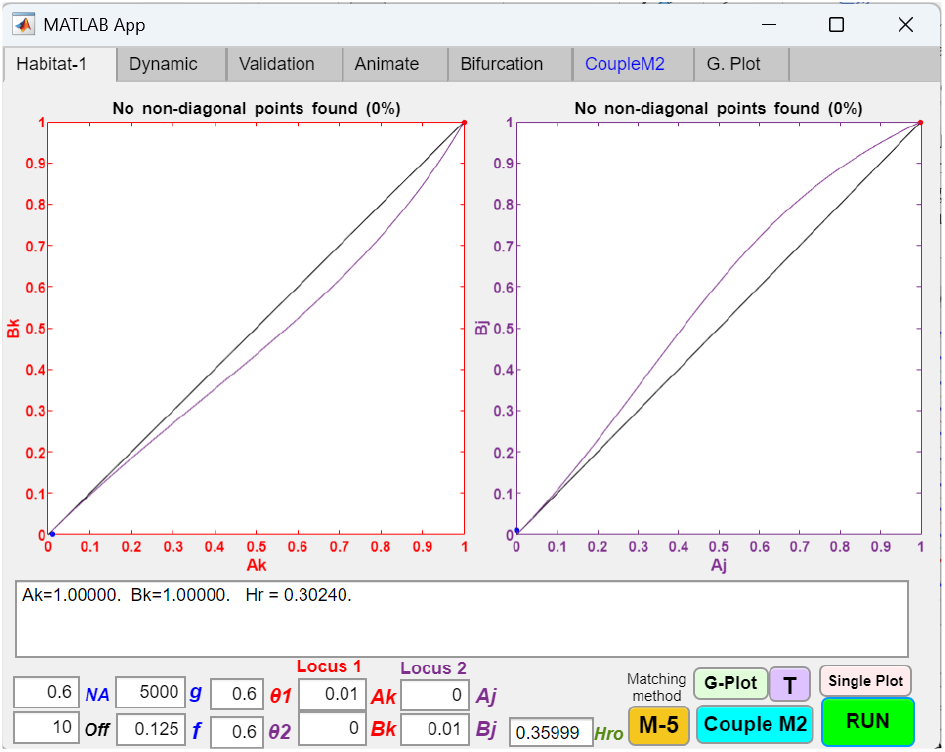
Two-allele, two-gene-locus model of an initial habitat-preference barrier (H2/L2, No) in a no-open-space system, showing that a mutant allele with habitat-preference bias at a single niche can invade when habitat-preference individuals exist in the opposite niche habitat. Given the parametric values in Fig 6a, if a small population of individuals with the *Bj* genotype (*Bj* = 0.01) exists in niche *B*, then both the *Ak* and *Bk* mutants can invade and reach fixation at *Ak* = *Bk* = 1 and *Aj* = *Bj* = 1, resulting in premating RI.

#### 4. Reversibility of the habitat-preference barriers

Next, we investigate the invasibility and the reversibility of the habitat preference barriers when external disruptive ecological selection is removed. In the open-space system, Fig 7a shows the phase portrait solution when the value of *f* in Fig 1b is increased to 0.5 in a one-allele model of habitat-preference barrier. The region of stationary population ratios in Fig 1b still persists in the phase portrait, while the rest of the population vectors converge to stationary points along the diagonal line. As *θ* is gradually increased toward 1, the area of stationary ratios in the upper-right corner gradually disappears, and when *θ* = 1, all vectors settle at stationary points along the diagonal line.

Similarly, Fig 7b shows the phase portrait when *f* in Fig 2b is increased to 0.5 in a two-allele model of habitat preference. The region of stationary ratios persists in the bottom-right corner, while the rest of the population vectors settle on a line that arcs away from the diagonal line toward the bottom-right corner. As *θ*1 and *θ*2 are increased to 1, the region of stationary ratios disappears, as does the arc, and all population vectors settle at stationary points along the diagonal line.

**Fig 7a.**
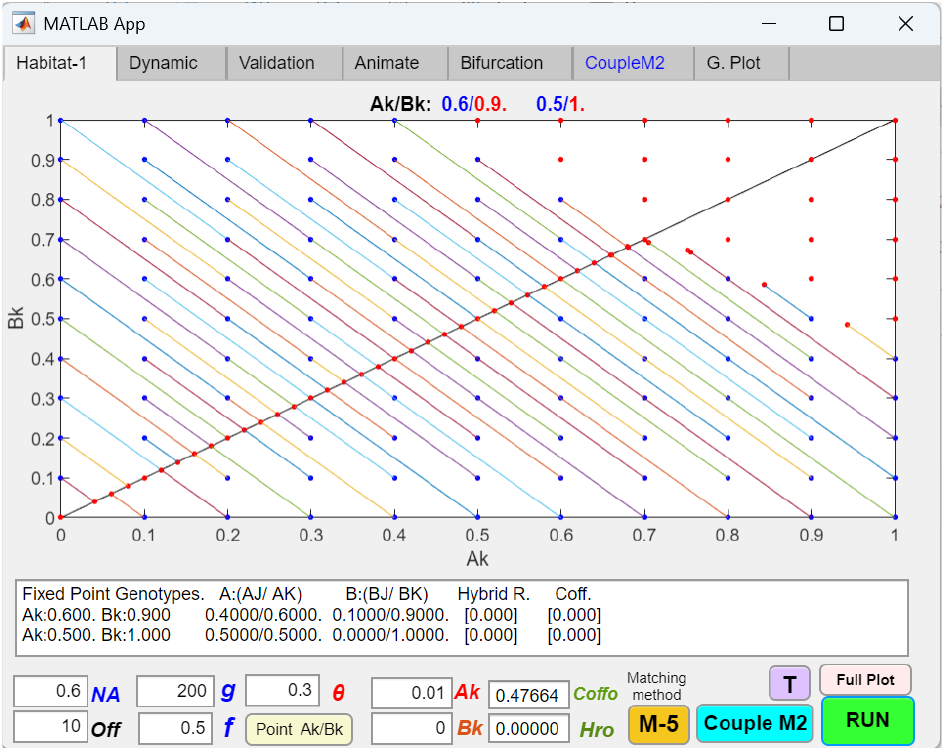
Reversibility of the one-allele, one-gene-locus model of a habitat-preference barrier (H1/L1, O) in an open-space system when disruptive ecological selection is removed. When the value of *f* in Fig 1b is increased from 0.125 to 0.5, the phase portrait shows that the upper-right area of stationary population ratios in Fig 1b persists, while the remaining population vectors move to stationary points on the diagonal line. Notice that *Hr* is always zero when *f* = 0.5. *Coff* = 0 when the *Ak*/*Bk* ratios remain at the stationary points in the upper right corner of the phase portrait. Along the diagonal line, depending on the position, the values of *Coff* range from *Coff* = 0.48 when *Ak*/*Bk* = 0/0 to *Coff* = 0 when *Ak*/*Bk* = 0.715/0.715.

**Fig 7b.**
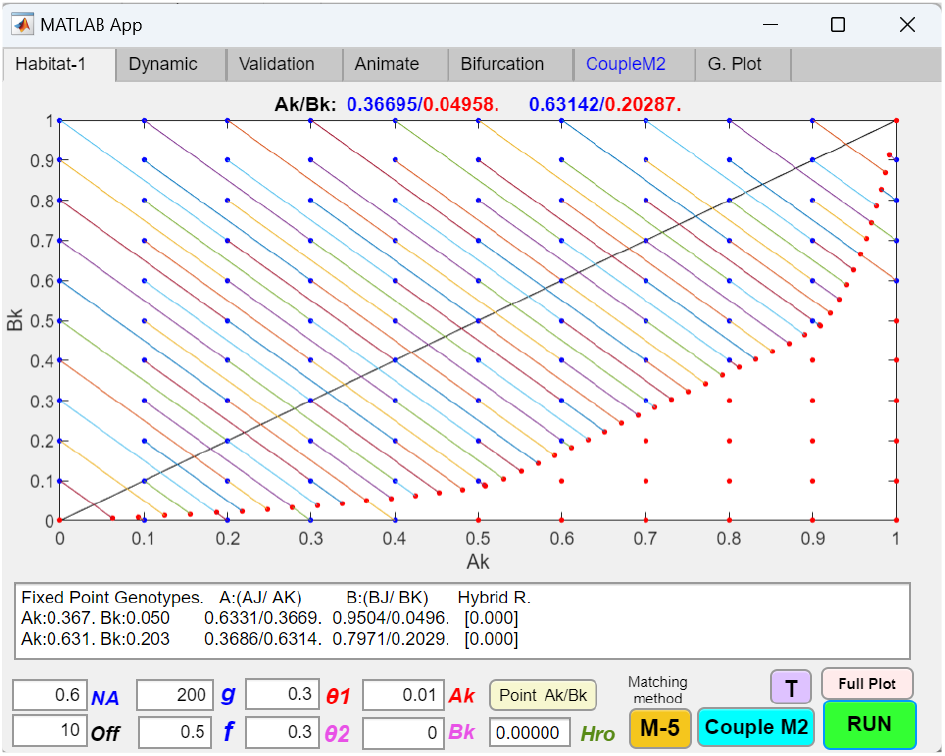
Reversibility of the two-allele, one-gene-locus model of a habitat-preference barrier (H2/L1, O) in an open-space system when disruptive ecological selection is removed. When the value of *f* in Fig 2b is increased from 0.125 to 0.5, the phase portrait solution shows that population vectors move to stationary points along a line that arcs away from the diagonal toward the bottom-right corner. The bottom-right region of stationary population ratios in Fig 2b persists in the phase portrait.

**Fig 7c.**
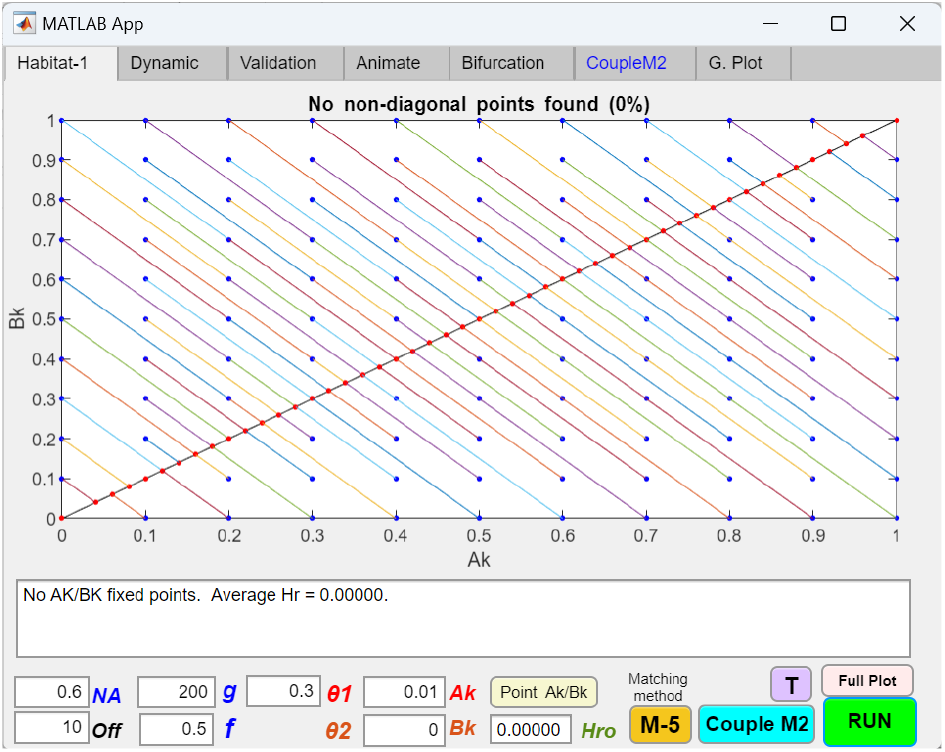
Reversibility of the one-allele, one-gene-locus model of a habitat-preference barrier (H1/L1, No) in a no-open-space system when disruptive ecological selection is removed. When the value of *f* in Fig 3 is increased from 0.125 to 0.5, the phase portrait solution shows that all population vectors move to stationary points on the diagonal line.

**Fig 7d.**
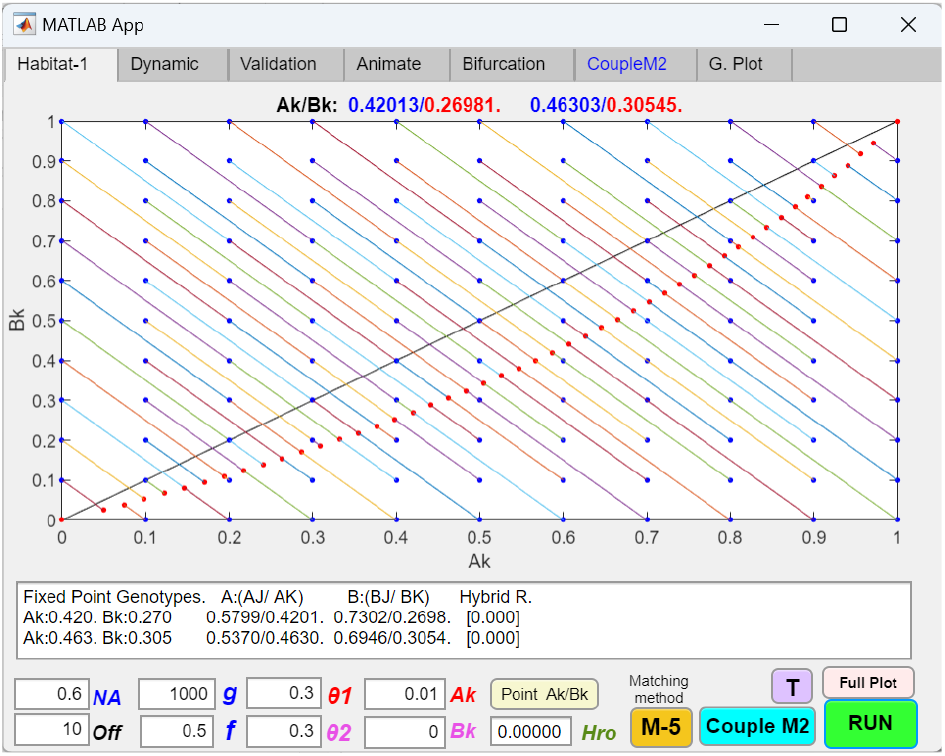
Reversibility of the two-allele, one-gene-locus model of a habitat-preference barrier (H2/L1, No) in an no-open-space system when disruptive ecological selection is removed. When the value of *f* in Fig 4a is increased from 0.125 to 0.5, the phase portrait solution shows that all population vectors move to stationary points along a line that arcs away from the diagonal toward the bottom-right corner.

In the no-open-space system, we observe similar phenomena. Fig 7c shows the phase portrait when *f* in Fig 3 is increased to 0.5 in a one-allele model of the habitat-preference barrier. No region of stationary population ratios is possible in the no-open-space system; therefore, all population vectors move to stationary points along the diagonal line.

Fig 7d shows the phase portrait when the value of *f* in Fig 4a is increased to 0.5 in a two-allele model of habitat-preference barrier. All vector trajectories converge to stationary points on an arc that bends toward the bottom-right corner. When *θ*1 and *θ*2 are increased to 1, all vector trajectories settle at stationary points along the diagonal line.

The results in Fig 7a to 7d show that when *f* = 0.5, there is no disruptive ecological selection, *Hr* = 0, and a mutant habitat-preference allele near the origin of the phase portrait cannot invade because there is no hybrid loss. However, once habitat-preference alleles reach sizable ratios in the population, they are not eliminated even when disruptive ecological selection is removed. These habitat‐preference alleles are able to maintain constant population ratios in the niches and continue to reduce inter‐niche encounters. In the two-allele models, there is no habitat-preference mediated RI on the diagonal line, where *θ*1 and *θ*2 are both 1, while RI increases as the *Ak*/*Bk* population ratios move toward the bottom-right corner. Therefore, for values of *θ*1 and *θ*2 less than 1, we can expect the population vectors to settle at ratios away from the diagonal line and toward the bottom‐right corner, indicating the presence of continuing habitat‐preference barrier effects.

The reversibility of the two-gene-locus models closely resembles that of the one-gene-locus models and does not reveal any new or unexpected findings.

#### 5. Effects of habitat-preference mutations in the same gene locus

We investigated the evolutionary dynamics of mutant alleles with stronger or weaker habitat preferences arising within the same gene locus. In general, as long as maladaptive hybrid loss persists, a more adaptive mutant allele with a stronger habitat preference will invade, eliminate the resident allele with weaker preference in the same gene locus, and result in stronger RI. This outcome holds for both one- and two-allele models. Conversely, a mutant allele with weaker habitat preference than the existing allele cannot invade and is eliminated.

Figs 8a–8b illustrate these competitive exclusion dynamics in a one-allele, one-gene-locus, open-space model of a habitat-preference barrier. Under the same parametric values as in Fig 1a, a *K*1 allele with *θ*1 = 0.6 can invade and become fixed in both niches *A* and *B*. Subsequently, when a more adaptive mutant *K*2 with stronger habitat preference (*θ*2 = 0.5) arises at an initial frequency of *Ak*2 = 0.01, it can successfully invade, eliminate *K*1, and reach fixation in both niches, thereby strengthening RI. By contrast, in Fig 8c, if the mutant *K*2 allele has weaker habitat preference (*θ*2 = 0.7) than the existing *K*1 allele (*θ*1 = 0.6), the vector-field directions of the Fig 8a phase portraits are reversed, indicating that the less adaptive *K*2 mutant cannot invade and is instead eliminated.

**Fig 8a.**
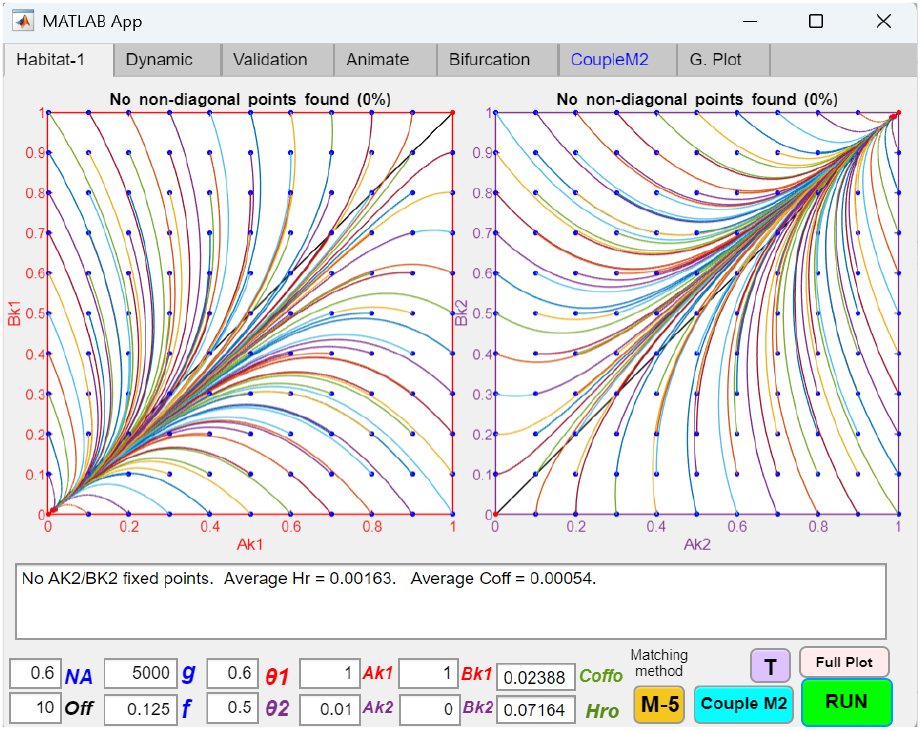
Phase portraits showing a mutant allele with stronger habitat preference replacing an existing allele with weaker habitat preference in a one-allele, one-gene-locus, open-space model of a habitat-preference barrier (H1/L1, O), resulting in stronger RI. As illustrated previously in Fig 1a, a habitat-preference allele *K*1 with *θ*1 = 0.6 can invade and rise to fixation in a one-allele, one-gene-locus open-space system. In this case, a more adaptive mutant allele *K*2 with *θ*2 = 0.5 arises and replaces *K*1. The phase portraits—plotting all possible initial population combinations of *Ak*1/*Bk*1 and *Ak*2/*Bk*2—show the shift in vector-field dynamics leading to the elimination of *K*1 and the fixation of *K*2, thereby strengthening RI.

**Fig 8b.**
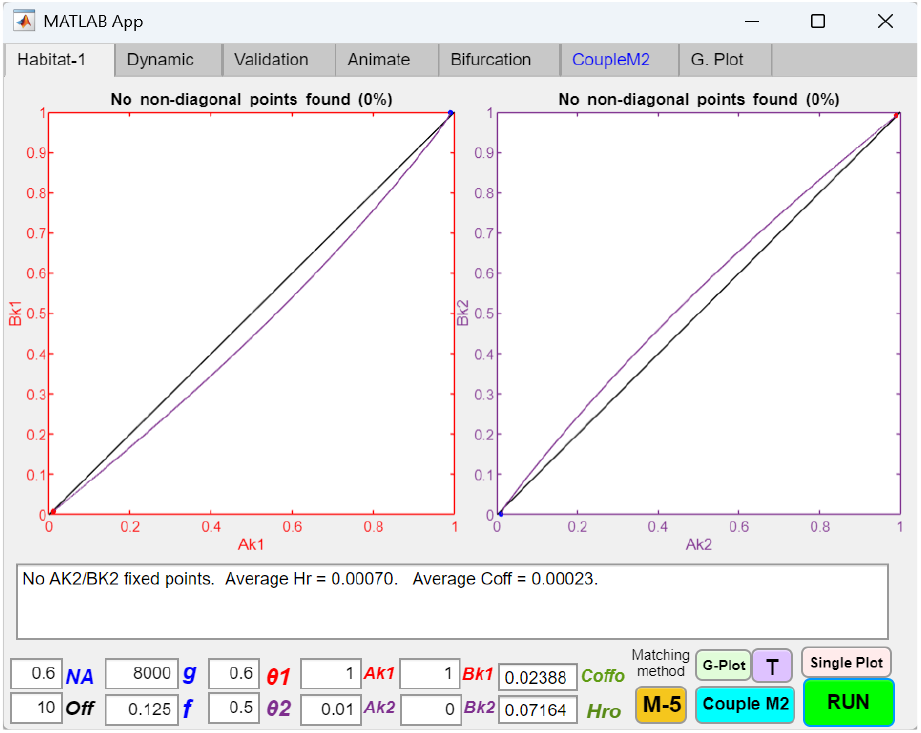
Single-plot phase portraits showing a mutant allele with stronger habitat preference eliminating and replacing an existing fixed allele with weaker habitat preference in a one-allele, one-gene-locus, open-space model of a habitat-preference barrier (H1/L1, O), resulting in stronger RI. Under the parameter values from Fig 8a, a *K*1 allele with *θ*1 = 0.6 initially invades and becomes fixed in both niches (specified by *Ak*1 = *Bk*2 = 1), reducing *Hr* to 0.07164 and *Coff* to 0.02388. Subsequently, a more adaptive mutant allele *K*2 with *θ*2 = 0.5 and an initial population of *Ak*2 = 0.01 invades, eliminating *K*1 and reaching fixation after 8000 generations. As a result, *Hr* is further reduced to 0.00070 and *Coff* to 0.00023, indicating less inter-niche mating and stronger premating RI.

In Fig 8d, a similar pattern is observed in a two-allele, one-gene-locus, no-open-space model of a habitat-preference barrier. In this case, allele *K*1 confers habitat preference *θ*1*A* = 0.3 in niche *A*, while allele *J*1 confers *θ*1*B* = 0.3 in niche *B*. Using the same parametric values as in Fig 4a, *K*1 and *J*1 can invade and establish a fixed-point polymorphism, producing premating RI. Subsequently, if *K*1 can evolve a more adaptive variant *K*2 (*θ*2*A* < 0.3), the new *K*2 mutant can invade and eliminate *K*1, strengthening RI; the same applies if *J*1 can evolve a more adaptive variant *J*2 (*θ*2*B* < 0.3). However, if *K*2 or *J*2 have weaker habitat-preference biases than their resident *K*1 and *J*1 alleles (i.e., *θ*2*A* > *θ*1*A* or *θ*2*B* > *θ*1*B*), they cannot invade and are eliminated.

**Fig 8c.**
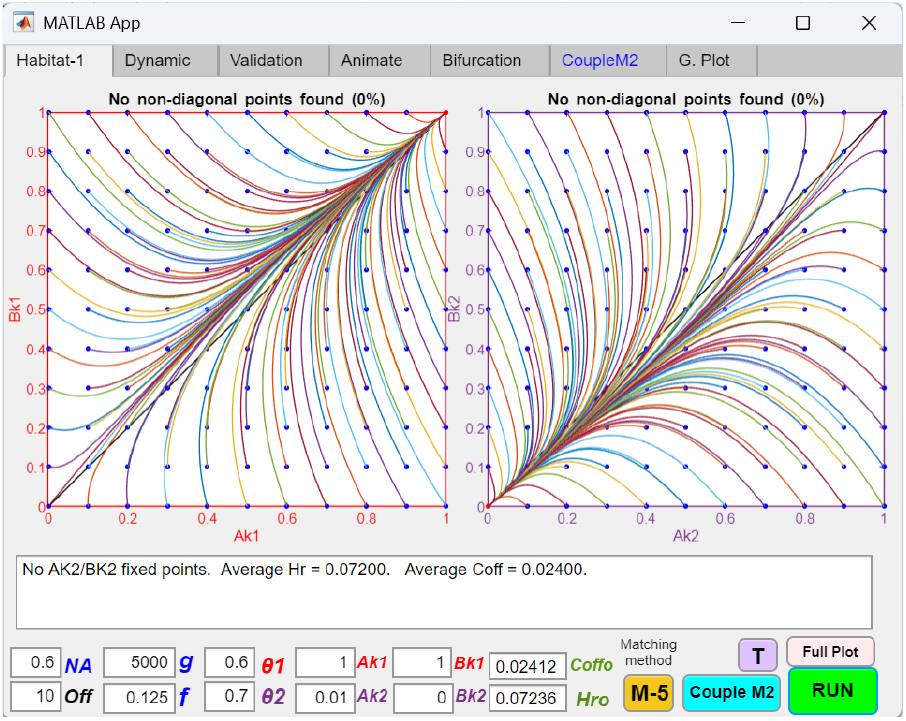
Phase portraits showing a mutant allele with weaker habitat preference failing to invade or replace an existing allele with stronger habitat preference in a one-allele, one-gene-locus, open-space model of a habitat-preference barrier (H1/L1, O), resulting in its elimination. If the *K*2 mutant allele in Fig 8a instead has a weaker habitat preference (*θ*2 = 0.7 > *θ*1 = 0.6), it is less adaptive than the existing fixed allele *K*1 and is driven to extinction by competitive exclusion at the same gene locus.

**Fig 8d.**
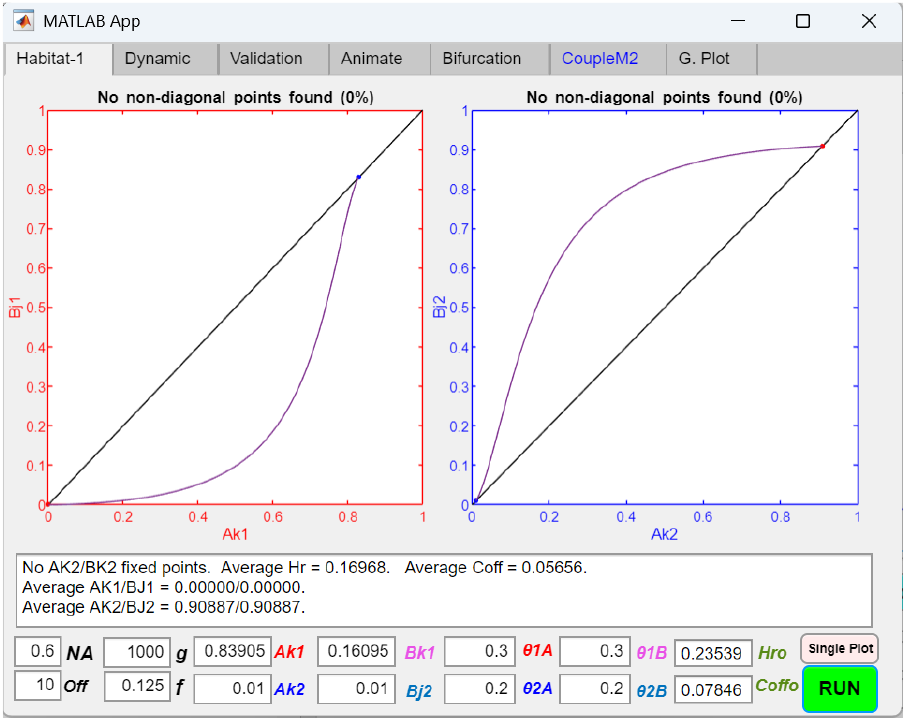
Single-plot phase portraits showing mutant alleles with stronger habitat preferences eliminating and replacing existing alleles with weaker habitat preferences in a two-allele, one-gene-locus, no-open-space model of a habitat-preference barrier (H2/L1, No), resulting in stronger RI. As illustrated in Fig 4a, two habitat-preference alleles *K*1 and *J*1 (*θ*1*A* = *θ*1*B* = 0.3) can invade and reach a fixed-point polymorphism at *Ak*1/*Bk*1 = 0.83905/0.16095 (or *Ak*1/*Aj*1 = 0.83905/0.83905), producing initial premating RI. Subsequently, if *K*1 in niche *A* mutates into a more adaptive allele *K*2 with stronger habitat preference (*θ*2*A* = 0.2) and *J*1 in niche *B* mutates into a more adaptive allele *J*2 (*θ*2*B* = 0.2), the phase portraits—initialized at the *Ak*1/*Bj*1 fixed-point values and initial mutant populations *Ak*2 = *Bj*2 = 0.01—show that *K*2 and *J*2 mutants can invade and completely replace *K*1 and *J*1 after 1000 generations. This produces a new fixed-point polymorphism at *Ak*2/*Bj*2 = 0.90887/0.90887, reduces *Hr* from 0.23539 to 0.16968 and *Coff* from 0.07846 to 0.05656, and results in stronger RI.

### Coupling Between Habitat-Preference and Mating-Bias Barriers

#### 1. Coupling between an initial mating-bias barrier and a second habitat-preference barrier

If mating bias forms the initial premating barrier under disruptive ecological selection, coupling with a subsequent habitat-preference barrier is straightforward. As long as maladaptive hybrid loss remains, a habitat-preference barrier can always invade and couple to reduce the hybrid loss to the extent allowable by its habitat-preference biases. This is true for both the one- and two-allele models as well as for the open-space and no-open-space systems.

Figs 9a–9d illustrate the coupling of a one-allele, one-gene-locus habitat-preference barrier in an open-space system following the establishment of an initial mating-bias barrier that has already created premating RI in a sympatric population under disruptive ecological selection. Because maladaptive hybrid loss persists in the system (*Hr* = 0.22141), a mutant allele *K* from the second habitat-preference barrier can invade and couple, thereby increasing overall RI. The invasion and population dynamics of the habitat-preference barrier are governed solely by the residual hybrid loss ratio (*Hr*) and proceed in the same manner as depicted in Figs 1a and 1b, as if the mating-bias barrier were not present.

Similarly, Fig 10 shows the invasion and coupling of a second two-allele, one-gene-locus habitat-preference barrier in a no-open-space system, following the establishment of an initial two-allele mating-bias barrier. Driven by the residual hybrid loss, the second habitat-preference barrier follows the same invasion and population dynamics as the two-allele habitat-preference model in a no-open-space system depicted in Figs 4a and 4b.

Fig 11a illustrates how removing disruptive ecological selection (i.e., by setting *f* = 0.5) affects the coupled barriers shown in Fig 9d. When *f* = 0.5, the *Ak*/*Bk* phase portrait of the one-allele, one-gene-locus habitat-preference barrier in an open-space system exhibits dynamics analogous to those observed in Fig 7a. If the invasion of the *K* allele is able to achieve complete RI between niches—i.e., by reducing *Hr* to zero and/or by settling at a fixed point in the stationary region of the phase portrait—the established *Ax*/*Bx* and *Ak*/*Bk* fixed points remain stable and cannot be reversed by removing disruptive ecological selection. However, for the remaining *Ak*/*Bk* population vectors that converge along the diagonal line, ongoing gene flow between niches ultimately erodes the *Ax*/*Bx* mating-bias RI and eliminates one of the mating-bias alleles.

Similarly, Fig 11b demonstrates the reversibility of the coupled barriers shown in Fig 10 when disruptive ecological selection is removed by setting *f* = 0.5. In this case, because the habitat-preference barrier is a two-allele, one-gene-locus model in a no-open-space system, complete RI between niches cannot be achieved unless the habitat-preference values, *θ*1 and *θ*2, are nearly zero. The resulting *Ak*/*Bk* phase portrait exhibits dynamics analogous to those shown in Fig 7d. Consequently, when disruptive ecological selection is removed, gene flow between niches drives the *Ax*/*Bx* phase portrait to become divergent and eliminates the *Ax*/*Bx* premating RI.

**Fig 9a.**
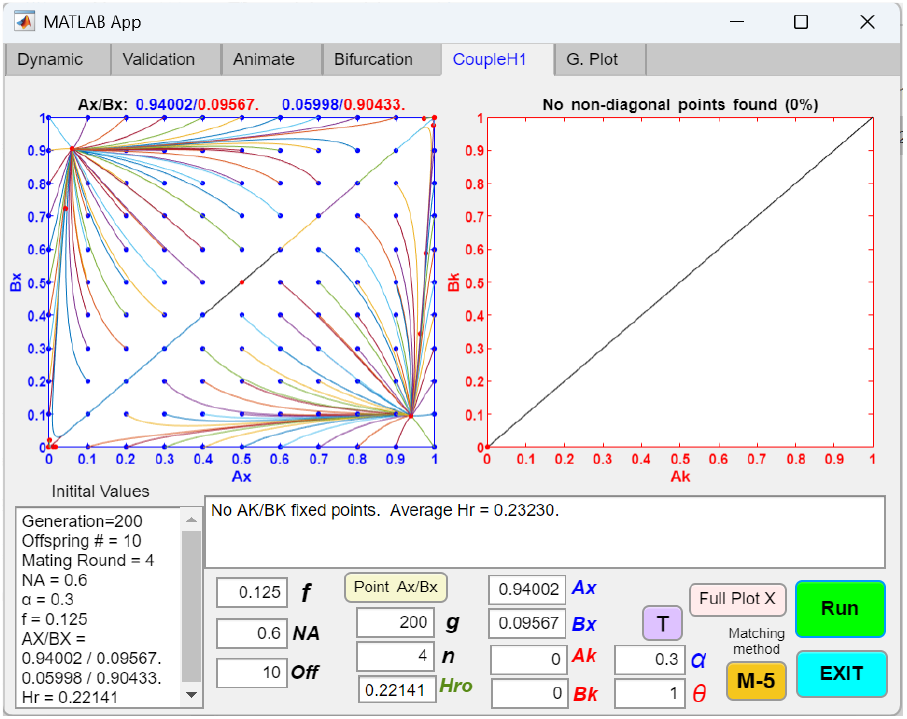
Coupling of an initial two-allele mating-bias barrier with a subsequent one-allele, one-gene-locus model of a habitat-preference barrier (M2, H1/L1, O) in an open-space system, showing the *Ax*/*Bx* phase portrait before coupling. The initial two-allele mating-bias barrier has converged to a fixed-point polymorphism at *Ax*/*Bx* = 0.94002/0.09567, given the parametric values shown in the table insert. The second one-allele habitat-preference barrier is disabled by setting *θ* = 1 and *Ak* = *Bk* = 0.

**Fig 9b.**
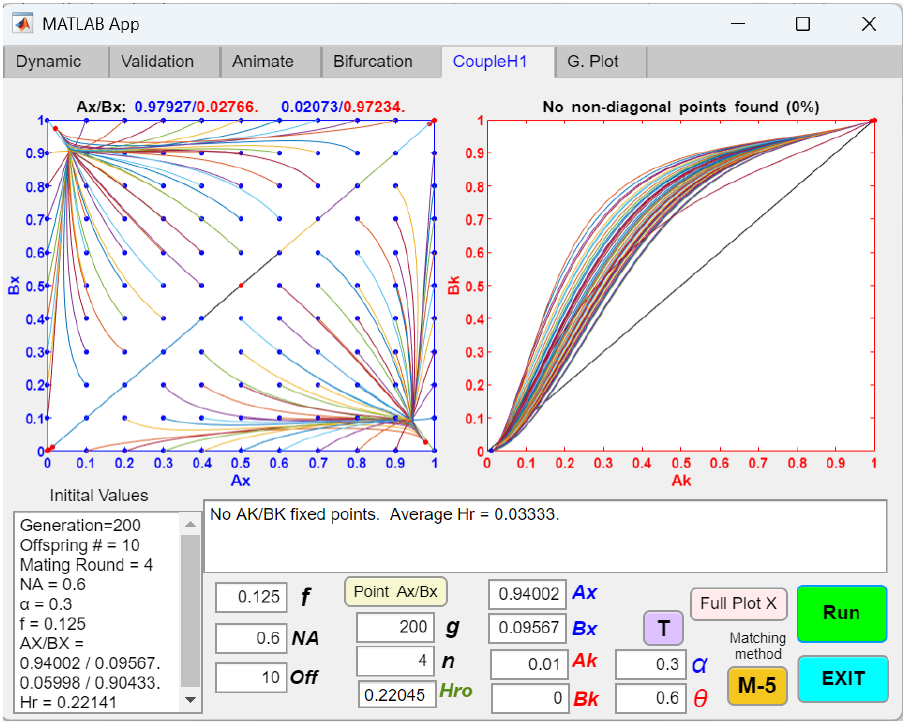
Coupling of an initial two-allele mating-bias barrier with a subsequent one-allele, one-gene-locus model of a habitat-preference barrier (M2, H1/L1, O) in an open-space system, showing the phase portraits after an invading habitat-preference allele, *K*, is fixed in both niches after coupling. In Fig 9a, a mutant habitat-preference allele, *K*, emerges in niche *A* with a habitat-preference bias of *θ* = 0.6 and an initial population ratio of *Ak* = 0.01. The *K* allele is able to invade and become fixed in both niches after 200 generations. As a result of the coupling between the two barrier systems, the *Ax*/*Bx* fixed point shifts closer to the bottom right corner of the phase portrait, moving from *Ax*/*Bx* = 0.94002/0.09567 to *Ax*/*Bx* = 0.97927/0.02766. The hybrid offspring ratio *Hr* decreases from 0.22045 to 0.03333, indicating increased overall RI.

**Fig 9c.**
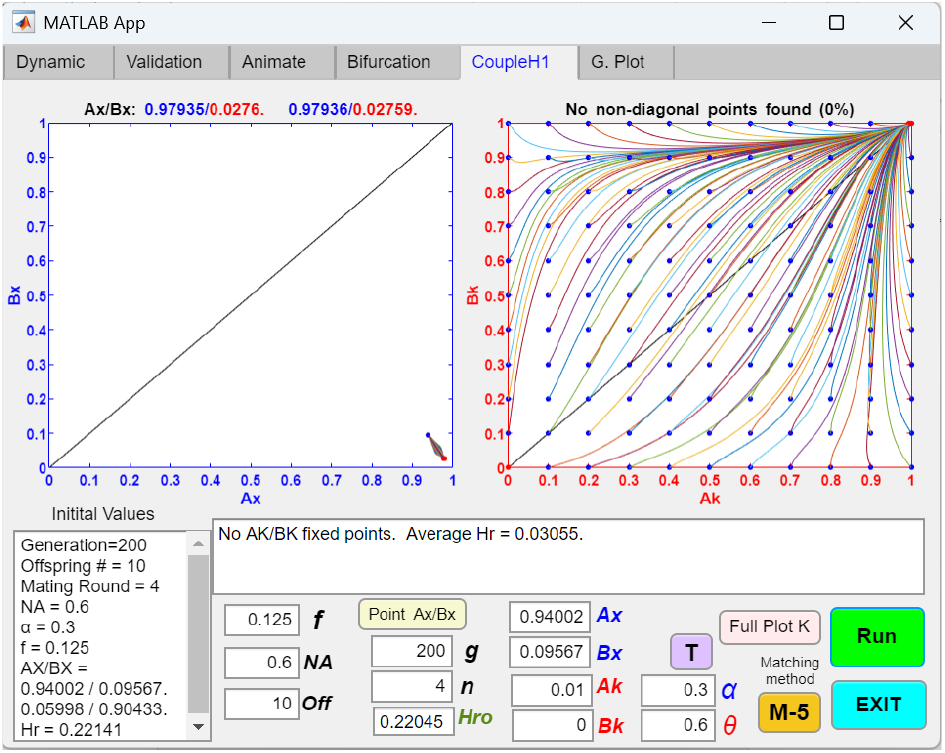
Coupling of an initial two-allele mating-bias barrier with a subsequent one-allele, one-gene-locus model of a habitat-preference barrier (M2, H1/L1, O) in an open-space system, showing the *Ak*/*Bk* phase portrait after coupling. Using the same parametric values as in Fig 9b, the phase portrait of *Ak*/*Bk*, plotted against the initial *Ax*/*Bx* fixed point values of *Ax*/*Bx* = 0.94002/0.09567, shows that all *Ak*/*Bk* population vectors become fixed at *Ak*/*Bk* = 1/1 after 200 generations.

**Fig 9d.**
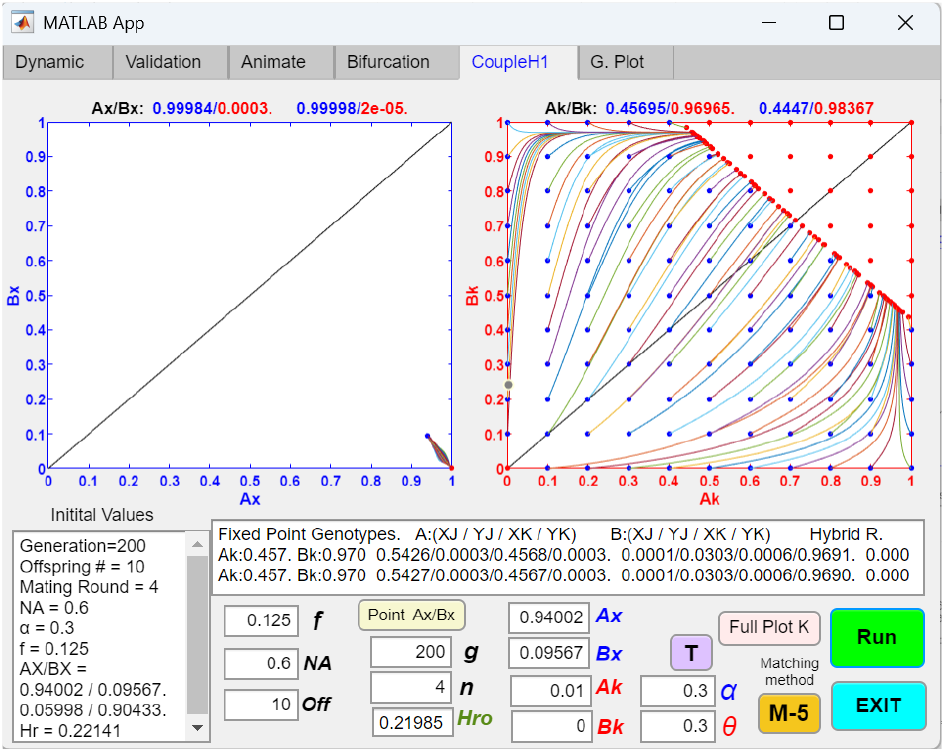
Coupling of an initial two-allele mating-bias barrier with a subsequent one-allele, one-gene-locus model of a habitat-preference barrier (M2, H1/L1, O) in an open-space system, showing stationary fixed points appearing in the *Ak*/*Bk* phase portrait when *θ* is low. If the value of *θ* in Fig 9c is decreased from 0.6 to 0.3, indicating greater habitat-preference bias, all the population vectors stop progressing once the hybrid offspring ratio, *Hr*, is reduced to zero and there is no longer any selection pressure to drive the invasion of the habitat-preference allele, *K*. This creates a region of stationary points in the upper right corner of the phase portrait, where individuals are able to find mates within their own niches and do not feel the need to venture out into open space to find mates.

**Fig 10.**
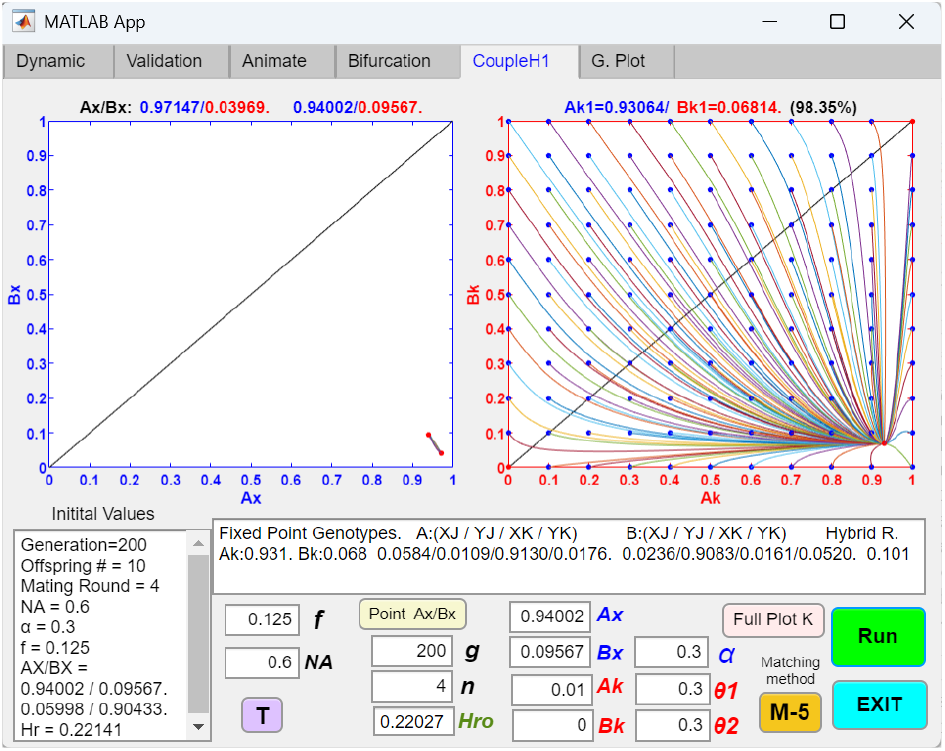
Coupling of an initial two-allele mating-bias barrier with a subsequent two-allele, one-gene-locus model of a habitat-preference barrier (M2, H2/L1, No) in a no-open-space system, showing the *Ak*/*Bk* phase portrait after coupling. The initial mating-bias barrier in Fig 9a is coupled with a subsequent two-allele habitat-preference barrier in a no-open-space system. *θ*1 = *θ*2 = 0.3. The *Ak*/*Bk* phase portrait shows a globally convergent pattern with a fixed point at *Ak*/*Bk* = 0.93064/0.06814. As a result of the coupling, the initial *Ax*/*Bx* fixed point shifts from *Ax*/*Bx* = 0.94002/0.09567 to *Ax*/*Bx* = 0.97147/0.03969, and *Hr* is reduced from 0.22027 to 0.101, resulting in stronger overall RI.

**Fig 11a.**
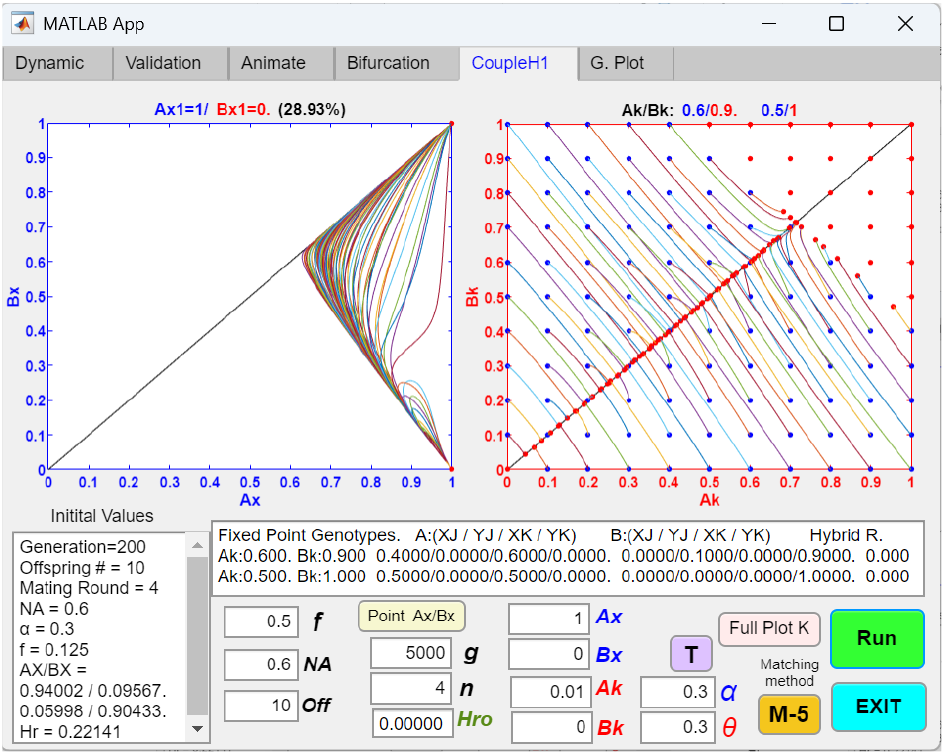
Reversibility of the coupling of an initial two-allele mating-bias barrier with a subsequent one-allele, one-gene-locus model of a habitat-preference barrier (M2, H1/L1, O) in an open-space system. Here, disruptive ecological selection for the coupled barriers in Fig 9d is removed by setting *f* = 0.5. Consequently, the *Ak*/*Bk* phase portrait exhibits a pattern similar to that shown in Fig 7a. If, after coupling, the *Ak*/*Bk* fixed point lies in the stationary region of the upper right corner of the *Ak*/*Bk* phase portrait—where *Hr* and *Coff* are zero, indicating complete reproductive isolation between the two niches—its corresponding *Ax*/*Bx* fixed point remains unchanged at *Ax*/*Bx* = 1/0 in the *Ax*/*Bx* phase portrait and cannot be reversed. For the rest of the *Ak*/*Bk* populations that are driven to the diagonal line, setting *f* = 0.5 destroys the *Ax*/*Bx* premating RI and shifts their corresponding *Ax*/*Bx* fixed points to the upper right corner of the *Ax*/*Bx* phase portrait, where *Ax*/*Bx* = 1/1 and the *Y* allele is eliminated.

**Fig 11b.**
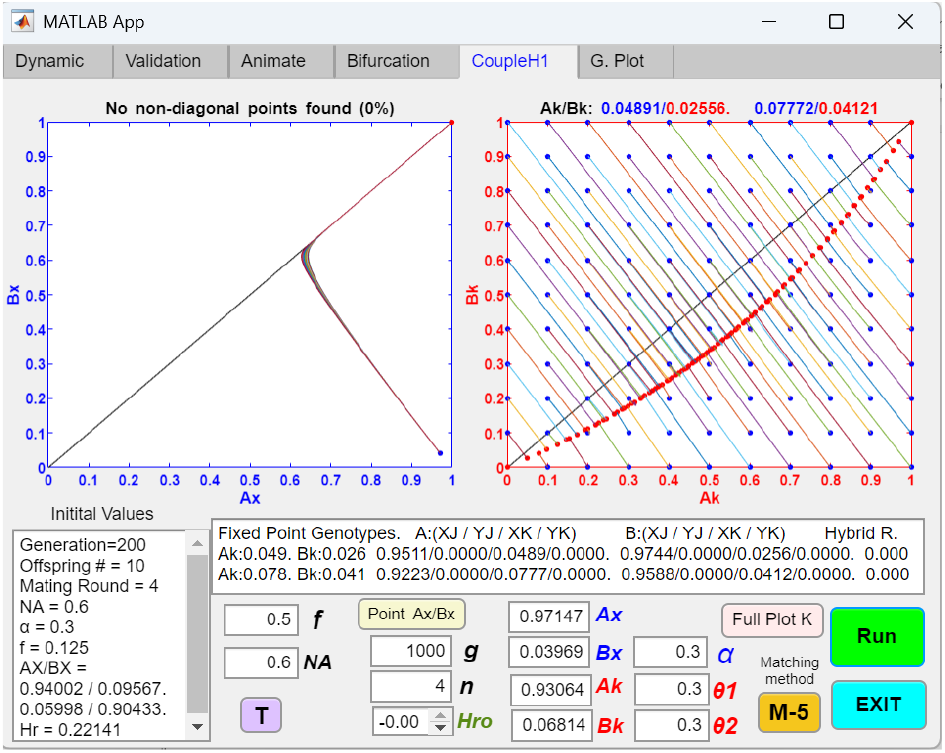
Reversibility of the coupling of an initial two-allele mating-bias barrier with a subsequent two-allele, one-gene-locus model of a habitat-preference barrier (M2, H2/L1, No) in a no-open-space system. Here, disruptive ecological selection for the coupled barriers in Fig 10 is removed by setting *f* = 0.5. The resulting *Ak*/*Bk* phase portrait resembles that shown in Fig 7d. In contrast to an open-space system, a no-open-space system lacks regions of stationary fixed points in its phase portraits. Therefore, unless RI between niches is nearly complete (i.e., when the values of *θ*1 and *θ*2 are less than 0.0025 in this example), removing disruptive ecological selection causes the *Ax*/*Bx* phase portrait to become divergent and eliminates the mating-bias RI.

#### 2. Coupling between an initial habitat-preference barrier and a second mating-bias barrier

In general, coupling a second mating-bias barrier after the establishment of an initial premating barrier in sympatric speciation is less straightforward. Unlike a second habitat-preference barrier—which can always invade and couple as long as hybrid offspring loss (*Hr*) persists in the system—a second mating-bias barrier requires more stringent conditions to successfully invade and couple. Even when *Hr* > 0, the system may remain divergent without generating mating-bias allele polymorphism or establishing premating RI, unless these optimal conditions are met. In our previous study of mating-bias barriers [21], we identified the following factors as most conducive to system convergence and the development of mating-bias RI: strong disruptive ecological selection (i.e., low values of *f*), high mating bias (low values of *α*), low assortative mating cost (achieved by a large number of mating rounds, *n*), and approximately equal niche sizes (*NA* = *NB*).

Fig 12a shows the successful coupling of an initial one-allele, one-gene-locus habitat-preference barrier with a subsequent two-allele mating-bias barrier in an open-space system. In this example, a mutant habitat-preference allele *K* with a bias of *θ* = 0.6 invades and becomes fixed in both niches, establishing initial premating RI by reducing the hybrid offspring ratio from 0.35856 to 0.07438 due to decreased inter-niche matings (see Fig 1a). Subsequently, a high-mating-bias *X* allele at a separate mating-bias barrier invades, reaches fixed-point polymorphism in the *Ax*/*Bx* phase portrait, and further reduces the hybrid offspring ratio from 0.07438 to 0.03262 (Fig 12a). Thus, the successful coupling of these two barrier mechanisms results in stronger overall RI between the two niche populations.

**Fig 12a.**
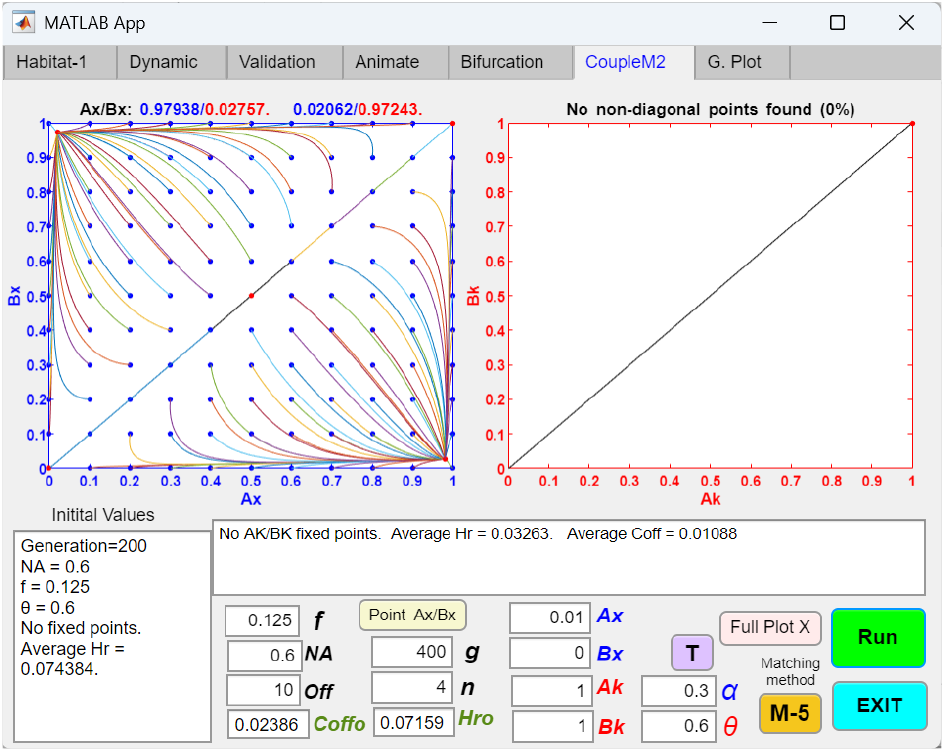
Coupling of an initial one-allele, one-gene-locus habitat-preference barrier with a subsequent two-allele mating-bias barrier (H1/L1, M2, O) in an open-space system, showing enhanced RI in the *Ax*/*Bx* phase portrait after coupling. The initial habitat-preference barrier from Fig 1a establishes the first premating RI (at *Ak*/*Bk* = 1/1, *θ* = 0.6) and is then coupled to the mating-bias barrier shown in Fig 9a. An *X* mutant allele with an initial population ratio of *Ax* = 0.01 is able to invade and reach a fixed point (*Ax*/*Bx* = 0.97938/0.02767) in the *Ax*/*Bx* phase portrait, which lies closer to the bottom right corner than the fixed point observed in the absence of the habitat-preference barrier (*Ax*/*Bx* = 0.94002/0.09567 in Fig 9a). After coupling, *Hr* decreases from 0.07159 to 0.03263, and *Coff* decreases from 0.02386 to 0.01088, indicating that the coupling has produced stronger overall RI.

Of note, in Fig 12a, the presence of the initial habitat-preference barrier results in an Ax/Bx fixed point closer to the bottom right corner of the Ax/Bx phase portrait (*Ax*/*Bx* = 0.97938/0.02757) than in the absence of the habitat-preference barrier (*Ax*/*Bx* = 0.94002/0.09567; see Fig 9a). This result indicates that a prior barrier reducing gene flow can strengthen the premating RI generated by a subsequent mating-bias barrier.

Similarly, Fig 12b shows the successful coupling of an initial two-allele, one-gene-locus habitat-preference barrier with a subsequent two-allele mating-bias barrier in a no-open-space system. After the habitat-preference barrier establishes fixed-point *Ak*/*Bk* polymorphism and initial premating RI, a high-mating-bias *X* allele invades at a separate mating-bias barrier, resulting in a shift of the *Ak*/*Bk* fixed point closer to the bottom right corner of the *Ak*/*Bk* phase portrait. This outcome suggests that the two barrier mechanisms can act synergistically to reinforce each other and generate stronger overall RI.

As shown in Fig 13a, strengthening the initial habitat-preference barrier from Fig 12a and substantially reducing gene flow prior to coupling can give rise to an invasion-resistant pattern in the *Ax*/*Bx* phase portrait of the mating-bias barrier. This occurs because reduced gene flow limits inter-niche mating, thereby weakening the fitness advantage that a high-mating-bias mutant allele would otherwise gain by producing fewer maladaptive hybrid offspring relative to its same-niche counterparts. At the same time, intra-niche sexual selection acts against the rare mutant due to its small population size and high mating bias.

Fig 13b shows that when gene flow between niches is completely stopped, the mating-bias barrier cannot couple. In this case, intra-niche sexual selection drives the predominant mating-bias allele in each niche to fixation and eliminates the less prevalent allele.

In Fig 13c, increasing the number of mating rounds (*n*) reduces the assortative mating cost and weakens intra-niche sexual selection against rare mutants, resulting in the elimination of the invasion-resistant pattern in the *Ax*/*Bx* phase portrait. Similarly, in Fig 13d, decreasing the niche size *NA* causes most individuals in niche *A* to engage in inter-niche rather than intra-niche mating. This increases the fitness advantage that a rare *Ax* mutant can derive from inter-niche matings while reducing the fitness disadvantage it incurs from intra-niche sexual selection. As a result, the *Ax* mutant may acquire a net fitness advantage to invade and couple.

**Fig 12b.**
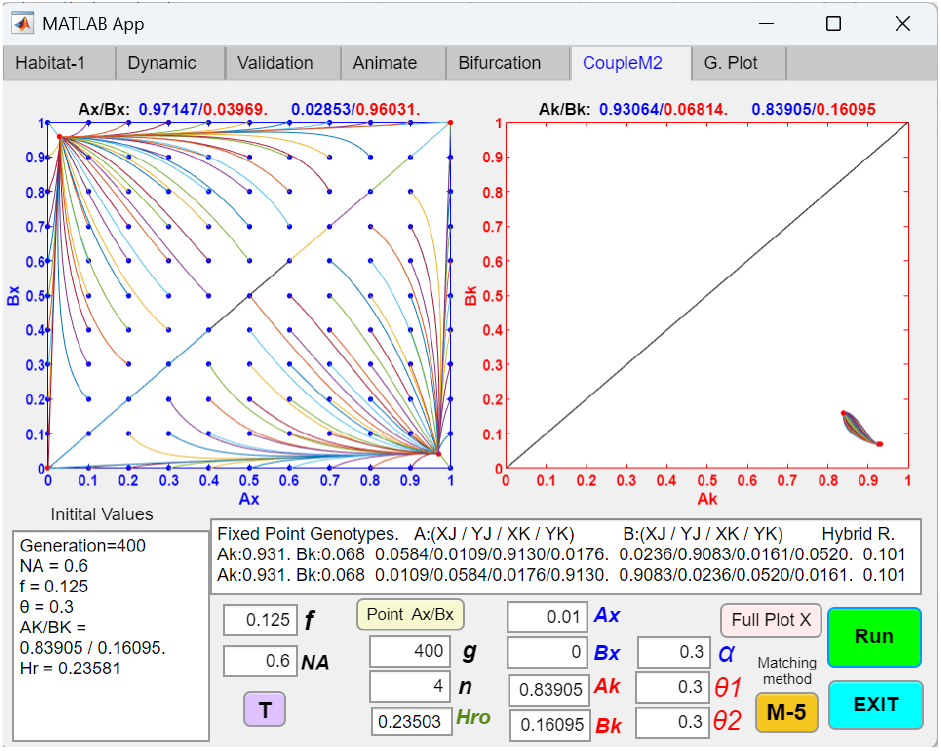
Coupling of an initial two-allele, one-gene-locus habitat-preference barrier with a subsequent two-allele mating-bias barrier (H2/L1, M2, No) in a no-open-space system, showing enhanced RI in the *Ax*/*Bx* phase portrait after coupling. The initial habitat-preference barrier from Fig 4a establishes the first premating RI (at *Ak*/*Bk* = 0.83905/0.16095, *θ*1 = *θ*2 = 0.3) and is then coupled to the mating-bias barrier shown in Fig 9a. An *X* mutant allele with an initial population ratio of *Ax* = 0.01 is able to invade and reach a fixed point (*Ax*/*Bx* = 0.97147/0.03969) in the *Ax*/*Bx* phase portrait, which lies closer to the bottom right corner than the fixed point observed in the absence of the habitat-preference barrier (*Ax*/*Bx* = 0.94002/0.09567 in Fig 9a). Meanwhile, the *Ak*/*Bk* fixed point also shifts closer to the bottom right corner of the *Ak*/*Bk* phase portrait (*Ak*/*Bk* = 0.93064/0.06814). After coupling, *Hr* decreases from 0.23503 to 0.101, indicating that the coupling has produced stronger overall RI.

**Fig 13a.**
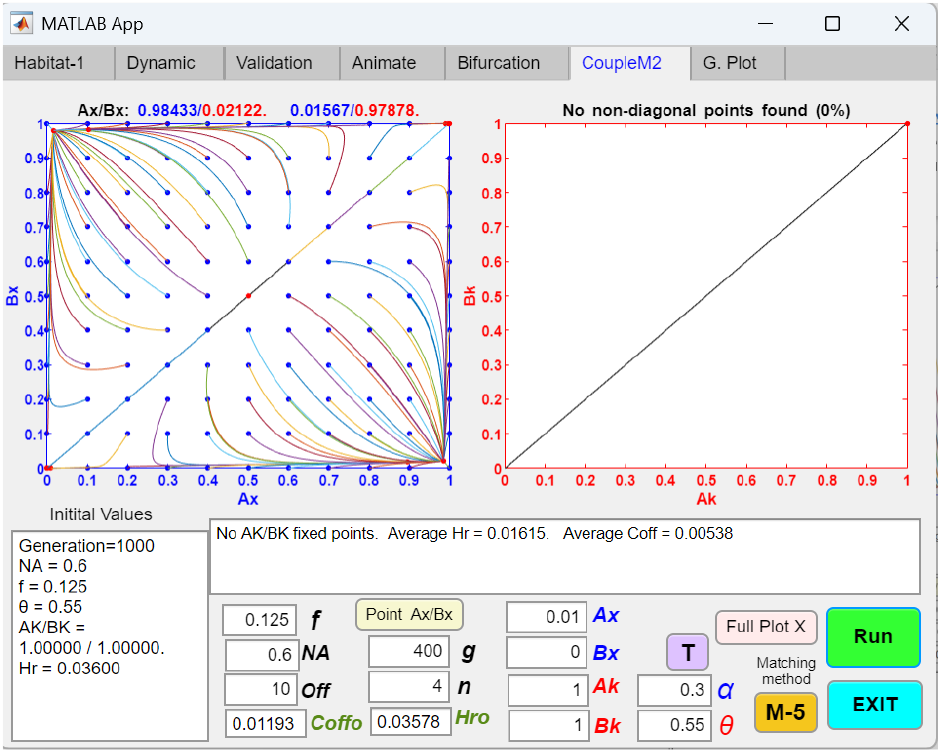
Coupling of an initial one-allele, one-gene-locus habitat-preference barrier with a subsequent two-allele mating-bias barrier (H1/L1, M2, O) in an open-space system, showing the emergence of an invasion-resistant pattern in the *Ax*/*Bx* phase portrait when gene flow is substantially reduced before coupling. Reducing the value of *θ* in Fig 12a from 0.6 to 0.55 strengthens the initial habitat-preference barrier and increases premating RI. The resulting reduction in gene flow produces an invasion-resistant pattern in the *Ax*/*Bx* phase portrait that hinders the invasion of a rare high-mating-bias *Ax* mutant and prevents successful coupling of the mating-bias barrier.

**Fig 13b.**
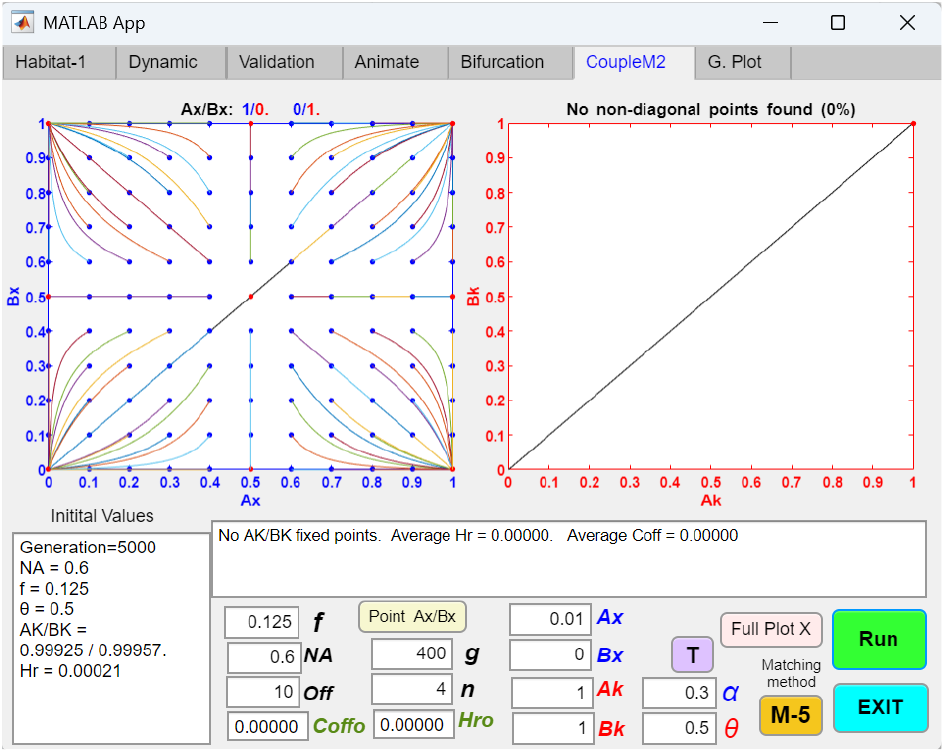
Coupling of an initial one-allele, one-gene-locus habitat-preference barrier with a subsequent two-allele mating-bias barrier (H1/L1, M2, O) in an open-space system, showing the *Ax*/*Bx* phase portrait when there is complete RI between niches. When the value of *θ* in Fig 13a is further reduced to 0.5, *Hr* = *Coff* = 0, and the habitat-preference barrier establishes complete RI. As a result, the second mating-bias barrier is unable to couple. In the *Ax*/*Bx* phase portrait, intra-niche sexual selection drives the predominant mating-bias allele in each niche to fixation and eliminates the less prevalent mating-bias allele.

**Fig 13c.**
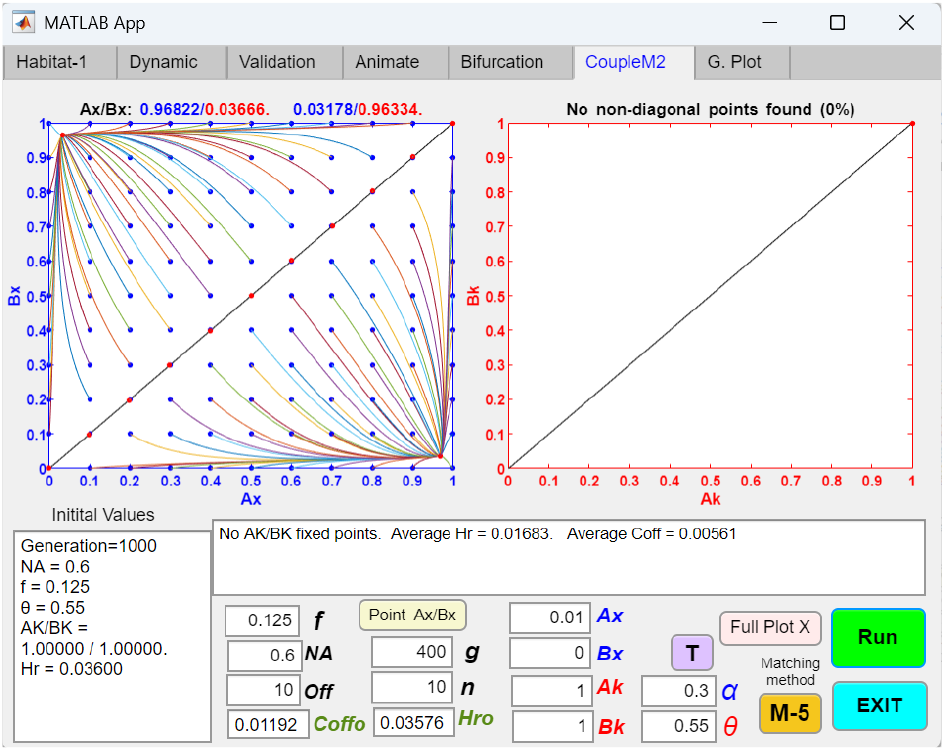
Coupling of an initial one-allele, one-gene-locus habitat-preference barrier with a subsequent two-allele mating-bias barrier (H1/L1, M2, O) in an open-space system, showing that an invasion-resistant pattern in the *Ax*/*Bx* phase portrait can be eliminated by increasing the number of matching rounds, *n*. Increasing the number of matching rounds, *n*, in Fig 13a from 4 to 10 eliminates the invasion-resistant pattern in the *Ax*/*Bx* phase portrait and allows a rare *Ax* mutant to invade and couple.

**Fig 13d.**
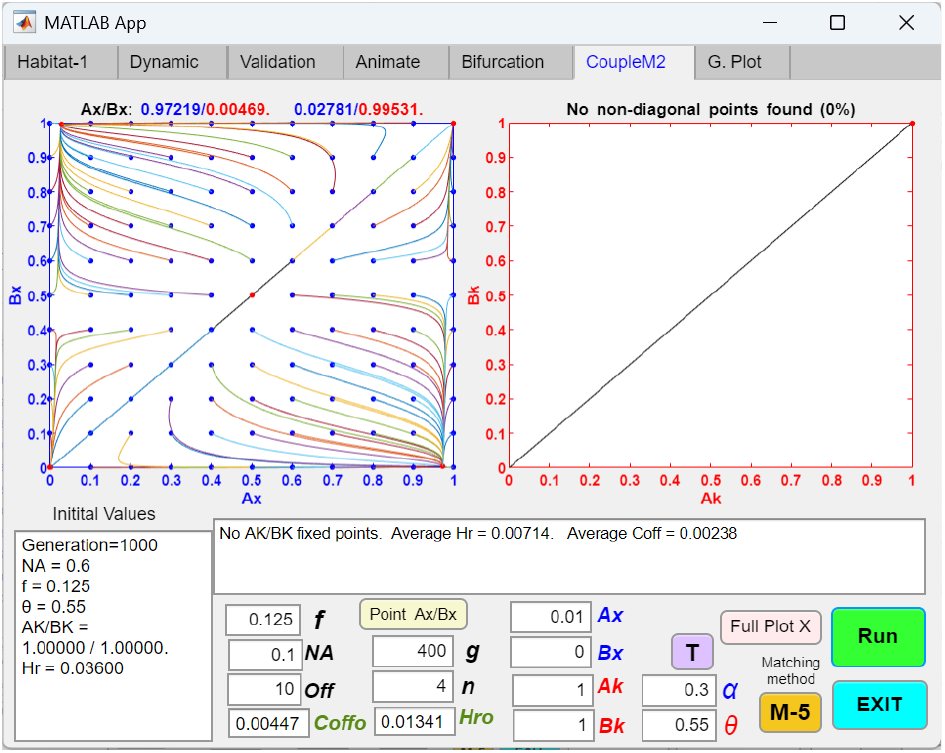
Coupling of an initial one-allele, one-gene-locus habitat-preference barrier with a subsequent two-allele mating-bias barrier (H1/L1, M2, O) in an open-space system, showing that reducing the niche size *NA* can facilitate the invasion of a rare *X* mutant allele by decreasing its invasion resistance in niche *A*. If the value of *NA* in Fig 13a is reduced from 0.6 to 0.1, the invasion-resistant pattern in Fig 13a becomes asymmetrical. A wedge-shaped region emerges along the x-axis, pointing toward the origin, where the barrier to invasion is removed. This allows a rare *Ax* mutant with an initial population ratio falling within this region to invade and establish fixed-point polymorphism. Outside this region, however, rare *Ax*/*Bx* mutant populations near the origin remain unable to invade.

Fig 14a shows an example in which the parametric values do not permit the phase portrait solution of a two-allele mating-bias barrier to converge to fixed points or establish premating RI. It also demonstrates that a weak initial habitat-preference barrier is insufficient to convert the divergent phase portrait into a convergent one or to enable the mating-bias barrier to couple.

However, in Fig 14b, when the initial habitat-preference barrier is sufficiently strong and effectively reduces gene flow between the niches prior to coupling, it can transform the divergent phase portrait of the mating-bias barrier observed in Fig 14a into a globally convergent one. This shift enables a high-mating-bias mutant allele to invade, reach fixed-point polymorphism, and establish premating RI. Thus, a robust initial barrier that limits gene flow can facilitate the coupling of a subsequent mating-bias barrier and strengthen overall RI. This is accomplished by suppressing the homogenizing effect of gene flow, which would otherwise inhibit the nonrandom assortment of mating-bias alleles between niches.

**Fig 14a.**
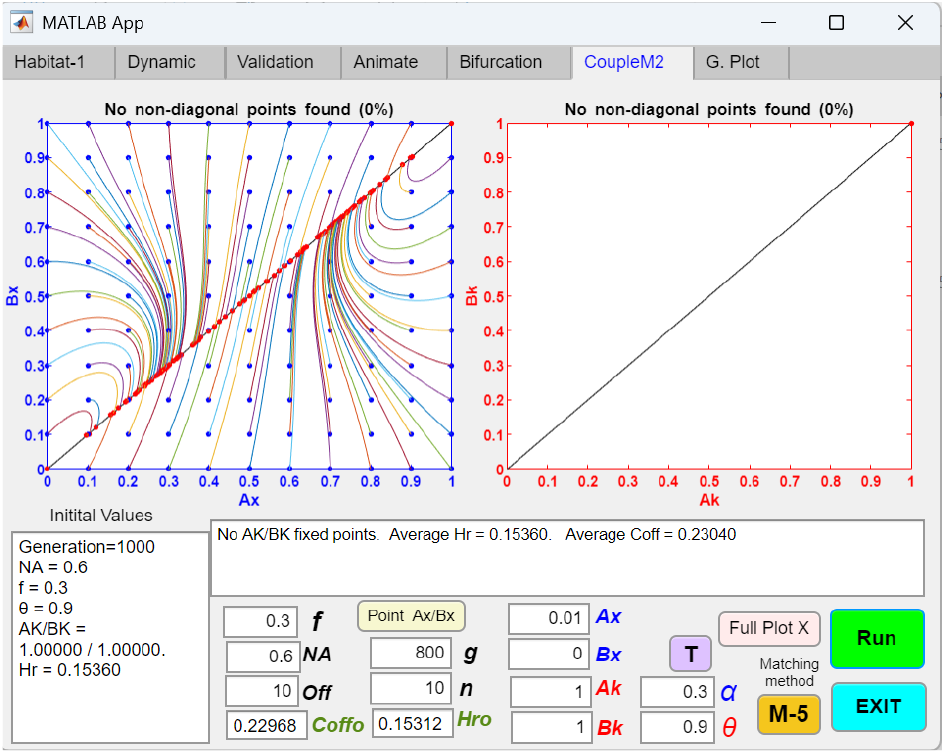
Coupling of an initial one-allele, one-gene-locus habitat-preference barrier with a subsequent two-allele mating-bias barrier (H1/L1, M2, O) in an open-space system, showing that a weak initial habitat-preference barrier is insufficient to convert a divergent mating-bias barrier into a convergent one. With the parametric values *NA* = 0.6, *f* = 0.3, *α* = 0.3, and *n* = 10, a two-allele mating-bias barrier is divergent—i.e., unable to produce fixed-point polymorphism or premating RI. Even with the establishment of a weak initial one-allele, one-gene-locus habitat-preference barrier (*θ* = 0.9), the mating-bias barrier cannot invade or couple and remain divergent.

**Fig 14b.**
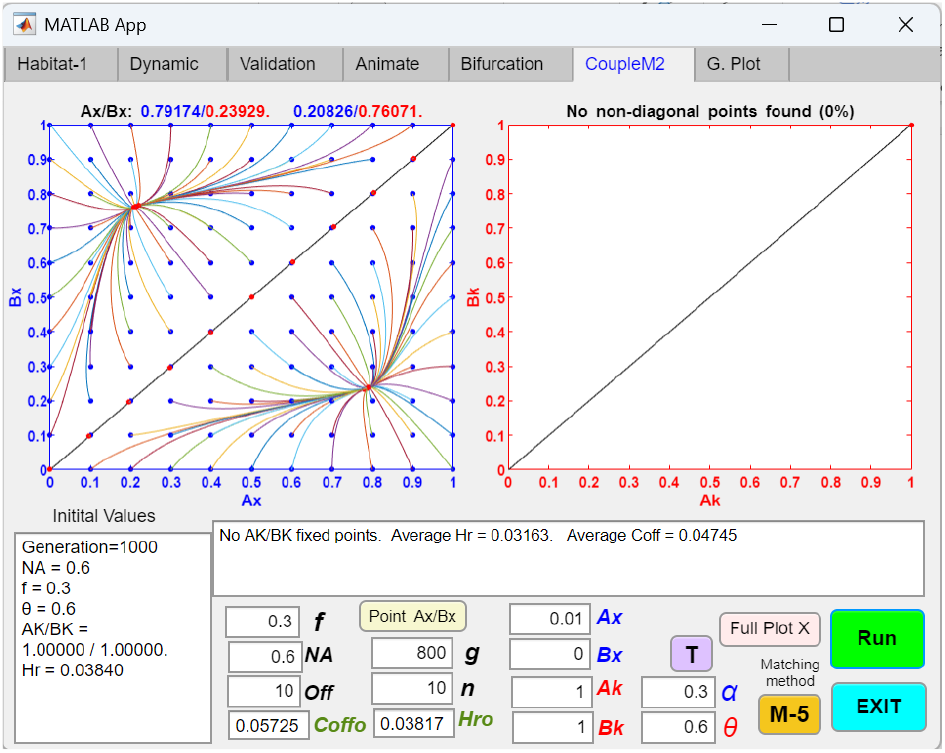
Coupling of an initial one-allele, one-gene-locus habitat-preference barrier with a subsequent two-allele mating-bias barrier (H1/L1, M2, O) in an open-space system, showing that when the initial habitat-preference barrier sufficiently reduces gene flow, it can convert a divergent mating-bias barrier into a convergent one and enable successful coupling. Reducing *θ* in Fig 14a from 0.9 to 0.6 strengthens the initial habitat-preference barrier and lowers gene flow enough to convert the divergent mating-bias barrier into a convergent one. The resulting *Ax*/*Bx* phase portrait displays a globally convergent pattern that allows a high-mating-bias mutant allele *X* in niche *A* to invade and couple. After coupling, a fixed point is established at *Ax*/*Bx* = 0.79174/0.23929, *Hr* decreases from 0.03817 to 0.03163, and *Coff* decreases from 0.05725 to 0.04745, indicating stronger overall RI.

**Fig 14c.**
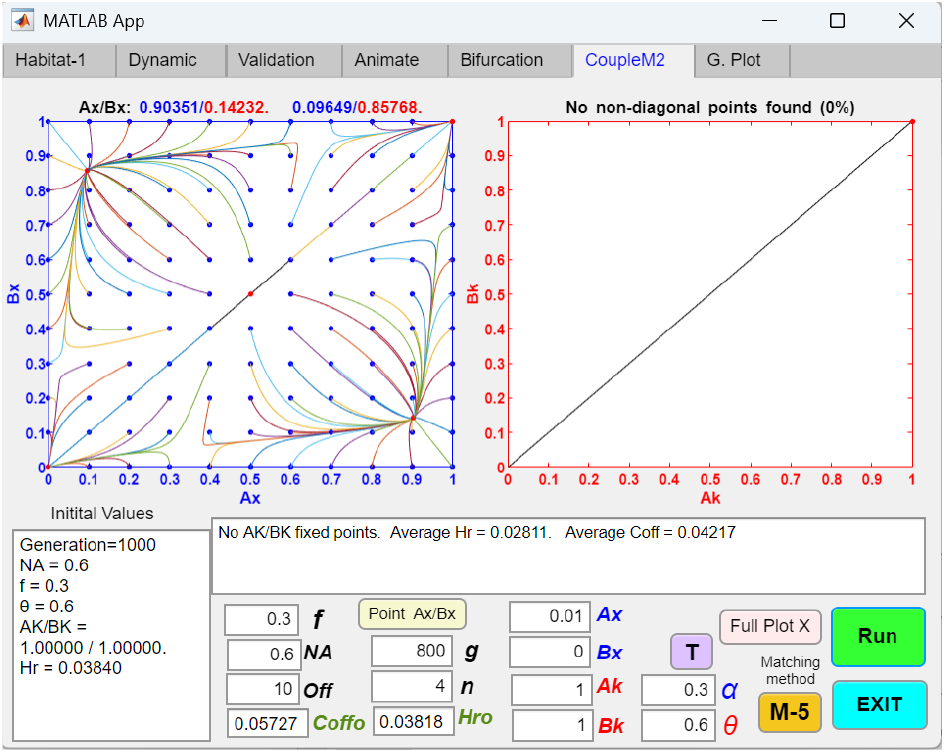
Coupling of an initial one-allele, one-gene-locus habitat-preference barrier with a subsequent two-allele mating-bias barrier (H1/L1, M2, O) in an open-space system, showing that decreasing the number of matching rounds, *n*, increases intra-niche sexual selection and creates an invasion-resistant pattern in the *Ax*/*Bx* phase portrait. Reducing the number of matching rounds, n, in Fig 14b from 10 to 4 causes an invasion-resistant pattern to emerge in the *Ax*/*Bx* phase portrait. Therefore, increasing the number of matching rounds—i.e., reducing the assortative mating cost—can mitigate intra-niche sexual selection against rare mutants and eliminate the invasion-resistant pattern.

**Fig 14d.**
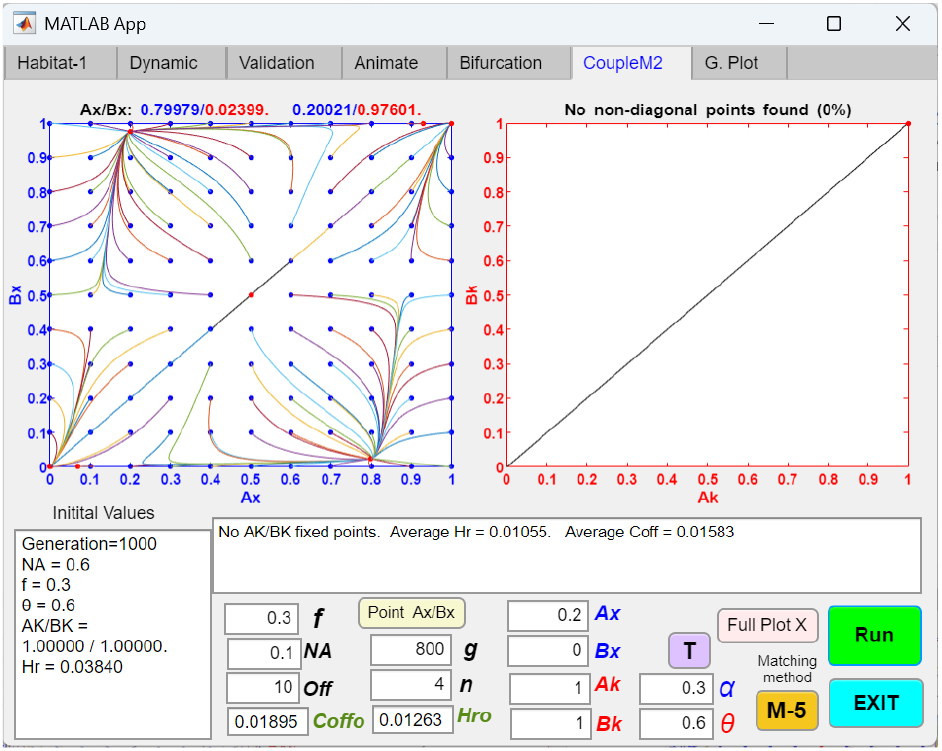
Coupling of an initial one-allele, one-gene-locus habitat-preference barrier with a subsequent two-allele mating-bias barrier (H1/L1, M2, O) in an open-space system, showing that decreasing the niche size *NA* can help a rare *Ax* mutant overcome invasion resistance and enable coupling. If the value of *NA* in 13c is reduced from 0.6 to 0.1, invasion resistance is eliminated within a wedge-shaped zone that emerges along the *x*-axis near the origin in the *Ax*/*Bx* phase portrait. As shown, an *Ax* mutant with a population ratio of *Ax* = 0.2 is able to invade, reach fixed-point polymorphism, and establish premating RI—whereas it was blocked from doing so by the invasion-resistant pattern in Fig 14c.

In Fig 14c, decreasing the number of matching rounds (*n*) increases the strength of intra-niche sexual selection in a low gene flow environment, leading to the emergence of an invasion-resistant pattern in the *Ax*/*Bx* phase portrait. As shown in Fig 14d, this invasion-resistant pattern can be mitigated by reducing the niche size *NA*, which allows a sufficiently large *Ax* mutant population to invade.

### Invasion and Coupling Dynamics of Different Habitat-Preference and Mating-Bias Barriers

Fig 15a shows the invasion generation plot of the mutant *X* mating-bias allele in Fig 12a. Following the establishment of an initial habitat-preference barrier, a small mutant population *Ax* = 0.01 in the *Ax*/*Bx* mating-bias barrier successfully invades and reaches fixed-point polymorphism at *Ax*/*Bx* = 0.97938/0.02757. Here, we define *T*_95_ as the number of generations required to reach 95% of the final *Ax*/*Bx* fixed-point values. Under the parametric conditions of Fig 12a, with *NA* = 0.6, *T*_95_ = 442 generations.

In Fig 15b, the value of *NA* in Fig 15a is reduced from 0.6 to 0.1. This change shortens *T*_95_ to 90 generations in the generation plot. In general, decreasing niche size (*NA*) shifts mating encounters for individuals in the niche from intra-niche to inter-niche. This shift increases the inter-niche mating advantage of a mutant *X* allele in the niche and reduces its intra-niche mating disadvantage, thereby facilitating its invasion.

Table 1 summarizes the invasion and coupling dynamics of the various habitat-preference and mating-bias barrier models in this study. Again, *T*_95_ is defined as the generation time required to reach 95% of either the fixation values or the steady-state fixed-point values in the phase portrait of a barrier mechanism. The table reports *T*_95_ values for the first and second barriers, as well as the combined *T*_95_ value for both. It also lists the hybrid offspring ratios (*Hr*) following the establishment of the first barrier and after coupling with the second barrier. As shown, all barrier combinations result in successful coupling except for *H*1(*L*2)_*M*2_*O*, where the initial barrier produces complete RI (*Hr* = 0), eliminating the hybrid loss needed to drive the coupling of a second barrier. Based on the results shown in Table 1, several general conclusions can be drawn regarding the mechanisms underlying barrier invasion and coupling.

**Fig 15a.**
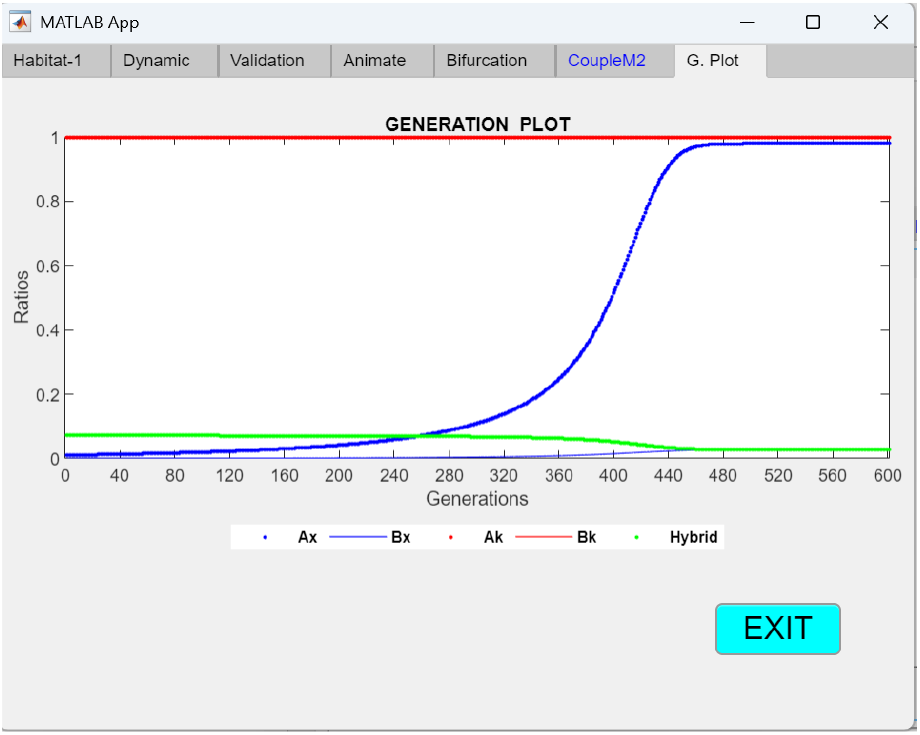
Coupling of an initial one-allele, one-gene-locus habitat-preference barrier with a subsequent two-allele mating-bias barrier (H1/L1, M2, O) in an open-space system, showing the generation plot following the invasion of a small mutant population (*Ax* = 0. 01) when *NA* = 0. 6. Using the same parametric values as in Fig 12a (*NA* = 0.6, *f* = 0.125, *α* = 0.3, *θ* = 0.6, *Ak* = *Bk* = 1), and following the establishment of an initial habitat-preference barrier, the generation plot depicts the invasion dynamics of a mutant *X* allele in niche *A* with an initial population ratio of *Ax* = 0.01. The allele reaches a stable fixed-point polymorphism at *Ax*/*Bx* = 0.97938/0.02757. The generation time required to reach 95% of the final fixed-point values (*T*_95_) is 442 generations.

**Fig 15b.**
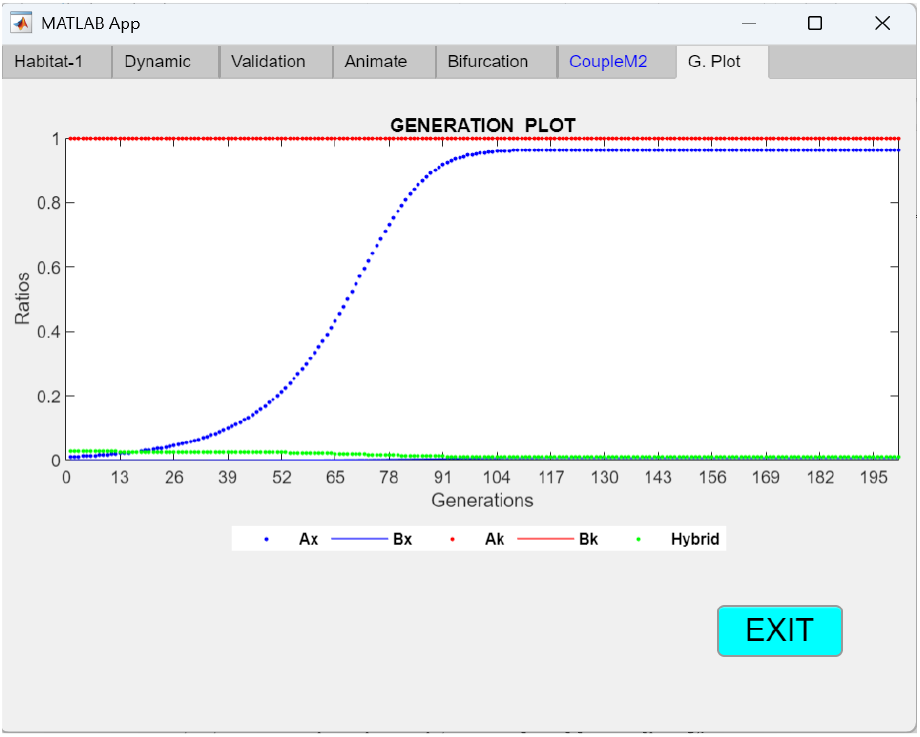
Coupling of an initial one-allele, one-gene-locus habitat-preference barrier with a subsequent two-allele mating-bias barrier (H1/L1, M2, O) in an open-space system, showing the generation plot following the invasion of a small mutant population (*Ax* = 0. 01) when *NA* = 0. 1. Decreasing the niche size *NA* in Fig 15a from *NA* = 0.6 to *NA* = 0.1 facilitates the invasion of the *X* allele in niche *A* and reduces the generation time required to reach 95% of the fixed-point values (at *Ax*/*Bx* = 0.96425/0.00638) from *T*_95_ = 442 generations to *T*_95_ = 90 generations.

**Table 1.** Comparative summary of the invasion and coupling dynamics for different combinations of habitat-preference and mating-bias barriers in open-space and no-open-space systems. The table displays *T*_95_, the generation time required to reach 95% of the fixed-point values, and *Hr*, the hybrid offspring ratio, during the coupling of different barrier mechanisms. The following parametric values are used for all simulations: *NA* = 0.6, *f* = 0.125, *α* = 0.3, *θ* = 0.6 (or *θ*1 = *θ*2 = 0.6 for two-allele models), *Ax*/*Bx* = 0.01/0, and *Ak*/*Bk* = 0.01/0 (or *Ak*/*Bk* = 0.01/0.99 for two-allele, one-locus models; *Ak*/*Bk* = 0.01/0 and *Aj*/*Bj* = 0/0.01 for two-locus models). The nomenclature for the barrier models is defined as follows: *H*1 = one-allele habitat-preference barrier, *H*2 = two-allele habitat-preference barrier, *L*1 = one-gene-locus, *L*2 = two-gene-locus, *M*2 = two-allele mating-bias barrier, *O* = open-space system, *No* = no-open-space system. The order of the barriers indicates the sequence of coupling; for example, *H*1(*L*1)_*M*2_*O* represents a one-allele, one-gene-locus habitat-preference barrier in an open-space system that is followed by the coupling of a two-allele mating-bias barrier. The final column indicates whether successful coupling between the two barriers is achieved.

| Barrier Models | $T_{95}$<br>First barrier | $Hr$<br>After first barrier | $T_{95}$<br>Second barrier | $Hr$<br>After both barriers | $T_{95}$ sum<br>Both barriers | Couple |
| --- | --- | --- | --- | --- | --- | --- |
| $H1(L1)_{M2\_O}$ | 69 | 0.072 | 442 | 0.02869 | 511 | Yes |
| $M2_{H1(L1)_{-O}}$ | 45 | 0.22141 | 127 | 0.02869 | 172 | Yes |
| $H1(L2)_{M2\_O}$ | 80 | 0 | 0 | 0 | 80 | No |
| $M2_{H1(L2)_{-O}}$ | 45 | 0.22141 | 176 | 0 | 221 | Yes |
| $H2(L1)_{M2\_O}$ | 61 | 0.12 | 150 | 0.043 | 211 | Yes |
| $M2_{H2(L1)_{-O}}$ | 45 | 0.22141 | 117 | 0.043 | 162 | Yes |
| $H2(L2)_{M2\_O}$ | 158 | 0.072 | 442 | 0.02869 | 600 | Yes |
| $M2_{H2(L2)_{-O}}$ | 45 | 0.22141 | 305 | 0.02869 | 350 | Yes |
| $H1(L1)_{M2\_No}$ | 1780 | 0.3024 | 51 | 0.16904 | 1831 | Yes |
| $M2_{H1(L1)_{-No}}$ | 45 | 0.22141 | 2338 | 0.16904 | 2383 | Yes |
| $H1(L2)_{M2\_No}$ | 1190 | 0.21254 | 71 | 0.10424 | 1261 | Yes |
| $M2_{H1(L2)_{-No}}$ | 45 | 0.22141 | 1490 | 0.10424 | 1535 | Yes |
| $H2(L1)_{M2\_No}$ | 74 | 0.339 | 46 | 0.193 | 120 | Yes |
| $M2_{H2(L1)_{-No}}$ | 45 | 0.22141 | 103 | 0.193 | 148 | Yes |
| $H2(L2)_{M2\_No}$ | 4690 | 0.3024 | 50 | 0.16963 | 4740 | Yes |
| $M2_{H2(L2)_{-No}}$ | 45 | 0.22141 | 5624 | 0.16962 | 5669 | Yes |

As an initial barrier system, a habitat-preference barrier generally invades more rapidly and establishes stronger premating RI in an open-space system than in a no-open-space system. As shown in the second and third columns of Table 1, for each type of habitat-preference barrier, the *T*_95_ value is consistently shorter—indicating faster invasion—and the resulting *Hr* value lower—indicating stronger premating RI— in open-space systems than in no-open-space systems. This difference arises because, in an open-space system, inter-niche mating occurs only when individuals leave their adaptive niches and enter the shared open space, where they may encounter and mate with individuals from the opposite niche. A habitat-preference mutation that limits such movement immediately reduces inter-niche encounters and the production of maladaptive hybrids, thereby gaining a fitness advantage to invade and spread. In contrast, in a no-open-space system, even if a niche ecotype evolves a habitat preference, individuals from the opposite niche can still enter its habitat, mate with it, and produce hybrid offspring. As a result, the fitness advantage of the habitat-preference mutation is diminished, making invasion and the establishment of premating RI more difficult.

In general, a one-allele barrier mechanism is easier to evolve than a two-allele mechanism. This is because the one-allele model is unaffected by gene flow and requires only a single mutation that can confer habitat preference in both niches. In contrast, the two-allele model requires two independently evolved mutations, each conferring preference only for a specific niche. This distinction may not be immediately apparent from the *T*_95_ values in the *H*2(*L*1) models, where a single gene locus carries two alleles (*K* and *J*), each favored in one of the two niches. However, it is important to recognize that the establishment of such a system still requires the sequential evolution of both alleles and will take longer to evolve.

Because a habitat-preference mutant allele can reach fixation in both niches in the one-allele models—rather than stabilizing at a fixed-point polymorphism as in the two-allele models—one-allele barrier mechanisms typically generate stronger premating RI than their two-allele counterparts. This is reflected in the third column of Table 1, where one-allele barriers yield lower residual *Hr* values in both open-space and no-open-space systems.

In a no-open-space system, individuals from different ecotype groups can visit one another in their respective adaptive habitats. As a result, in two-allele models, a mutation that confers habitat preference in only one habitat cannot gain a fitness advantage to invade unless a corresponding habitat preference also arises in the opposite habitat. This occurs because inter-niche mating persists: home-bound individuals in one habitat continue to encounter and mate with free-roaming individuals visiting from the other habitat, thereby negating the selective advantage of habitat preference. This phenomenon is observed in simulation runs of the *H*2(*L*2)*M*2_*No* model in Table 1. When a small mutant population (*Ak* = 0.01) with a habitat-preference bias of *θ* = 0.6 arises in niche *A*, it fails to invade if individuals in niche *B* lack habitat preference (i.e., *Bj* = 0). However, when habitat-preference mutations are present in both niches (e.g., *Ak* = *Bj* = 0.01), the two habitat-specific mutant populations can reinforce each other and establish fixed-point polymorphism and premating RI, reaching 95% of equilibrium values in approximately 4690 generations (*T*_95_ = 4690).

In contrast, this invasion barrier for single-niche habitat-preference mutations in two-allele models does not occur in open-space systems, where individuals do not visit the habitats of other niche ecotypes. In open-space systems, a niche-specific habitat-preference mutation arising in only one niche can always invade. Similarly, in the one-allele model of a no-open-space system (*H*1(*L*1)*M*2_*No*), a mutant habitat-preference allele arising in niche A (*Ak* = 0.01) can eventually seed habitat preference in niche *B* and establish RI between the two niches—although the process is much slower (*T*_95_ = 1780) compared to the open-space case (*T*_95_ = 69). If some habitat preference is already present in niche *B* as well (e.g., *Ak* = *Bk* = 0.01), *T*_95_ is reduced from 1780 to 1190. As shown in Table 1, the two-allele, one-gene-locus model (*H*2(*L*1)*M*2_*No*) in a no-open-space system can reach fixed-point polymorphism rapidly (*T*_95_ = 74), as niche-specific habitat-preference alleles are already present in the model by design (i.e., each individual carries either the *K* or *J* allele at the single gene locus).

When habitat preference acts as the first barrier and mating bias as the second barrier, the speed at which the mating-bias barrier invades and reaches equilibrium—as indicated by the *T*_95_ of the second barrier in the fourth column of Table 1—is determined by the residual hybrid offspring ratio (*Hr*) remaining after the first barrier is established (third column). The results in Table 1 show that the greater the residual *Hr*, the more rapidly the second mating-bias barrier can invade and couple.

The order in which habitat-preference and mating-bias barriers are coupled does not appear to affect the final residual *Hr* (fifth column of Table 1) or the overall level of RI achieved. However, it does influence the minimal total *T*_95_ time required to reach that final level of RI, as shown by the combined *T*_95_ values in the sixth column of Table 1.

## IV. DISCUSSION

Habitat preference is a widely recognized isolating mechanism in speciation, particularly under conditions of gene flow. By reducing mating encounters between individuals from different habitats, it is especially well suited to promote sympatric speciation, where high gene flow would otherwise hinder the nonrandom assortment of locally adaptive alleles between niche ecotypes [7, 14, 15]. When habitats are spatially separated, a habitat-preference barrier can limit mating between locally adapted ecotypes in a sympatric population under disruptive ecological selection and help establish premating reproductive isolation (RI).

Habitat-preference barriers are considered both early-stage and adaptive [35]. They are early-stage because, like mating-bias barriers, they can establish initial premating RI in sympatric speciation to jumpstart the speciation process. They are adaptive because the invasion and spread of a habitat-preference mutant allele are driven by maladaptive hybrid loss. Without hybrid loss from inter-niche mating, such a mutation cannot gain a fitness advantage over its same-niche counterparts to invade and proliferate.

In this study, we developed mathematical models to describe the one-allele and two-allele models of habitat-preference barriers in a sympatric population under disruptive ecological selection. We then used MATLAB to develop GUI applications that solve the mathematical models and display the results as phase portraits, allowing us to gain insight into the invasion and population dynamics of the habitat-preference barriers.

In the one-allele model, a single mutant allele produces the same effect across different niche ecotypes, and only the substitution of a single allele is needed to produce RI. As a result, the one-allele model is unaffected by recombination caused by gene flow. In contrast, the two-allele model requires different mutant alleles to exert effects in different ecotypes. This model is more difficult to establish under gene flow because recombination tends to break down the linkage disequilibrium between the mutant alleles and their associated ecotypes and disrupt their nonrandom assortment [12, 13].

In our study, habitat-preference barriers are modeled under two distinct spatial arrangements. The open-space system is inspired by sympatric cichlid fishes in a lake, where niche habitats are small relative to the overall lake size. In this scenario, ecotypes do not enter the habitats of other niche ecotypes. Inter-niche mating occurs only in a shared, communal space (open water) when individuals leave their adaptive habitats and encounter members of other ecotypes. In contrast, the no-open-space system is inspired by sympatric hawthorn and apple maggot flies living in an orchard, where individuals move directly between trees without lingering in midair. In this case, ecotypes can enter one another’s adaptive habitats and mate with individuals on the trees where they land. (See Methodology for details.)

Just because habitat-preference barriers can limit mating encounters between two niche ecotypes does not, by itself, imply that these populations are reproductively isolated or meet the criteria of the biological species concept [6]. They are still populations of the same species, isolated only because they are prevented from encountering one another. Consequently, coupling habitat-preference barriers with mating-bias barriers is necessary to develop true reproductive isolation based on sexual discrimination, so that premating RI can persist even when barriers to mating encounters are removed.

To explore this, we developed additional GUI applications to investigate the invasion and coupling dynamics between various habitat-preference barriers and a two-allele mating-bias barrier developed in a previous study [21]. The results provide insight into how different barrier mechanisms interact and reinforce one another during the speciation process. The key findings of our study are summarized below:

### 1. As long as hybrid loss exists, a habitat-preference barrier can always invade to reduce the hybrid loss to the extent allowable by its habitat-preference biases

In our habitat-preference models, two key variables—the offspring return ratio (*f*) and the habitat-preference bias (*θ*)—govern invasion dynamics. The variable *f* reflects the strength of disruptive ecological selection and determines the extent of hybrid loss (*Hr*) available to drive the invasion and coupling of habitat-preference barriers. The variable *θ* represents the strength of habitat preference conferred by the barrier and determines how effectively it can reduce maladaptive hybrid loss (*Hr*)

Our simulations consistently show that as long as maladaptive hybrid loss is present, a habitat-preference barrier can always invade and reduce hybrid loss to the extent permitted by its habitat-preference bias. This result holds across both one- and two-allele models, as well as in open-space and no-open-space systems.

In an open-space system, as long as *Hr* > 0, a one-allele habitat-preference mutant can always invade and move toward fixation in both niches, generating the maximum habitat-preference RI permitted by its preference bias. In the phase portrait, population vectors either converge to fixation (at *Ak* = *Bk* = 1) when the bias *θ* is insufficient to eliminate all hybrid loss (Fig 1a), or they stop progressing when *Hr* is reduced to zero and no further selection pressure remains to drive change (Fig 1b). In the latter case, a region of stationary population ratios may emerge in the phase portrait, representing a state in which ecotypes can find both food and mates within their own adaptive habitats and thus have no incentive to leave or venture into open space. At each of these stationary points, *Hr* = 0.

In contrast, a two-allele habitat-preference model tends to converge toward fixed-point polymorphism rather than fixation in the phase portrait (Fig 2a). In this case, the closer the fixed point lies to the bottom right corner of the phase portrait (*Ak*/*Bk* = 1/0), the greater the assortment of habitat-specific preference alleles between niches and the stronger the resulting RI. If hybrid loss (*Hr*) is exhausted before the system reaches a fixed point, the population vectors halt at the boundary of a region of stationary population ratios that emerges in the phase portrait (Fig 2b). Because the two-allele model converges to fixed-point polymorphism rather than fixation, the proportion of functional alleles exerting habitat-preference effects within each niche is lower. As a result, the two-allele model generally produces weaker RI than the one-allele model.

The phase portrait dynamics of the no-open-space system closely resemble those of the open-space system (Figs 3 and 4a). However, because niche ecotypes can directly visit each other’s habitats in a no-open-space system—without having to meet in a communal open space—inter-niche mating always occurs unless habitat-preference isolation is complete. As a result, hybrid loss (*Hr*) is never reduced to zero, and a region of stationary population ratios does not emerge in the phase portraits of habitat-preference barriers in the no-open-space system.

Therefore, as long as *Hr* > 0 in a no-open-space system, a one-allele habitat-preference mutant allele can always invade and become fixed in both niches to reduce hybrid loss (Fig 3). Similarly, a two-allele habitat-preference model always converges to fixed-point polymorphism in a no-open-space system (Fig 4a). At the extremes, when the biases are maximal (*θ*1 = *θ*2 = 0), the fixed point shifts to the bottom right corner of the phase portrait (*Ak*/*Bk* = 1/0), reducing *Hr* to zero. When habitat-preference biases are absent (*θ*1 = *θ*2 = 1), all population vectors converge to stationary points along the diagonal line (Fig 4b), where allele ratios are equal in both niches (*Ak* = *Bk*). Notably, vector trajectories never terminate above the diagonal line, as this would imply an increased tendency toward free-roaming or negative habitat-preference biases—i.e., the *K* allele preferring habitat *B* over *A*, and vice versa for the *J* allele—which is not allowed in our model.

### 2. Residual hybrid loss allows for the effective coupling of additional habitat-preference barriers

Our simulation results demonstrate that as long as hybrid loss remains after the establishment of a first barrier, additional habitat-preference barriers can readily couple to further reduce that loss. This applies to both the one-allele and two-allele models in the open-space system (Figs 5a and 5b), as well as to the one-allele model in the no-open-space system (Table 1). In each case, whenever hybrid loss persists, an additional habitat-preference mutant allele can always invade and reduce the remaining hybrid loss to the extent permitted by its preference bias.

A caveat applies to the two-allele, two-gene-locus model in the no-open-space system. In this case, when a habitat-specific preference mutation emerges in one habitat, it cannot invade unless a corresponding habitat-preference mutation also occurs in the opposite habitat. This constraint arises because, in a no-open-space system, ecotypes can directly visit each other’s habitats to mate. As a result, even if a habitat-specific preference mutant appears in a niche, it gains no fitness advantage over its same-niche counterparts, since individuals from the other niche can still enter its habitat and mate with it. This effect is illustrated in Figs 6a and 6b: a habitat-specific preference mutant with an initial population ratio of *Ak* = 0.01 in niche *A* cannot invade unless a corresponding mutant (*Bj* = 0.01) is also present in niche *B*. When both mutations are present, they can then reinforce each other, successfully invade, and rise to fixation in their respective niches.

This invasion barrier does not occur in the open-space system, where ecotypes do not enter each other’s habitats and all inter-niche mating takes place in a shared communal open space. As a result, a habitat-specific preference mutant that remains in its adaptive habitat gains an immediate fitness advantage over its same-niche counterparts and can successfully invade. A similar outcome occurs in the two-allele, one-gene-locus model in the no-open-space system, where both habitat-specific preference alleles (i.e., allele *K* for habitat *A* and allele *J* for habitat *B*; see Fig 4a) are already present on a single gene locus by design.

### 3. An established habitat-preference barrier can only be reversed by habitat-preference associated costs

Although habitat preference is an adaptive barrier mechanism, it is more resistant to reversal than other adaptive barriers when disruptive ecological selection is relaxed [40]. This is because, as long as maladaptive hybrid loss persists, a habitat-preference barrier can continue to invade and spread, reducing hybrid loss to the extent permitted by its associated preference bias, *θ*.

Once a habitat-preference barrier has successfully invaded or coupled, removing disruptive ecological selection by setting *f* = 0.5 does not eliminate the established habitat-preference alleles or the premating RI they generate. This is demonstrated by the examples shown in Figs 7a–7d and 10a–10b. Unlike mating-bias barriers, where sexual selection drives one of the two mating-bias alleles to extinction when disruptive selection is removed, there is no comparable intra-niche selective pressure in our model acting against habitat-preference alleles. When *f* = 0.5, hybrid loss is absent, and a habitat-preference allele no longer gains a fitness advantage through inter-niche mating compared to same-niche individuals lacking the preference. However, it also incurs no intra-niche fitness disadvantage. As a result, it maintains equal fitness with other individuals in the niche and can persist, thereby preserving the premating RI established by its presence.

In real-world systems, however, habitat-preference alleles may not remain selectively neutral in the absence of disruptive ecological selection. Costs associated with reduced dispersal—such as increased local competition for resources or missed opportunities to locate mates or novel food sources—can shift the fitness balance against these alleles. If such costs impose a fitness disadvantage on habitat-preference genotypes, they may ultimately lead to the elimination of these alleles from the population.

### 4. A more adaptive mutant tends to eliminate and replace a less adaptive resident allele in the same gene locus, whereas an adaptive mutant in a different gene locus can couple with alleles in other loci to exert additive effects

In our models, each gene locus can represent a distinct type of habitat-preference barrier. Our simulations show that when a more adaptive mutant conferring stronger habitat preference arises in such a locus, it tends to invade and displace the resident allele with weaker habitat preference. This outcome occurs in both one- and two-allele models of habitat-preference barriers (Figs 8a–8d). The principle of competitive exclusion appears to govern this process, preventing allele variants from coexisting in stable polymorphism within a single locus. As long as maladaptive hybrid loss persists, the more adaptive mutant will always gain a fitness advantage over its same-locus counterparts, leading to their displacement. Consequently, successive adaptive mutations can arise over time, each replacing its predecessor, and progressively strengthening the habitat-preference barrier conferred by that locus.

We observed the same phenomenon in our earlier studies of the two-allele model of mating-bias barriers [21, 35], where more adaptive mating-bias alleles consistently displaced less adaptive alleles within the same locus. This parallel outcome suggests a general principle across isolating barriers: adaptive alleles arising within a locus exclude less adaptive variants.

By contrast, when an adaptive mutant arises in a different locus (e.g., representing another habitat-preference barrier), the new allele can couple with alleles at existing loci to additively and synergistically reduce maladaptive hybrid loss, thereby increasing overall RI. Thus, the evolutionary impact of an adaptive mutation depends on whether it arises in the same locus—where it replaces resident alleles—or in a different locus—where it couples with existing alleles to strengthen overall RI.

### 5. Coupling a habitat-preference barrier following the establishment of an initial mating-bias barrier is facile and effective

In general, our simulation results show that coupling a habitat-preference barrier after the establishment of an initial mating-bias barrier is consistently feasible, provided that maladaptive hybrid loss (*Hr* > 0) remains in the system. This holds for both the one-allele and two-allele habitat-preference models, as well as for the open-space and no-open-space systems (Figs 9a–9d and 10). As long as hybrid loss occurs in inter-niche mating, a rare habitat-preference mutant always gains a net fitness advantage over its same-niche ecotypes lacking the preference, as there is no opposing selection pressure acting against it in intra-niche mating.

### 6. By reducing gene flow, an initial habitat-preference barrier can facilitate the coupling of a two-allele mating-bias barrier; however, very low gene flow may lead to an invasion resistance pattern in the phase portrait of the mating-bias barrier

Intuitively, in a sympatric population under disruptive ecological selection with two divergent ecological niches, a high–mating-bias mutant allele that can reduce inter-niche matings and lower the production of maladaptive hybrid offspring gains a fitness advantage and is favored to invade and spread. However, the invasion and coupling of a two-allele mating-bias barrier are less consistent than those of habitat-preference barriers. Even in the presence of maladaptive hybrid loss, a high–mating-bias mutant allele may fail to invade if optimal conditions are not met. In our previous study [21], we identified the following parameters as most conducive to the emergence of fixed-point polymorphism and premating RI in a two-allele mating-bias barrier: strong disruptive ecological selection (i.e., low values of *f*), high mating bias (low values of *α*), low assortative mating cost (achieved through a large number of mating rounds, *n*), and approximately equal niche sizes (*NA* = *NB*). The examples in Figs 12a and 12b demonstrate that when these favorable conditions are present, a two-allele model of a mating-bias barrier can be coupled to an initial habitat-preference barrier to further reduce maladaptive hybrid loss and enhance overall RI.

The primary obstacles to establishing two-allele mating-bias barriers arise from the homogenizing effect of gene flow, which prevents the differential assortment of mating-bias alleles between niches, and from intra-niche sexual selection, which acts against rare mutants with high mating biases and hinders their invasion.

As illustrated by the example in Figs 14a and 14b, when an initial habitat-preference barrier is sufficiently strong, it can facilitate the coupling of a subsequent two-allele mating-bias barrier by converting an otherwise divergent phase portrait of the mating-bias barrier into a convergent one with fixed-point polymorphism. It achieves this by reducing gene flow and, consequently, its homogenizing effect, which tends to inhibit the nonrandom assortment of mating-bias alleles between niches. This finding is supported by previous studies [7, 15].

Mechanistically, this outcome arises in our simulation because offspring genotypes are formed by random assortment of parental alleles. When a high– mating-bias mutant allele *X* arises in niche *A* (genotype *Ax*) and attempts to invade a population under disruptive ecological selection composed only of individuals carrying the *Y* mating-bias allele (i.e., *Ay* and *By* genotypes), inter-niche matings between *Ax* and *By* individuals produce offspring in equal proportions of *Ax, Ay, Bx*, and *By*, due to random assortment of parental alleles. This reduces the offspring return ratio of an *Ax* parent by half, as only 50% of its niche-*A* offspring share its genotype (*Ax*). By contrast, for most individuals in niche *A* carrying the *Ay* genotype, inter-niche matings with *By* genotypes yield 100% of the niche-*A* offspring that share the same genotype (*Ay*) as their parents. Consequently, the *Ax* mutant suffers a relative fitness disadvantage in inter-niche mating compared to its same-niche counterparts with the *Ay* genotype.

This effect is illustrated in Fig 14a: when the initial *Ax* population is small, the *Bx* ratio increases more rapidly than the *Ax* ratio, and the population vectors converge toward the diagonal line rather than a fixed point. Because gene flow transfers equal proportions of the *X* allele to the *B* ecotype and the *Y* allele to the *A* ecotype, it acts to homogenize the *X* and *Y* allele ratios between the niches and prevent differential assortment. A habitat-preference barrier reduces this homogenizing effect by limiting gene flow, thereby diminishing the relative fitness disadvantage of the *Ax* mutant in inter-niche mating due to gene flow while leaving the fitness of the *Ay* genotypes unaffected. As a result, the net invasion fitness of the *Ax* mutant increases, facilitating the evolution and coupling of the mating-bias barrier.

However, when the initial habitat-preference barrier further restricts gene flow between the niche ecotypes beyond an optimal range, an invasion-resistant pattern appears in the phase portrait of the mating-bias barrier (Fig 14c). This occurs because, aside from the inter-niche disadvantage caused by the homogenizing effect of gene flow, the invasion fitness of a high–mating-bias mutant allele in a niche is shaped by the balance between two opposing forces. The first is the relative fitness advantage the mutant can gain in inter-niche mating due to reduced production of maladaptive hybrid offspring compared to its same-niche counterparts. The second is the fitness disadvantage it can incur in intra-niche mating due to sexual selection acting against rare, incompatible mutants.

Because the *Ax* genotype has a higher mating bias (low *α*), it experiences reduced inter-niche mating success when encountering the predominantly *By* genotype from niche *B*, compared to its same-niche *Ay* counterparts. This premating mating-bias barrier simultaneously decreases the inter-niche disadvantage of the *Ax* mutant caused by the homogenizing effect of gene flow and increases its inter-niche advantage from reduced maladaptive hybrid loss, relative to the *Ay* genotype in the same niche. As gene flow is further reduced by a habitat-preference barrier, most matings shift from inter-niche to intra-niche, diminishing the relative fitness advantage that the mutant derives from inter-niche mating. At the same time, increased intra-niche mating makes sexual selection within the niche more likely to eliminate small populations of high– mating-bias mutants and to cause population ratios near the origin in the phase portrait to collapse back toward the origin. These dynamics produce an invasion-resistant pattern in the *Ax*/*Bx* phase portrait.

This invasion-resistant pattern can be eliminated by increasing the number of mating rounds (*n*) or by decreasing the niche size (*NA*) in the model. Increasing the number of mating rounds reduces the assortative mating cost and weakens intra-niche sexual selection against rare high-mating-bias mutants. As a result, the invasion-resistant pattern disappears, and the phase portrait becomes globally convergent, as illustrated in Figs 14b and 14c.

Alternatively, reducing the relative size of the niche (*NA*) in which the high-mating-bias mutant (*Ax*) arises shifts most matings for individuals in the niche from intra-niche to inter-niche. This can restore the mutant’s fitness advantage from inter-niche mating and give it a net fitness advantage to invade. In the phase portrait, this is manifested by the appearance of a wedge-shaped zone along the *x*-axis near the origin, where invasion resistance is eliminated and rare mutants are able to invade (Fig 14d). However, notice that the presence of compatible mating-bias mutants in the opposite niche (e.g., *Bx* > 0) can erode the inter-niche fitness advantage of the invading mutant (*Ax*) by increasing its inter-niche mating success, since no mating bias exists between *Ax* and *Bx* genotypes.

In general, under disruptive ecological selection, a smaller niche size (*NA*) increases the fitness advantage that a habitat-preference or mating-bias mutant allele (*Ak* or *Ax*) can gain from inter-niche mating due to reduced maladaptive hybrid loss, thereby facilitating its invasion in the smaller niche. In the invasion generation plot, this is reflected by a shorter *T*_95_—the number of generations required for the mutant population to reach 95% of its final steady-state values—as niche size decreases (Figs 15a and 15b). In the two-allele model of mating-bias barriers, although reducing niche size (*NA*) allows the mutant allele (*Ax*) to invade more rapidly, the resulting fixed point is often located farther from the bottom right corner of the phase portrait than when the niche sizes are approximately equal (i.e., when *NA* = *NB* = 0.5).

Lastly, strong selection against hybrids (i.e., a low value of *f*), whether due to extrinsic disruptive ecological selection (ecologically maladaptive hybrids) or intrinsic hybrid incompatibility (due to postzygotic barriers), can increase the relative inter-niche mating advantage of the high-mating-bias mutant and thereby facilitate its invasion and spread. In Fig 14c, reducing *f* from 0.3 to 0.125 eliminates the invasion-resistant pattern in the *Ax*/*Bx* phase portrait.

In Fig 14c, increasing the mating bias of the *X* mutant allele by reducing *α* from 0.3 toward 0 appears to have little effect on the size of the invasion-resistant region. This likely occurs because, although a greater mating bias enhances the mutant’s inter-niche mating advantage, it simultaneously intensifies intra-niche sexual selection against the rare mutant population, offsetting the overall benefit. On the other hand, increasing *α* to 0.4 makes the *Ax*/*Bx* phase portrait globally divergent again, with no fixed points remaining.

Eventually, when the initial habitat-preference barrier reduces gene flow to near zero, intra-niche sexual selection becomes too strong to overcome, and the *Ax*/*Bx* phase portrait pattern shown in Fig 13b emerges, indicating complete RI between the niches. As a result, the less prevalent mating-bias allele in each niche is driven to extinction, and the mating-bias barrier cannot invade or couple.

### 7. Comparison of invasion and coupling dynamics between habitat-preference and mating-bias barriers in open-space and no-open-space systems

Table 1 summarizes the invasion and coupling dynamics of the various barrier models examined in this study. Overall, the results confirm that both habitat preference and mating bias function as adaptive barrier mechanisms, meaning their invasion and coupling depend on the presence of maladaptive hybrid loss (*Hr*). The greater the hybrid loss, the stronger the selection pressure driving their invasion and coupling, as reflected by shorter *T*_95_ times.

The one-allele barrier model is easier to evolve and tends to produce stronger RI than the two-allele model. This is because it is unaffected by the homogenizing effect of gene flow and requires only the evolution of a single mutation that is effective in both niches. In the one-allele model, a habitat-preference mutant allele can reach fixation in both niches, rather than stabilizing as a fixed-point polymorphism with lower effective allele frequencies in each niche, as is the case in the two-allele model. As a result, the one-allele model can generate stronger premating RI and leaves a smaller residual hybrid loss (*Hr*) following establishment compared to its two-allele counterpart.

Habitat-preference barriers are easier to establish and tend to produce stronger RI in the open-space system than in the no-open-space system. This difference arises because, in the open-space system, niche ecotypes do not enter each other’s adaptive habitats, and inter-niche mating occurs only in a shared communal space. A habitat-preference mutation that prevents individuals from leaving their adaptive habitat to mate in the open space with foreign ecotypes—and thus avoids producing maladaptive hybrid offspring—immediately gains a fitness advantage to invade and spread.

In contrast, in the no-open-space system, ecotypes are free to enter each other’s adaptive habitats. As a result, a habitat-preference mutation gains less fitness advantage from avoiding inter-niche mating, since foreign ecotypes can still enter its habitat and mate with it. Moreover, in the no-open-space system, the habitat-preference barrier cannot completely eliminate hybrid offspring loss (*Hr*) unless preference is absolute (i.e., *θ* = 0). This limits the maximum RI that can be achieved compared to the open-space system. Consequently, the residual hybrid offspring loss (reflected by the value of *Hr*) following the establishment of a habitat-preference barrier remains higher in the no-open-space system than in its open-space counterpart.

Our simulation results show that changing the order in which habitat-preference and mating-bias barriers are coupled does not affect the final residual hybrid offspring loss (*Hr*) or the overall RI achieved, as each barrier independently reduces hybrid loss to the extent permitted by its mechanism. However, the order of coupling does influence the minimum generation time required to reach the final RI (as reflected by the sum of *T*_95_ values in column 6 of Table 1). This variation arises because the *T*_95_ of the second barrier depends on the residual *Hr* remaining after the first barrier is established. A low residual *Hr* value reduces the selection pressure available to drive the invasion and coupling of the second barrier, thereby increasing its *T*_95_.

## V. LIMITATIONS OF THE STUDY

All of our barrier models assume no viable hybrids, with the strength of disruptive ecological selection represented by the offspring return ratio, *f*. If viable hybrids are present, they would effectively increase *f* and weaken ecological selection. In this way, *f* can still be used to model the impact of hybrid viability. Using a single variable *f* greatly simplifies computation and helps clarify the invasion and coupling dynamics of various barrier mechanisms, and it should not compromise the core validity of our findings. In a separate study, we examine the effects of incorporating viable ecological and mating-bias hybrids into the barrier models [41].

In our models, individuals are assumed to be haploid hermaphrodites. Theoretically, the results should remain the same if gonochoric individuals (with separate male and female sexes) are used. This is because natural systems tend to maintain equal sex ratios through negative feedback mechanisms, leading parents to produce equal numbers of male and female offspring with similar niche allele compositions. Since individuals of the same sex cannot mate and are effectively invisible to one another in the mating pool, the calculations should ultimately yield equivalent results. This prediction appears to be supported by previous modeling studies [42, 43], although we have not explicitly implemented or tested it in the present study.

In our study, genetic inheritance from parents to offspring is modeled through the random assortment of alleles at each gene locus. This approach is biologically realistic when the relevant genes are located on different chromosomes. However, if the genes are located on the same chromosome or in regions with variable recombination rates, specific recombination ratios must be incorporated into the model to accurately represent allele inheritance [44].

Similarly, for the open-space and no-open-space systems, we assume two behavioral extremes. In the open-space system, habitats are small, and ecotypes do not visit each other’s habitats. In the no-open-space system, ecotypes randomly visit both habitats, even though resources are limited in the incompatible habitat. In reality, encounter behavior likely falls somewhere between these two extremes—and so do the outcomes of our results.

Our mathematical algorithm is designed to ensure a “perfect match,” meaning that every individual successfully encounters a partner in each matching round, with no one left unpaired—subject to spatial constraints (open-space or no-open-space systems) and individual behavior (i.e., whether ecotypes visit one another’s habitats). This optimization can lead to unusual patterns in the phase portrait, such as regions of stationary population ratios, as shown in Figs 1b and 2b. In these regions, all individuals can find mates within their own niches and thus have no incentive to venture into the communal open space, where there is neither food nor anyone else to meet. In real-world scenarios—take cichlid fish as an example—it is unlikely that individuals would exhibit such perfectly optimized behavior and remain confined to one area. Given the opportunity, they are more likely to roam freely throughout the lake and explore—because that is what fish do.

## VI. CONCLUSION

Habitat preference is an important barrier mechanism in early-stage sympatric speciation. In a sympatric population under disruptive ecological selection, a mutation that confers habitat preference and causes ecotypes to remain in their adaptive habitats can immediately gain a fitness advantage by reducing maladaptive hybrid loss from inter-niche mating, enabling it to invade and proliferate [3, 7, 15].

In this study, we developed mathematical models and used computer simulations to investigate the properties of different habitat-preference barriers, as well as their invasion and coupling dynamics with a two-allele mating-bias barrier model developed in a previous study [21]. We modeled the habitat-preference barriers under two spatial configurations: an open-space system, in which ecotypes do not visit each other’s habitats and encounters occur only in a communal open space; and a no-open-space system, where ecotypes can visit each other’s habitats directly.

Our results confirm the role of habitat-preference barriers as an early-stage, adaptive barrier mechanism in sympatric speciation [35]. They are considered early-stage because habitat preference can serve as the initial barrier that limits gene flow and facilitates the emergence and coupling of late-stage barriers such as chromosomal inversions, mutations reducing hybrid viability, and Bateson–Dobzhansky–Muller (BDM) incompatibilities [35, 45, 46]. They are considered adaptive because their invasion and establishment are driven by selection pressures resulting from maladaptive hybrid loss.

In a sympatric population under disruptive ecological selection, as long as maladaptive hybrid loss exists, a habitat-preference barrier can always invade to reduce hybrid loss to the extent permitted by its habitat-preference biases. In general, a one-allele model of habitat-preference barriers is easier to evolve and produces stronger reproductive isolation (RI) than a two-allele model because a one-allele model is not affected by the homogenizing effects of gene flow.

The spatial arrangements of habitats affect the invasion and coupling dynamics of habitat-preference barriers. These barriers evolve more readily and tend to produce stronger RI in open-space systems than in no-open-space systems. In open-space systems, ecotypes cannot enter each other’s habitats, and all encounters occur in a communal space, giving a mutation that confers habitat preference an inherent fitness advantage by reducing inter-niche matings. In contrast, in no-open-space systems, where ecotypes can visit each other’s habitats, this advantage is weakened by continued encounters with ecotypes from the opposite niche.

Thus, the degree of spatial separation and contiguity among habitats within a sympatric environment appears to influence how readily different barrier mechanisms become established. When distinct habitats are few and well-separated, habitat-preference barriers are more likely to evolve and initiate premating RI between niche ecotypes. By contrast, when small habitats are intermingled in a contiguous mosaic pattern, mating-bias barriers may offer a more feasible route to initiating premating isolation, as limited separation hinders the formation of spatially distinct habitats and the restriction of inter-niche encounters [25].

Once established, a habitat-preference barrier appears to exhibit a degree of resistance to reversal, even when disruptive ecological selection is removed. This persistence arises because, unlike mating-bias alleles [47], habitat-preference alleles are not subject to negative intra-niche selection, even after their inter-niche fitness advantage disappears. As a result, they can only be eliminated through costs associated with habitat preference, such as increased local competition or missed opportunities to explore and acquire new resources outside the habitat.

With both habitat-preference and mating-bias barriers, adaptive mutants can arise repeatedly within a gene locus and displace less adaptive variants, leading to progressively stronger barrier effects. The principle of competitive exclusion prevents allele variants from coexisting in stable polymorphism within a single gene locus.

After the establishment of an initial barrier, as long as maladaptive hybrid loss remains, additional habitat-preference barriers can always couple to reduce that loss, to the extent allowed by their preference biases. This contrasts with the two-allele mating-bias barrier, which can invade and couple only under specific favorable parametric conditions.

In a sympatric population under disruptive ecological selection, a new mutation can invade and spread within a niche only if it gains a net fitness advantage over resident individuals lacking the mutation. In our two-allele mating-bias barrier model, the invasion fitness of a high-mating-bias mutant allele relative to its same-niche counterparts is determined by three main factors: (1) the inter-niche mating disadvantage caused by the homogenizing effects of gene flow, (2) the inter-niche mating advantage resulting from reduced maladaptive hybrid offspring loss, and (3) the intra-niche mating disadvantage arising from sexual selection against rare mutant populations. The sum of these effects must be positive for the mating-bias mutant to successfully invade and spread, whereas a negative sum results in extinction. By contrast, factor (3) does not apply to habitat-preference mutant alleles, as there is no intra-niche selection against rare habitat-preference types. The absence of such intra-niche selection gives habitat-preference mutants a higher net invasion fitness and greater resistance to reversal.

When habitat preference serves as the initial barrier, it can facilitate the coupling of a subsequent two-allele mating-bias barrier. It does so by reducing inter-niche matings and mitigating the homogenizing effects of gene flow, which would otherwise prevent the non-random assortment of mating-bias alleles [7, 15].

However, as gene flow becomes significantly reduced, an invasion-resistant pattern may emerge in the phase portrait of the mating-bias barrier. This occurs because reduced gene flow shifts most matings from inter-niche to intra-niche, diminishing the fitness advantage that a high-mating-bias mutant would otherwise gain from reduced inter-niche mating relative to its same-niche counterparts. At the same time, increased intra-niche mating intensifies sexual selection against the rare high-mating-bias mutant, further hindering its invasion. This invasion resistance can often be overcome by increasing the number of matching rounds or decreasing niche size, both of which reduce intra-niche sexual selection against the minority mutant population. Strengthening selection against hybrids can also help reduce the invasion-resistant pattern by enhancing the mutant’s inter-niche mating advantage. Alternatively, if the invading mutant can attain a sufficiently large initial population—such as might occur through drift or in a secondary contact scenario—it may escape the invasion-resistant zone near the origin of the phase portrait, thereby overcoming the early selective disadvantage imposed by sexual selection against rare genotypes. Consequently, the mutant population can successfully invade, reach a fixed-point polymorphism, and establish premating RI. Finally, as premating RI continues to strengthen, gene flow may be reduced to such an extent that too little hybrid loss remains to drive the invasion and coupling of additional mating-bias barriers.

A habitat-preference mutation can function as a “magic trait” by simultaneously contributing to ecological adaptation and premating isolation [48, 49]. In contrast, mating-bias traits are not inherently associated with ecological adaptations, making nonrandom assortment of mating-bias alleles more difficult to achieve [1].

The relative ease and effectiveness of habitat-preference barriers make them strong candidates for establishing initial premating RI in sympatric speciation. Once established, these barriers can also facilitate the invasion and coupling of additional mating-bias barriers, further strengthening overall RI. Given ongoing criticisms that stringent conditions are often required for mating-bias barriers to initiate premating RI in sympatric populations, a “habitat-preference-first, mating-bias-second” pathway offers a compelling alternative to address these concerns and initiate the speciation process.

In conclusion, our results reaffirm habitat preference as an important barrier mechanism in sympatric speciation. We have described key properties of various habitat-preference barriers across different spatial arrangements, as well as their invasion and coupling dynamics with a two-allele mating-bias barrier, subject to the limitations of our models. Unlike mating-bias barriers, which establish premating RI through mating trait incompatibilities during encounters, habitat-preference barriers achieve RI by reducing the likelihood of such encounters. Theoretically, distinct niche resources do not need to be spatially separated for the habitat-preference mechanism to operate. The same mechanism can generate premating barriers across multiple dimensions—spatial, temporal, behavioral, or mechanical—as long as distinct habitats exist along these axes [36-39]. Thus, the habitat-preference mechanisms described in this study may be broadly applicable to a wide range of reproductive barriers that function similarly by limiting opportunities for mating encounters.

## VII. FUTURE RESEARCH

While geographical isolation is undoubtedly a major mechanism of speciation, many speciation events in nature likely occurred in the presence of gene flow [50]. In this study, we explored how habitat preference can reduce mating encounters between divergent populations and promote RI. At first glance, habitat-preference isolation may appear to function similarly to geographical isolation. However, a key distinction in our model is that offspring from inter-niche matings can return freely to their adaptive habitats and access resources optimally. By contrast, offspring produced across a geographical barrier are typically confined to suboptimal environments, resulting in reduced fitness. Thus, when divergent ecological niches are separated by geographical barriers, selection against maladaptive immigrants is likely to be a major force driving the evolution of RI.

Properly modeling the effects of geographical isolation likely requires a two-island or continent– island framework [7, 14, 15, 28]. As a direction for future research, our existing models could be extended to simulate two sympatric habitats with movement between them regulated by a migration parameter [7, 14, 51]. This would enable a direct comparison of the dynamics of speciation driven by geographical separation versus those driven by habitat-preference barriers.

In a secondary contact scenario—where two allopatrically diverged ecotypes re-encounter one another and hybridize in a suture zone [29, 44, 52-54]—habitat preference is likely to be the first barrier to evolve to reduce maladaptive hybrid loss. The evolution of additional mating-bias traits in this context may introduce incompatibilities between frontier ecotypes in the suture zone and their larger parental populations in the background, potentially creating a fitness disadvantage for the frontier ecotypes.

However, if hybridizing ecotypes in the suture zone evolve new mating-bias traits that are selectively neutral with respect to the background populations, this incompatibility disadvantage may be avoided. In such cases, the habitat-preference and novel mating-bias traits that arise to minimize hybrid loss within the suture zone could diffuse into the background populations through neutral drift. Nonetheless, ongoing gene flow between individuals in the hybrid zone and the broader parental populations may erode the reproductive isolation developing in the suture zone, potentially making it less effective than what can be achieved in a purely sympatric setting. Future studies could usefully explore these complex and nuanced dynamics associated with secondary contact.

## VIII. ACKNOWLEDGMENTS

We would like to express our grateful appreciation to David Hernandez, Taylor Rios, Lauren Gieck, and James Lin for reviewing the manuscript and offering valuable comments.

## IX. APPENDIX

### Derivation of the Encounter Matrices

In this study, habitat-preference barriers are modeled under two distinct spatial configurations: an open-space system and a no-open-space system (see Methodology for details). This appendix presents the mathematical derivations of the encounter probability matrices (Figs M7 and M9) associated with habitat-preference barriers in each spatial context. The models are constructed to simulate a “perfect match” scenario, in which every individual in the sympatric population successfully encounters a potential mate during a matching round, ensuring that no individuals are left unpaired.

#### Case 1: Habitat-Preference Barriers in an Open-Space System

In habitat-preference models under open-space systems, individuals can encounter one another in one of three distinct regions: habitat *A*, habitat *B*, or a shared communal open space (Fig M8). It is assumed that niche ecotypes do not enter the habitat of the opposite ecotype. Within each ecotype, the presence of habitat-preference mutations causes a proportion of individuals to remain in their native habitat (i.e., home-bound), while the rest are free to leave and explore the communal open space (i.e., free-roaming).

In habitat *A*, individuals genetically predisposed to remain in their adaptive habitat form group *AH* (*A*-home). The remaining individuals, who are not constrained and are free to move into the open space, constitute group *AF* (*A*-free). Similarly, in habitat *B*, individuals that stay within their adaptive habitat form group *BH* (*B*-home), while those that roam freely are classified as group *BF* (*B*-free).

In summary, individuals in groups *AH* and *BH* are restricted to encounters within their respective habitats, while those in groups *AF* and *BF* may interact within their native habitats or in the communal open space. To model encounter behavior, we assume that individuals in *AF* and *BF* visit a given region—habitat *A*, habitat *B*, or the open space—at rates proportional to the number of individuals present in that region relative to the total number of individuals across all accessible regions.

For example, individuals in group *AF* are assumed to encounter others in the open space at a rate defined by:

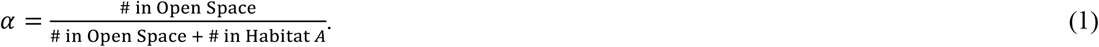

Similarly, the rate at which individuals in group *BF* encounter others in the open space is defined by:

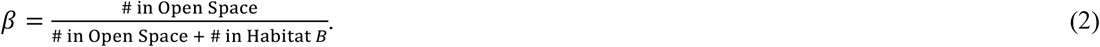

Alternatively, expressed in terms of the defined variables, we obtain the following system of equations that describe the encounter rates in the open space:

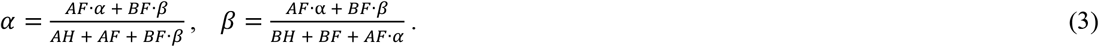

If either *AF* or *BF* equals zero, the solution to equations (3) is trivial (i.e., *α* = *β* = 0), indicating that all individuals remain in their adaptive habitats and no encounters occur in the open space. However, when the solution is non-trivial—i.e., both *AF* and *BF* are nonzero—the system yields the following closed-form solutions:

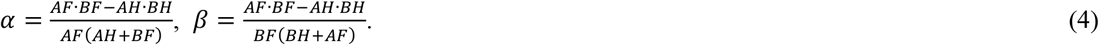

However, as shown in the solutions in (4), nonnegative values of *α* and *β* require that *AF* · *BF* ≥ *AH* · *BH*. Immediately after the invasion of a habitat-preference mutation, this condition is satisfied, but over time, the product *AH* · *BH* gradually increases as more individuals become home-bound, while *AF* · *BF*—representing free-roaming individuals—steadily declines. When *AF* · *BF* = *AH* · *BH*, both *α* and *β* reach zero, indicating that all individuals remain in their adaptive habitats and no encounters occur in the open space. Since *α* and *β* cannot take negative values, any scenario in which *AF* · *BF* < *AH* · *BH* yields only the trivial solution (*α* = *β* = 0). In this case, complete habitat isolation is achieved: the open space remains unoccupied, and all encounters occur exclusively within each ecotype’s adaptive habitat.

When the system solution is non-trivial, we define 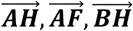 and 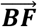 as vectors representing the genotype distributions within their respective groups. From these, we construct a normalized population vector: 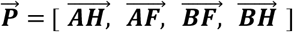 where the sum of all elements in 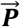 equals 1. This normalized population vector forms the basis for constructing the encounter probability matrix shown in Fig M7.

Based on these definitions, the probability submatrix for encounters occurring in the habitat *A* region (i.e., between individuals from 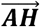 and 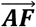) is given by:

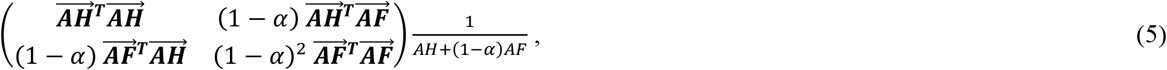

Here, the normalization constant, 1/(*AH* + (1 − *α*)*AF*), ensures that the values in the submatrix sum to *AH* + (1 − α)*AF*, which represents the total population within habitat *A*.

Similarly, the encounter probability submatrix for interactions within the habitat B region (i.e., between individuals in 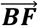 and 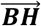) is given by:

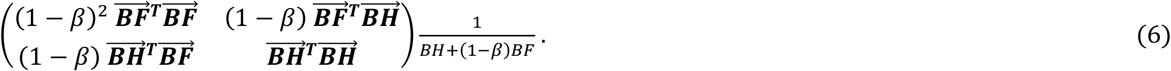

Lastly, the encounter probability submatrix for interactions in the open-space region (i.e., between individuals in 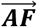 and 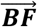) is given by:

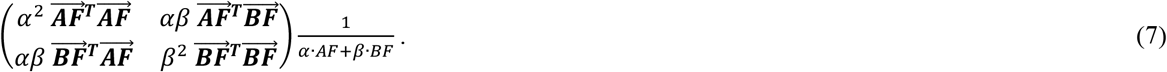

Adding the three submatrices from (5)–(7) and simplifying yields the full encounter probability matrix for the entire population:

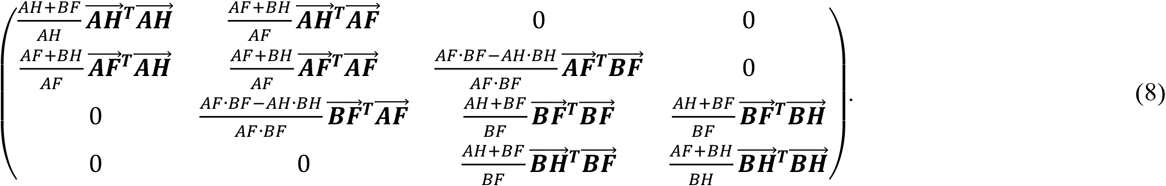

All entries in this matrix sum to 1, consistent with the normalization of the population vector.

#### Case 2: Habitat-Preference Barriers in a No-Open-Space System

In habitat-preference models under no-open-space systems, niche ecotypes are allowed to enter each other’s habitats, and all encounters occur in either habitat *A* or habitat *B*. Within each ecotype, the presence of habitat-preference mutations causes a proportion of individuals to remain in their adaptive habitat (i.e., home-bound), while the remainder are free to leave and visit the other habitat (i.e., free-roaming).

For habitat *A*, all niche-*A* ecotypes that carry the habitat-preference mutation and are required to stay in their adaptive habitat are placed into group *AH*, based on the habitat-preference bias *θ*, which defines the proportion of individuals with the mutation that remain home-bound. The remaining niche-*A* individuals, who are free to move between habitats, are classified as group *AF*. Individuals from habitat *B* are grouped in the same way: those constrained to stay in their adaptive habitat form group *BH*, while those that can move freely are assigned to group *BF*. Finally, all free-roaming individuals from both ecotypes—those in groups *AF* and *BF*—are combined into a single group, denoted *FF*, representing all individuals capable of visiting either habitat.

As a result, individuals in groups *AH* and *BH* are restricted to meet within their respective adaptive habitats, while individuals in *FF* may encounter and mate in either habitat. Assuming that individuals in *FF* visit habitats at rates proportional to the number of individuals present in each habitat, we define a probability ratio *μ*, which represents the likelihood that a member of *FF* will visit and meet in habitat *A*:

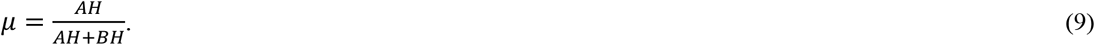

Accordingly, 1 − *μ* represents the likelihood that a member of *FF* will visit and meet in habitat *B*. In the special case where *AH* + *BH* = 0, we define *μ* = 0.5. While any value in the interval [0,1] would be acceptable in this scenario, setting *μ* = 0.5 avoids division by zero in subsequent calculations.

We define 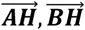 and 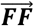 as vectors representing the genotype distributions for their respective groups. Here 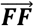 is composed of all free-roaming individuals and is defined as: 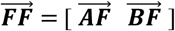. The normalized population vector is then: 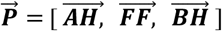. Note that the sum of all elements in 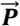 equals 1. Based on these definitions, the probability submatrix for encounters occurring in habitat *A* is given by:

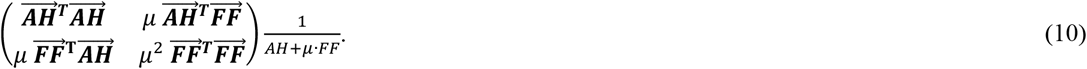

The normalization constant, 1/(*AH* + *μ* · *FF*), ensures that the values in the habitat *A* submatrix sum to *AH* + *μ* · *FF*, which corresponds to the total population within habitat A.

Likewise, the encounter probability submatrix for interactions in habitat *B* is given by:

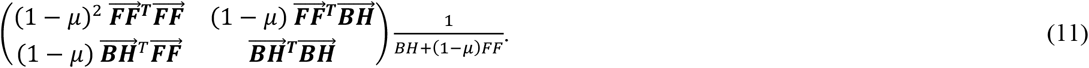

Define the following constants:

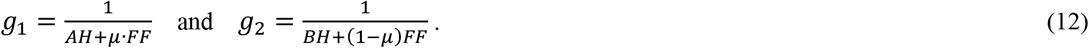

Combining submatrices (10) and (11) yields the full encounter probability matrix for the entire population:

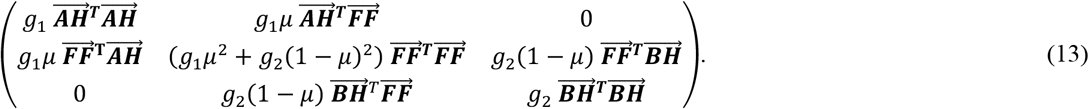

All elements in the matrix sum to 1, consistent with the normalization of the total population.

